# An Amygdalar-Vagal-Glandular Circuit Controls the Intestinal Microbiome

**DOI:** 10.1101/2024.06.02.594027

**Authors:** Hao Chang, Matthew H Perkins, Leonardo S Novaes, Feng Qian, Wenfei Han, Ivan E de Araujo

## Abstract

Psychological states can regulate intestinal mucosal immunity by altering the gut microbiome. However, the link between the brain and microbiome composition remains elusive. We show that Brunner’s glands in the duodenal submucosa couple brain activity to intestinal bacterial homeostasis. Brunner’s glands mediated the enrichment of gut probiotic species in response to stimulation of abdominal vagal fibers. Cell-specific ablation of the glands triggered transmissible dysbiosis associated with an immunodeficiency syndrome that led to mortality upon gut infection with pathogens. The syndrome could be largely prevented by oral or intra-intestinal administration of probiotics. In the forebrain, we identified a vagally-mediated, polysynaptic circuit connecting the glands of Brunner to the central nucleus of the amygdala. Intra-vital imaging revealed *that* excitation of central amygdala neurons activated Brunner’s glands and promoted the growth of probiotic populations. Our findings unveil a vagal-glandular neuroimmune circuitry that may be targeted for the modulation of the gut microbiome. The glands of Brunner may be the critical cells that regulate the levels of *Lactobacilli* species in the intestine.

## Introduction

Gut microbiomes, *i*.*e*., the microbial communities of the intestinal mucosa, co-evolved with their hosts to promote nutrient digestion and protection against food-borne pathogens^1–4^. Growing evidence indicates that psychological states impact on systemic immunity by altering the host’s bacterial microbiome^3,5–7^, pointing to a causal link between brain activity and gut bacterial homeostasis. Indeed, numerous human subject and preclinical studies report associations between psychological stress and dysbiosis^8–11^. In non-human primates, stress models such as maternal separation induce significant decreases in probiotic *Lactobacilli* levels concomitantly to increasing vulnerability to opportunistic infections^12^. Consistently, rodent studies show that probiotic administration may improve emotional and physiological markers in models of anxiety^13^.

Such influence of brain states on the gut microbiome may occur via changes in mucosa-bacterial interactions^7,14–17^. Thus, in children, psychosocial stress was associated with deficient mucosal immune protection, reflecting both lower concentrations of secretory immunoglobulin A and increased occurrences of opportunistic infections^18^. Conversely, the opposite effects are observed upon induction of psychological relaxation^19^. Indeed, it is presumed that stress-induced suppression of *Lactobacilli* levels in primates was driven by an inhibition of gut mucous secretion^12^. Unfortunately, it remains unknown how brain states may control mucosal secretion in such a way to impact on the microbiome.

Our main objective was to identify neuronal pathways that enable the brain to impact on the mucosal-microbiome system. The glands of Brunner, an exocrine structure confined to the duodenal submucosa, primarily consist of mucus-producing cells, and have been traditionally believed to function as buffers against gastric acidity^20–22^. Critically, the glands of Brunner are distinctively targeted by nerve terminals, with neural stimulation being necessary for mucus secretion^20,23,24^. Of note, a significant proportion of these terminals are of vagal origin^23^. We thus hypothesized that the vagal innervation of the mucosal glands of Brunner mediates the influence of brain activity on the microbiome. Accordingly, we probed this neural-glandular circuit using cell-specific approaches for single nuclei sequencing, ablation, intra-vital imaging, electrophysiological, and behavioral studies.

## Results

### Nutrient-sensing gut hormones activate Brunner’s glands via the vagus nerve

Brunner’s glands (“BG”) are located beneath the epithelium, within the upper, ampullary duodenal submucosa (Figure S1A). We first aimed to establish the influence of the vagus nerve on BG activity *in vivo*. BG is known to respond to the pro-digestive, vagus-mediated gut hormone cholecystokinin (“CCK”)^23,24^. We confirmed that intraperitoneal CCK administration acts on BG to induce both mucus secretion and vesicle trafficking to the apical membrane (Figures S1B-F). Using this model, we proceeded to evaluate the behavior of the glands *in vivo*. Since GLP-1 receptors are molecular markers of BG^20^, we generated Glp1r-ires- Cre×Ai148(TIT2L-GC6f-ICL-tTA2)-D[GCamp6]×Ai9 (“Glp1r[GCamp6]”) mice to cell-specifically express the fluorescent calcium-indicator GCamp6 in BG (Figures S1G-J). We then implanted Glp1r[GCamp6] mice with an abdominal glass window to perform intra-vital imaging of intracellular calcium activity in BG (Figure 1A and Supplemental Movie 1). CCK administration induced robust calcium transients throughout BG (Figures 1B-E and Supplemental Movie 2), with calcium currents following a proximal-to-distal spatiotemporal pattern (Figure S1K). To assess the role of the vagus nerve in this phenomenon, we performed these same experiments in Glp1r[GCamp6] mice sustaining either subdiaphragmatic vagotomies or gut-specific sensory vagal denervation. Interrupting vagal transmission in either way completely abolished BG responses to CCK (Figures 1F-I and S1L-P). Thus, abdominal vagal fibers are required for gut hormonal recruitment of BG *in vivo*.

**Figure 1.**
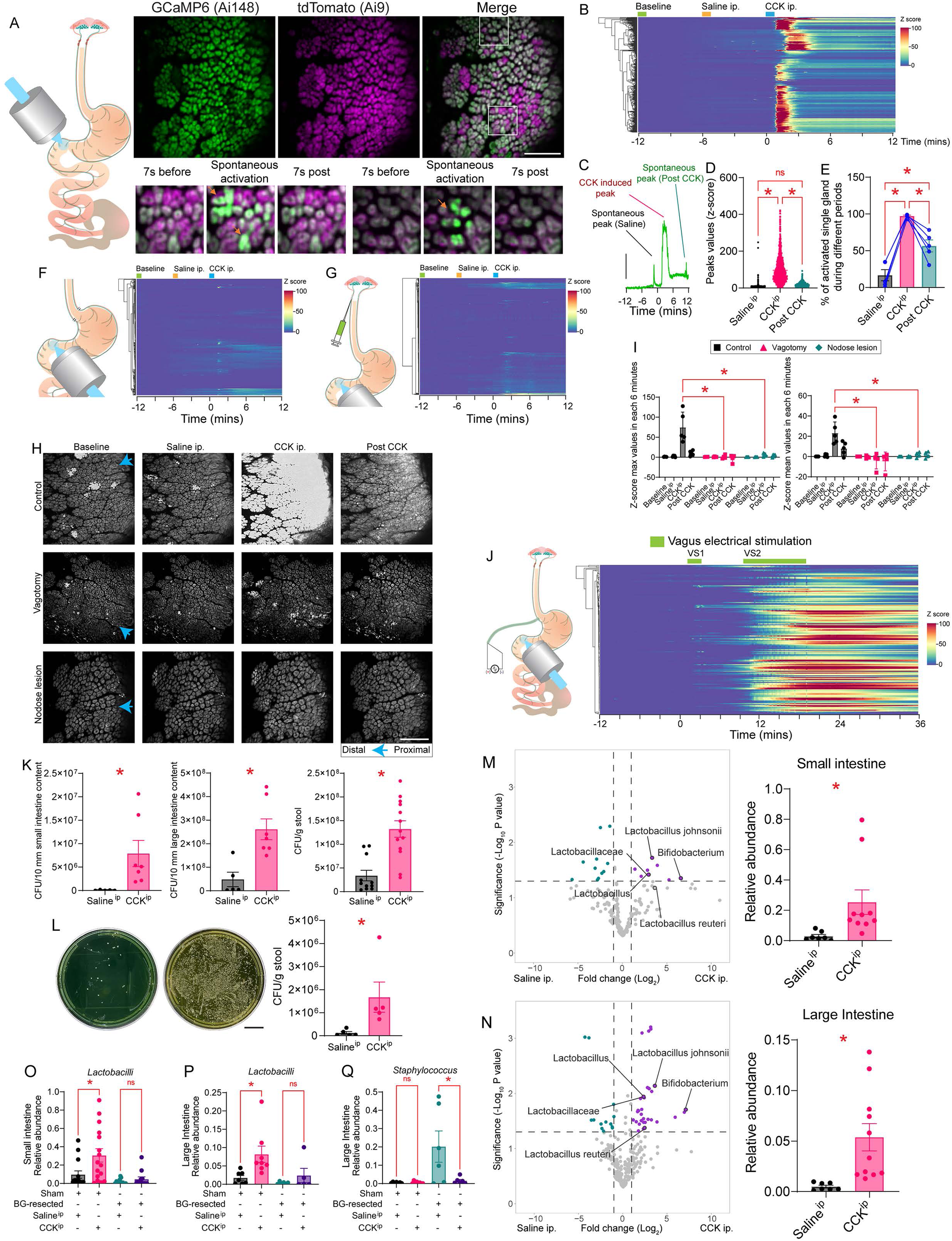
Vagal signals control the microbiome via the glands of Brunner. **A.** Left: Schematics of BG intravital imaging. Right: Using the triple reporter Glp1r-ires- Cre×Ai148(TIT2L-GC6f-ICL-tTA2)-D[GCamp6]×Ai9 (Glp1r[GCamp6]) mice, we detected in BG GCamP6 transients but an unchanging tdTomato (red channel) signal, ruling out movement artifacts. Bar=200μm. **B.** BG exhibited strong calcium transients after intraperitoneal CCK (“CCK^ip^”, 10μg/Kg) stimulation. Heatmap of Z-scores *vs.* baseline period after CCK^ip^, T=-6min: Saline^ip^, T=0: CCK^ip^; Glands from 5 mice; hierarchical/agglomerative clustering was employed normalize and rearrange individual glands. **C.** An example of calcium transient signals from a single gland shows spontaneous peaks before and after CCK^ip^, separated by a conspicuous CCK-induced peak. **D.** Maximum Z-score values for spontaneous peaks before CCK^ip^, CCK^ip^- induced peaks, and spontaneous peaks after CCK^ip^. One-way ANOVA, p<0.0001. Bonferroni post hoc, ∗p<0.0001. **E.** Percentage of activated single glands across animals during a 6-min interval for the Saline^ip^, CCK^ip^, and post-CCK periods. One-way ANOVA, p<0.0001. Bonferroni post hoc, ∗p<0.05. **F-G.** Vagotomy abolished BG responses to CCK. **F:** Left: BG calcium transients in Glp1r[GCamp6]mice sustaining subdiaphragmatic vagotomies. Right: As in panel ***E*,** but subdiaphragmatic vagotomy cases. **G:** Vagal sensory ablation abolished BG responses to CCK. Left: BG calcium transients in Glp1r[GCamp6]mice after CCK-SAP. Right: As in panels ***E*** and ***F***. **H.** Maximum intensity projections of calcium transients in intact, vagotomized, and nodose-ablated Glp1r[GCamp6]mice. Analyses performed for pre-infusion baseline, Saline^ip^, CCK^ip^, and post-CCK^ip^ conditions, each defined as a 6 min-long timelapse stack. Robust calcium influxes were observed after CCK^ip^ in intact mice only. Bar=200μm. **I.** Across-animals group analyses of intravital imaging Z-scores. Left: Z-score maximal values from the four conditions in the three groups of mice. Two-way mixed RM-ANOVA, p<0.001. Bonferroni post- hoc: Intact mice *vs*. vagal lesions in CCK^ip^, both ∗p<0.0001; Vagotomy vs. CCK-SAP, p>0.9999. Right: as before, but for Z-score mean values, p<0.001. Bonferroni post-hoc: Intact mice *vs*. vagal lesions in CCK^ip^, both ∗p<0.0001; Vagotomy vs. CCK-SAP, p=0.4121. **J.** As in panel ***E***. Electrical stimulation of the vagus nerve (∼1.8mA) induces robust calcium transients in BG during intravital imaging. VS1=Vagal stimulation period 1 (short); VS2=Vagal stimulation period 2 (long). **K.** CCK^ip^ enhances intestinal and fecal levels of *Lactobacilli* as measured by culturing samples in MRS plates. Left: Small intestine; Middle: Large intestine; Right: Feces, 2-sample t- tests CCK *vs.* vehicle, all, ∗p<0.05. **L.** Wild-type mice were treated with vehicle and CCK^ip^ for 3 days and then fed a dose of the chloramphenicol-resistant strain *Lactobacillus rhamnosus*. CCK^ip^ promoted *L. rhamnosus* proliferation as detected in stool. Left, example clones cultured with chloramphenicol-containing media; Right: Total clone counts, 2-sample t-test, ∗p=0.0432. Bar=20mm. **M.** Left: Volcano plot contrasting relative abundances of bacterial contents in small intestine using 16S sequencing after CCK^ip^ *vs*. vehicle injections. Cyan represents downregulated, and purple upregulated, bacterial species. Right: Relative abundances of total *Lactobacilli* count after CCK^ip^ *vs*. vehicle injections, 2-sample t-test, ∗p=0.0382. **N.** Same as ***M*** but for large intestine, ∗p=0.0119. **O-Q.** Effects of BG resection on the microbiome. **O.** Relative abundances of total *Lactobacilli*count between sham+saline^ip^, sham+CCK^ip^, BG-resected+saline^ip^, and BG-resected+CCK^ip^ from small intestinal samples. One-way ANOVA, p=0.0003. Bonferroni post hoc, Sham+saline^ip^ vs. Sham+CCK^ip^, ∗p=0.0172. BG- resected+saline^ip^ vs. BG-resected+CCK^ip^, p>0.9999. **P.** Same as ***O*** but for large intestinal contents. p=0.0071. Bonferroni post hoc, Sham+saline^ip^ vs. Sham+CCK^ip^, ∗p=0.0372; BG- resected+saline^ip^ vs. BG-resected+CCK^ip^, p>0.9999. **Q.** Same as ***P*** but for *Staphylococci* instead of *Lactobacilli* for large intestinal contents, p=0.0032. Bonferroni post hoc, Sham+saline^ip^ vs. Sham+CCK^ip^, p>0.9999; BG-resected+saline^ip^ *vs*. BG-resected+CCK^ip^, ∗p=0.0187.

### Electrical stimulation of the vagus nerve activates Brunner’s glands in vivo

To support the findings above, we implanted a cuff electrode on the vagal trunk to deliver electrical pulses to the nerve in Glp1r[GCamp6] mice. We concomitantly performed intra-vital imaging of BG. We found that vagal stimulation was sufficient to induce robust calcium transients across BG, with sensitivity for gland mobilization markedly increasing with the duration of the pulses (Figure 1J). Thus, vagal fiber activity is both necessary and sufficient for BG activation.

### Cholecystokinin promotes probiotic bacterial proliferation via Brunner’s glands

Based on the above, we then hypothesized that BG may link nerve activity to microbiome composition. Given the ability of CCK to engage BG secretion, we first evaluated whether the gut hormone can alter the microbiome. Unexpectedly, administration of CCK promoted the proliferation of *Lactobacilli* species in both small and large intestinal tissues, an effect also detected in fecal samples (Figure 1K). These findings were obtained and independently replicated by measuring the culturing intestinal and fecal samples in *Lactobacillus*-specific media; resulting clones were confirmed to be *Lactobacilli* using supplemental sequencing (Figures S1Q-S). Consistently, daily CCK treatment induced marked proliferation of *Lactobacillus rhamnosus* after seeding the gut of mice with this strain (Figure 1L). Unbiased volcano plot analyses of 16S sequencing of small and large intestinal samples revealed that CCK promoted the proliferation of different *Lactobacilli* species, in particular *L. johnsonii* (Figures 1M-N). To investigate if these effects were mediated by BG, we designed a surgical approach to resect BG from the submucosa that surrounds the duodenal ampulla while preserving luminal and myenteric tissues (see STAR Methods and Supplemental Table 1 for details). CCK-induced probiotic population growth was entirely abolished after BG resection (Figures 1O-P and S1T-AA). Importantly, BG ablation induced a dysbiosis-like increase in the relative abundance of potentially pathogenic *Staphylococci* species in both small and large intestines (Figures 1Q and S1BB). Additional details of sequencing results are shown in Figure S1CC.

In sum, while the ability of the vagal-dependent hormone CCK to modulate the microbiome requires intact glands of Brunner, resection of the glands leads to abnormal growth of potentially pathogenic bacteria.

### Vagal efferents are synaptically connected to Brunner’s glands, which express pro-secretory cholinergic receptors at the single-cell level

If it is indeed the case that vagal parasympathetic fibers signal directly onto BG, these glands must be in contact with synaptic endings of vagal origin. Also, they must additionally express receptors known to enable cholinergic action on exocrine cells. To verify the first of these criteria, we first injected the *Cre*-dependent construct AAV9-hSyn-DIO-GFP into the dorsal motor nucleus of vagus (“DMV”) of Chat-IRES-Cre mice, resulting in the labelling of vagal efferent fibers arising from DMV. Dense efferent innervation was detected on BG, but not on epithelial goblet mucous cells across the intestine (Figures 2A-B). This suggests that vagal influence on mucus secretion is specific to BG, and does involve epithelial mucous cells. Moreover, to detect the presence of vagal synaptic endings on BG, we injected the *Cre*-dependent construct AAV9- hSyn-FLEx-mGFP-2A-Synaptophysin-mRuby into DMV of Chat-IRES-Cre mice. Most vagal synaptic endings were clearly detected on BG, and virtually none on the overlying villi (Figure 2C).

**Figure 2.**
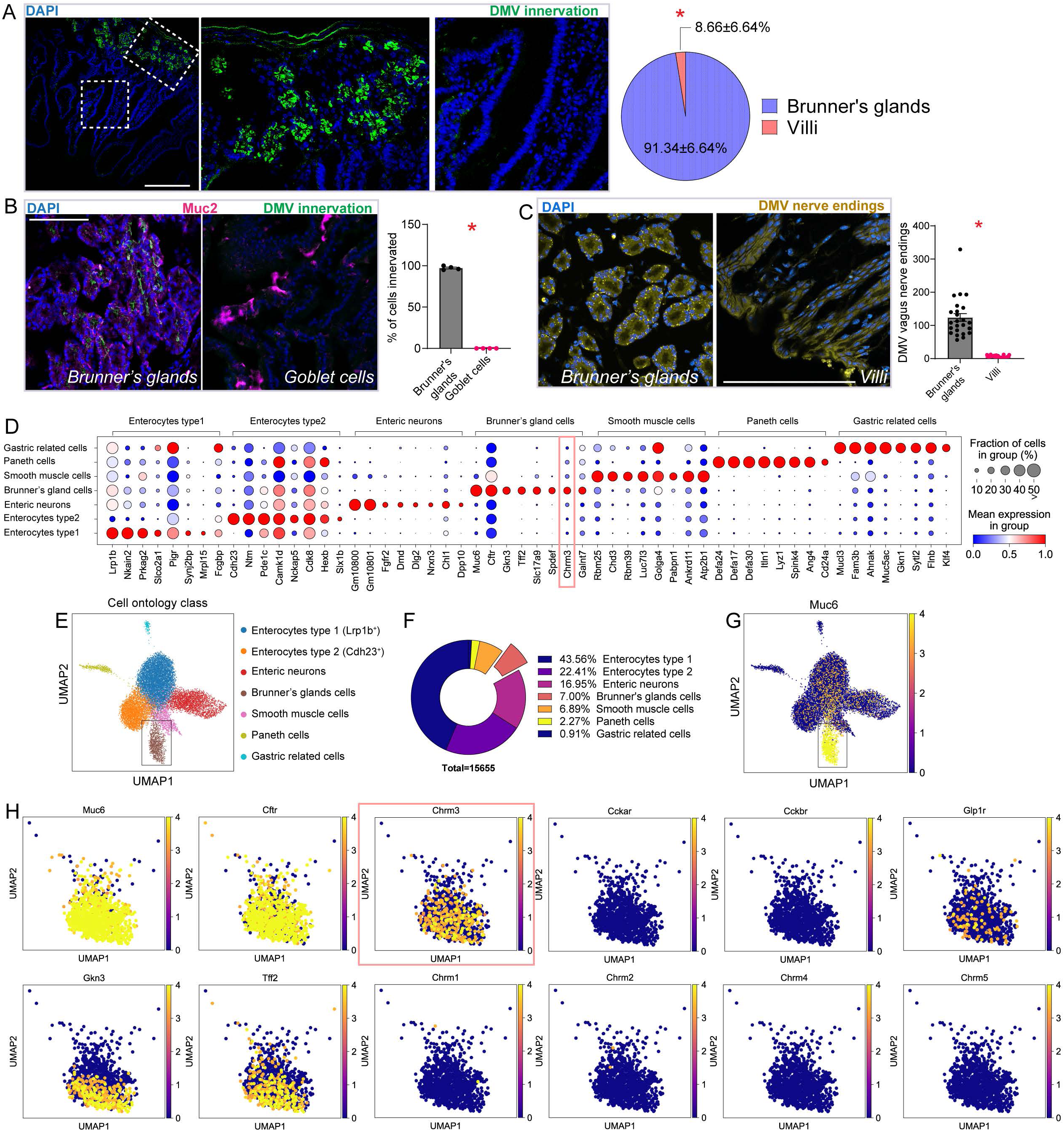
Vagal efferents innervate Brunner’s glands, which express pro-secretory cholinergic receptors at the single-cell level. **A.** Left: The AAV9-hSyn-DIO-GFP construct was injected into dorsal motor nucleus of vagus (DMV) of Chat-ires-Cre mice to label vagal efferents. Center and Right: Amplified view of selected regions. Bar=100μm. Pie plot: Percentage of BG *vs.* villus innervation, paired t-test, ∗p<0.0001. **B.** As in panel *A*. Panel shows confocal sections stained using anti-Muc2 antibody. Strong labeling of vagal fibers was visualized in BG but not near goblet (Muc2+) cells; percentage of innervated BG *vs.* goblet cells, paired t-test, ∗p<0.0001. Bar=100μm. **C.** Left: AAV9-hSyn-FLEx-mGFP-2A-Synaptophysin-mRuby was injected into DMV of Chat-ires-Cre mice to label efferent vagal synaptic endings. Confocal imaging revealed that synaptic endings were localized on Brunner’s glands but not on the overlying villi. Right: DMV synaptic endings on glands *vs.* villi, 2-sample t-test, ∗p<0.0001. Data pooled from 5 mice. Bar=100µm. **D.** Top (eight) most enriched marker genes for the seven major cell types, namely, Enterocytes type I (Lrp1b^+^), Enterocytes type II (Cdh23^+^), enteric neurons, Brunner’s glands cells, smooth muscle cells, Paneth cells, and gastric-related cells. Note the cholinergic secretory muscarinic receptor 3, *Chrm3*, among the topmost enriched genes in Brunner’s glands cells. **E.** Cell ontology class of duodenal tissue containing Brunner’s glands, as defined by single nuclei sequencing. **F.** Percentage of the total number of validated cells for each cell type. In proximal duodenum, 7% of cells were identified as Brunner’s glands cells. G. *Muc6* is the major marker used to annotate Brunner’s glands cells. **H.** Previously reported marker genes for Brunner’s glands cells: *Muc6*, *Cftr*, *Gkn3* and *Tff2* and co-expression with different receptor types present in Brunner’s glands cells. Note the presence of *Chrm3* (red border) and absence of CCK receptor *Ccka* and *Cckb* transcripts.

To verify the criterion of cholinergic receptor expression in BG, we performed single nuclei sequencing of RNA transcripts from proximal duodenal tissue. Our specific question was whether BG cells co-express at the single cell level the muscarinic receptor M3 – the cholinergic receptor that is critical for parasympathetic control of exocrine secretion^25^. We found that cells expressing the mucin gene *Muc6*, distinctive of BG among duodenal cells^21^, constituted the only cell type clearly co-expressing the M3 receptor gene. Overall, the results above demonstrate that mucous BG cells express the machinery required for vagus-induced secretion. Consistently, we found that other genes equally associated with cellular mucus production, such as *Galnt7* and *Tff2*, co-express at the single-cell level with both *Muc6* and the M3 receptor gene. Finally, but equally important, mucous BG cells failed to express any CCK receptors transcripts, reinforcing the notion that the action of CCK on BG is entirely mediated by abdominal vagal fibers. Details on the single nuclei sequencing analyses are shown in Figures 2D-H and S2.

### Cell-specific ablation of the glands of Brunner leads to spleen abnormalities, mortality upon pathogen infection, and exaggerated intestinal permeability

Based on the observation that BG secretory activity modulates the microbiome, we predicted that ablation of the glands would lead to immunological and intestinal barrier dysfunctions. We designed a strategy to cell-specifically ablate BG. We generated triple mutant Glp1r[GCamp6]×ROSA26iDTR mice to express the diphtheria toxin receptor exclusively in BG cells. To evaluate the efficacy of this approach, we compared the effects of injecting diphtheria toxin (DTx) into the duodenal submucosa of Glp1r[GCamp6]×ROSA26iDTR mice with those produced by the surgical resection method in wild-type mice. Overall, the surgical and DTx- induced ablations produced nearly indistinguishable effects. Importantly, both approaches eliminated most BG cells while preserving duodenal and pancreatic GLP1R+ cells (Figures 3A- D and S3A-E).

**Figure 3.**
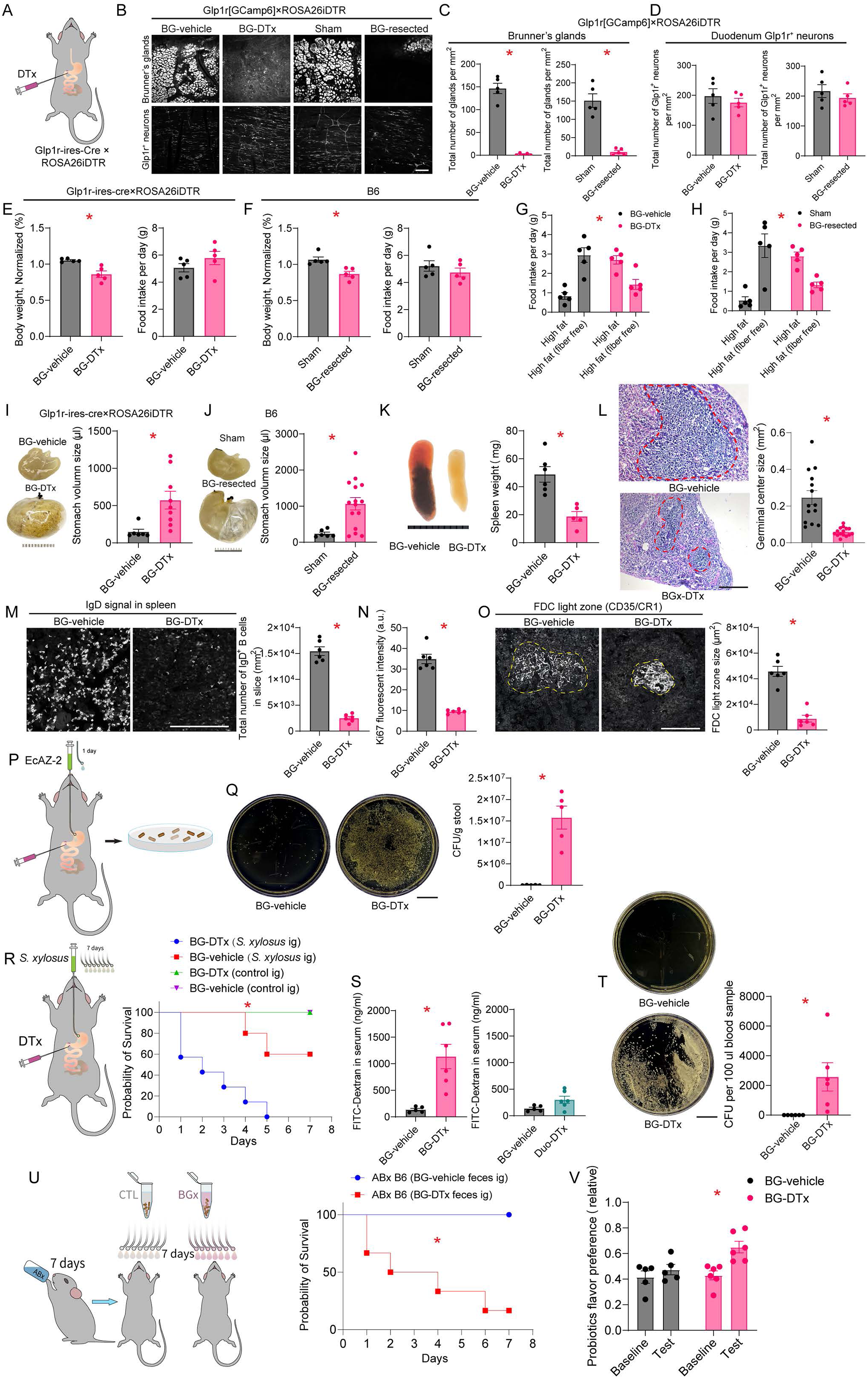
Ablation of the glands of the Brunner leads to an immunodeficiency syndrome and to mortality upon intestinal infection. **A.** Strategy for cell-specific ablation of BG using diphtheria toxin (DTx) injections into the upper duodenal submucosa of Glp1r-ires-Cre×ROSA26iDTR mice. **B.** Using Glp1r[GCamp6]×ROSA26iDTR triple transgenic mice enabled visualizing ablation efficiency. Confocal imaging reveals nearly total ablation of BG in DTx-injected (“BG-DTx”) mice, while other duodenal Glp1r^+^ enteric cells were preserved. Equivalent results were obtained with the resection approach (“BG-resected”) mice. Bar=300µm. **C.** Left: Total number of glands/mm^2^ in proximal duodenum in BG-vehicle *vs.* BG-DTx mice, 2-sample t-test, ∗p<0.0001. Right: Total number of glands/mm^2^ in proximal duodenum in sham *vs.* BG-resected Glp1r[GCamp6]×ROSA26iDTR mice, 2-sample t-test, ∗p<0.0001. Bar=300µm. **D.** Equivalent analyses for Glp1r^+^ enteric neurons, Left: p=0.4677 (BG-vehicle *vs.* BG-DTx); Right: p=0.3979 (sham *vs.* BG-resected). **E.** Normalized body weight (Left) and daily food intake (in gram, Right) of BG-DTx *vs.* BG-vehicle mice. Ablated mice showed decreased body weights independently of food intake. Normalized body weight: 2-sample t-test, ∗p=0.0032; Daily food intake: p=0.2439. **F.** Same as *E* but for sham *vs.* BG-resected wild-type (B6) mice. Normalized body weight: ∗p=0.0048. Daily food intake: p=0.3715. **G.** BG-vehicle and BG-DTx mice displayed significantly differences in food preferences, with BG-DTx showing higher preferences for fiber-rich fatty pellets, 2-way mixed RM-ANOVA, Surgery x food type, ∗p=0.0001. **H.** Same as *G*, with similar effects found for sham *vs.* BG-resected mice, ∗p=0.001. **I.** Significant stomach distension in BG- DTx *vs.* BG-vehicle mice, 2-sample t-test, ∗p=0.0146. Bar=1cm. **J.** Same as *I*, similar effects for sham *vs.* BG-resected mice, ∗p=0.0077. Bar=1 cm. **K-O.** BG ablation alters spleen characteristics. **K:** Significantly smaller spleen sizes in BG-DTx *vs.* BG-vehicle mice, Left: representative samples from BG-DTx *vs.* BG-vehicle mice, Right: spleen weights, 2-sample t- test, ∗p=0.0018. Bar=1cm. **L:** Smaller spleen germinal center sizes in BG-DTx *vs.* BG-vehicle mice, ∗p<0.0001. Bar=200µm. **M:** Marked suppression of B cell population in spleen of BG-DTx mice. Left: IgD immunostaining in spleen sections from BG-DTx *vs.* BG-vehicle mice. Right: total number of IgD^+^ B cells/mm^2^ of spleen sections, 2-sample t test, ∗p<0.0001. Bar=100 µm. **N.** Suppression of cell proliferation markers in spleen of BG-DTx mice. Average Ki67 fluorescent intensity in BG-vehicle *vs.* BG-DTx mice, 2-sample t test, ∗p<0.0001. **O.** Suppression of follicular dendritic cell (FDC) markers in spleen of BG-DTx mice. FDC light zone size in spleens of BG-DTx *vs.* BG-vehicle mice. Left: Immunostaining against CD35/CR1. Right: FDC light zone sizes, 2-sample t-test, ∗p<0.0001. Bar=100 µm. **P.** Gut colonization with the kanamycin- resistant *E. coli* strain EcAZ-2 (10^10^ CFUs) in BG-vehicle *vs.* BG-DTx mice. **Q.** Enhanced EcAZ- 2 proliferation in BG-DTx mice. Left: Representative examples of 1mg fecal samples plated on kanamycin-LB agar. Right: Fecal EcAZ-2 counts were significantly greater in BG-DTx *vs.* BG- vehicle mice, 2-sample t-test, ∗p<0.0001. Bar=2cm. **R.** Survival curves following gut colonization with pathogen *Staphylococcus xylosus*. After seeding gut with *S. xylosus* (10^8^ CFUs) via oral gavages, BG-DTx mice sustained significant death rates whereas all BG-vehicle mice survived the infection, Log Rank (Mantel-Cox), ∗p<0.0001. **S.** “Leaky guts” in BG-DTx mice. Intestinal permeability was assessed by measuring FITC-dextran in systemic circulation 3hs after intraluminal administration of 4kDa FITC-dextran. Left, BG-vehicle *vs*. BG-DTx: 2-sample t-test, ∗p<0.0035. Right: control experiments measuring gut permeability in Glp1r-ires- cre×ROSA26iDTR mice injected with DTx in myenteric layer of duodenum (∼1.5mm underneath BG, Duo-DTx). BG-vehicle *vs* Duo-DTx: p=0.0646. **T.** Translocation of pathogenic bacteria into systemic circulation in BG-DTx mice. Left: blood samples were cultured on BHI agar plates after two-days of gavaged *S. xylosus*. Right: *S. xylosus* invasion of blood circulation in BG-DTx, not BG-vehicle, mice, 2-sample t-test, ∗p=0.0221. Bar=2cm. **U.** Left: Mortality due to transmissible BG-DTx dysbiosis. Transplantation of fecal samples from BG-DTx and BG-vehicle mice into C57BL/6J SPF mice following 7 days of treatment with antibiotic cocktail (“Abx”). Fecal sample solutions were gavaged once/day for another seven days. Survival analyses show that B6 SPF mice treated with BG-DTx fecal samples sustained significant death rates, Log Rank (Mantel- Cox) ∗p<0.001. All BG-vehicle mice survived the treatment. **V.** Greater preferences for probiotic solutions in BG-DTx mice, two-way mixed RM-ANOVA, ablation effect, ∗p=0.0016.

Both ablation approaches produced a mild but significant drop in body weight independently of changes in food intake (Figures 3E-F). Interestingly, both ablation procedures induced higher preferences for fiber-rich *vs.* fiber-free fatty food pellets (Figures 3G-H and S3F-K). Finally, both approaches induced gastric bloating. The appropriate control groups for each approach were phenotypically indistinguishable (Figures 3I-J and S3L-O).

We then proceeded to investigate in greater depth the effects of cell-specifically eliminating BG using diphtheria toxin injections into the duodenal submucosa of Glp1r-ires-Cre×ROSA26iDTR mice (“BG-DTx” mice). Control animals of this strain were injected with vehicle (“BG-vehicle”). We first detected marked abnormalities in spleen morphology in BG-DTx mice, including reductions in volume, germinal center area, follicular dendritic cell counts, IgD^+^ B-cell counts, and cell proliferation markers (Figures 3K-O and S3P).

We hypothesized that these changes in spleen characteristics were indicative of a suppressed immune function. We thus seeded the gut of both BG-DTx and BG-vehicle control mice with the *E. coli* strain EcAZ-2 via oral gavage and detected increases in EcAZ-2 counts in the excrements of BG-DTx mice (Figures 3P-Q). We then performed the same experiment using the pathogenic *Staphylococcus xylosus* strain, an opportunistic pathogen^26^. While the infection led to marked mortality in BG-DTx mice, all BG-vehicle control mice survived the contamination (Figure 3R).

Finally, we observed greater intestinal permeability in BG-DTx mice as assessed by measuring FITC-dextran in systemic circulation after intraluminal administration (Figures 3S and S3Q). Consistently, after gut colonization, high levels of *Staphylococcus xylosus* were detected in blood of BG-DTx, while remaining undetected in control mice (Figure 3T). Thus, overall, ablation of the glands of Brunner leads to a markedly weakened ability to fight intestinal infections and to a physiological state akin to the “leaky gut” syndrome^27^.

### Fecal transplants from animals lacking Brunner’s glands increase mortality

We then assessed the physiological consequences of dysbiosis in BG-DTx mice. Our microbiome sequence analyses had shown that *Staphylococcus xylosus* is strongly and spontaneously enriched in the gut of BG-resected mice (see Figures S1V-W). To confirm that dysbiosis can lead to vulnerability to intestinal infections in BG-DTx mice, we collected feces from both BG-vehicle and BG-DTx mice and prepared two liquid dilutions that were administered via gavage to two groups of wild-type mice previously treated with antibiotics for seven days. We found that fecal transplants from BG-DTx mice robustly induced significant mortality, whereas all animals receiving fecal transplants from BG-vehicle mice survived the experiment (Figures 3U and S3R). Confirming that the BG-ablation induced dysbiosis affected the animals’ physiology, behavioral tests revealed a striking increased preference for solutions containing live probiotics in BG-DTx, but not in control, mice (Figure 3V). Of note, these effects were independent of any changes in duodenal morphology or motility (Figures S3S-T). Increased probiotic preference is possibly analogous to the greater ingestion of fiber-containing foods after BG ablation (Figures 3G-H); indeed, both outcomes suggest a motivation to behaviorally compensate for dysbiosis. The findings also show that BG-ablation induces a transmissible dysbiotic syndrome associated with immunodeficiency.

### Brunner’s glands contribute to changes in the microbiome induced by bariatric surgery

It is known that bariatric surgery produces beneficial effects on the gut microbiota composition that reflect obesity remission^28^. We therefore investigated the impact of preserving BG in the alimentary tract after restrictive bariatric surgery. We recorded physiological and behavioral parameters in three groups of wild-type mice. In addition to a sham-operated group, one second group sustained a bypass of the upper intestine that preserved BG in the alimentary tract (“BG+” mice). This was achieved by performing a duodenal ligation proximal to the duct of Oddi, such that the duodenal ampulla and the distal jejunum were sutured side-to-side (Figures 4A-C; note that BG is located within the ampullar submucosa as in Fig. 1A). A third group sustained a bypass of the upper intestine that excluded BG from the alimentary tract (“BG-”) via a ligation just distal to the pylorus, such that the pylorus and distal jejunum were sutured end-to-side (Figures 4D-E). Note that neither surgery involved ablating BG. See *STAR Methods* and Supplemental Table 1 for further details.

**Figure 4.**
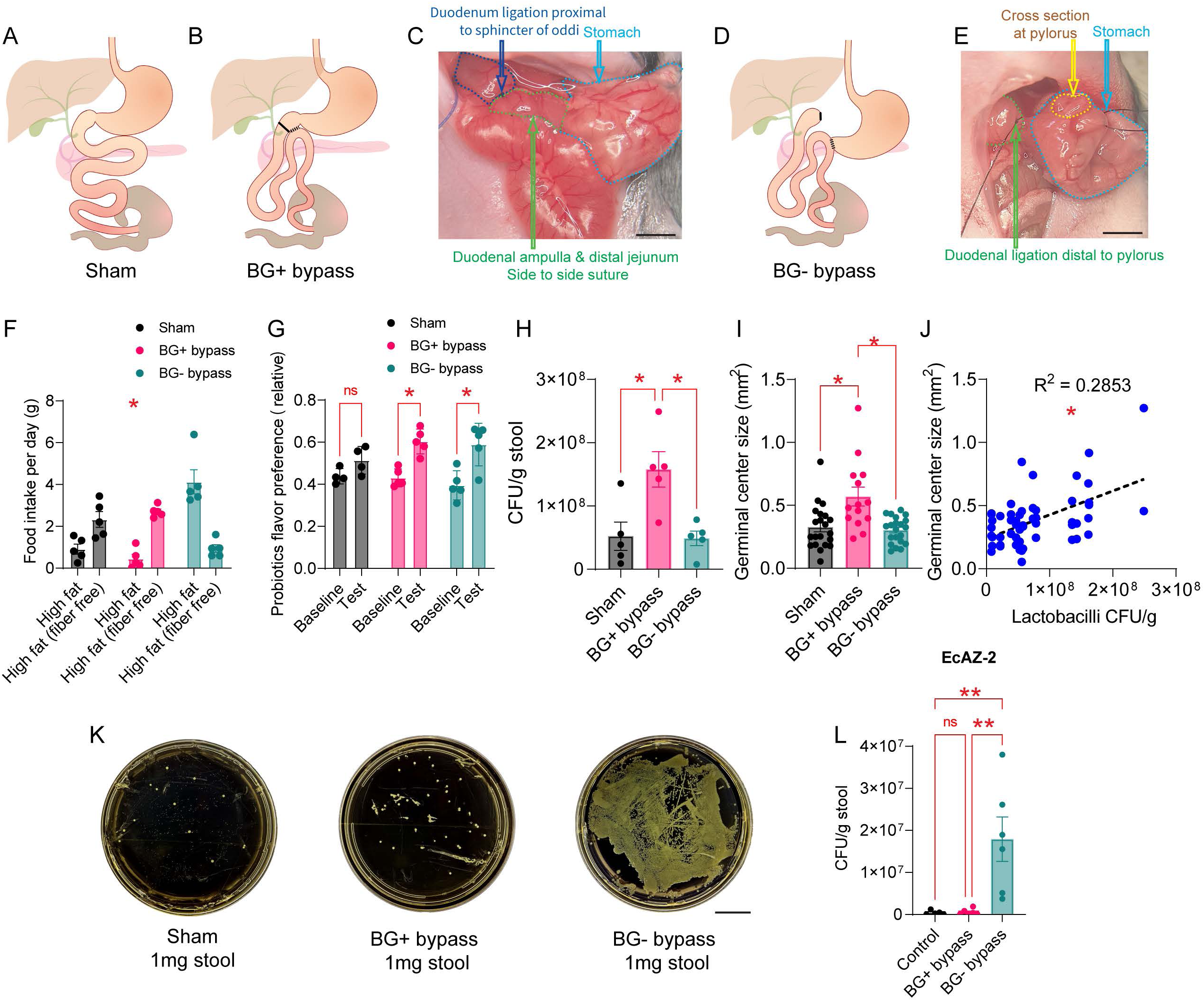
Brunner’s glands contribute to changes in the microbiome induced by bariatric surgery. **A-B.** Schematics showing post-surgical alimentary tracts of Sham (***A***) and Brunner’s glands- preserving (BG+, ***B***) models. **C.** Wide-field view of the BG+ approach. Bar=1cm. **D.** Schematics showing post-surgical alimentary tract of Brunner’s glands-excluding (BG^-^) model. **E.** Wide-field view of the BG- approach. Bar=1cm. **F.** Unlike Sham and BG+, BG- mice displayed significant preferences for fiber-rich fatty pellets, 2-way mixed RM-ANOVA, surgery × food type ∗p<0.001. **G.** Compared to baseline preferences, BG+ and BG-, but not Sham, mice increased post- conditioning preferences for flavors associated with probiotics solutions, 2-way mixed RM- ANOVA, training, p<0.0001; Bonferroni post hoc paired t-tests comparing pre (baseline) *vs.* post-conditioning, Sham: p=0.3329; BG+ and BG-: ∗p<0.004. **H** Higher counts of *Lactobacilli* from fecal samples of BG+ mice compared to both sham and BG- mice, one-way ANOVA, surgery, p=0.0058, Bonferroni BG+ *vs*. Sham and BG-, both ∗p<0.02. **I.** Larger germinal center areas, as defined by HE staining, in BG+ compared to both Sham and BG- mice, one-way ANOVA, surgery, p=0.0002; Bonferroni BG+ *vs*. Sham and BG-, both ∗p<0.002. **J.** Across- animals significant positive linear within-subject correlation between fecal *Lactobacilli* counts and germinal center areas, (Pearson) R^2^=0.2853, ∗p<0.001. **K-L.** Seeding the gut with *E. coli* (kanamycin-resistant EcAZ-2) results in robust proliferation in BG-, but not in Sham or BG+ mice. **K:** Representative examples of 1mg stool on cultured on kanamycin LB agar plates. **L:** EcAZ-2 clone counts, one-way ANOVA, surgery, p=0.0022, Bonferroni Sham *vs*. BG+ p>0.9999; Sham *vs*. BG-, ∗p=0.0069; BG+ *vs.* BG-, ∗∗p=0.0051. Bar= 2cm.

Consistent with the BG ablation results, we observed that BG-, but not BG+ or sham, mice showed greater preferences for fiber-rich fatty food pellets (Figure 4F). These effects occurred independently of changes in blood glucose (Figures S4A-C). However, no robust effects were observed for flavored solutions containing probiotics (Figure 4G). Interestingly, we detected considerably higher levels of *Lactobacilli* in fecal samples from (chow-fed) BG+ compared to both sham and BG- mice (Figure 4H), suggesting that the beneficial effects of bariatric surgery on microbiome composition may depend on preserving BG in the alimentary tract. We also detected augmented germinal center areas in the spleen of BG+ mice compared to both sham and BG- mice (Figures 4I and S4D-E). We then predicted, based on these data, a positive association between intestinal *Lactobacilli* counts and spleen germinal centers. Indeed, we found a strong within-subject correlation between *Lactobacilli* counts and germinal center areas (Figure 4J). Finally, we seeded the gut of mice in the three groups with the *E. coli* strain EcAZ-2 via oral gavage. Again, and in direct agreement with the BG ablation experiments above (see Figures 3P-Q), we detected marked increases in EcAZ-2 counts in the excrements of BG-, but not of BG+ or sham, mice (Figures 4K-L).

### Probiotic or mucin administration counteract the syndrome associated with the ablation of the glands of Brunner

We hypothesized that, if the immunological syndrome that follows BG ablation was indeed due to dysbiosis, then restoring bacterial equilibrium should attenuate its signs. To test this hypothesis, we implanted a catheter into the cecum of BG-vehicle and BG-DTx mice (*i*.*e*., the mice sustaining cell-specific ablation of BG) for administration of probiotic or neutral solutions under a 2x2 design (Figure 5A). We found that administering the probiotics-containing solution to the cecum of BG-DTx mice completely reversed the marked abnormalities in gross spleen morphology that follow BG ablation (Figures 5B-C). The probiotic treatment also restored body weights of BG-DTx mice to normal levels (Figure S5A). We observed similar normalizations of germinal center total areas, IgD^+^ B cell counts, Ki67 proliferation markers, and FDC signal intensities (Figures 5D-I). These effects occurred in the absence of significant changes in food intake or exploratory activity (Figures S5B-C). In addition, the cecum infusions with probiotics in BG-DTx mice significantly decreased mortality after gut infection with the pathogen *S. xylosus* (Figures 5J-L). Finally, probiotics prevented bacterial proliferation after seeding the gut of BG- DTx mice with the *E. coli* strain EcAZ-2 (Figure 5M).

**Figure 5.**
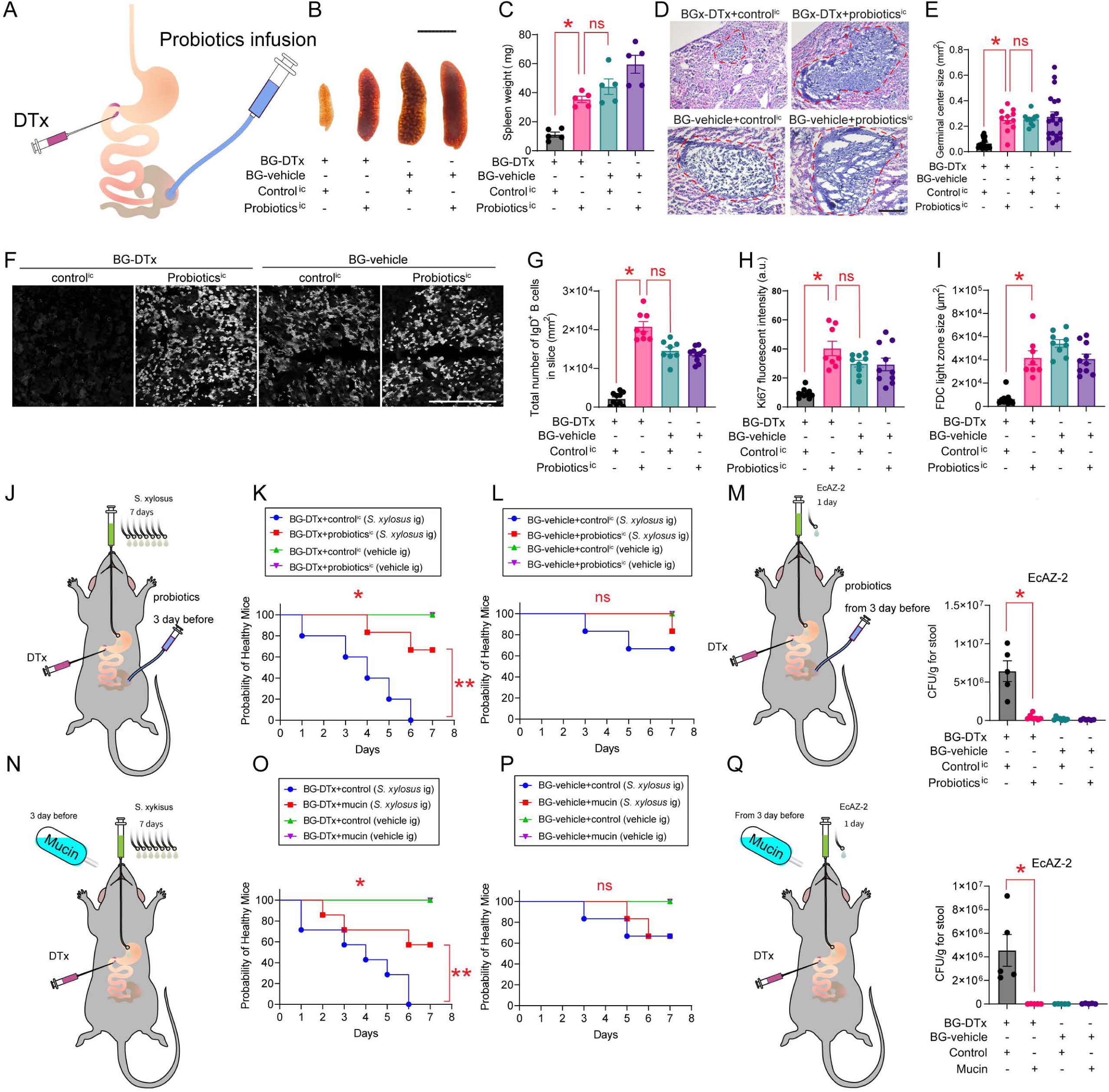
Probiotics and mucin restore immune functions and promote survival in animals lacking glands of Brunner. **A.** Schematics showing cell-specific (DTx-induced) Brunner’s gland ablation combined with intra-cecal administration of probiotic (“probiotics^ic^”, or control^ic^) solutions. **B.** Spleen weights rescued by probiotics^ic^. Representative examples of whole spleens dissected from BG- DTx+control^ic^, BG-DTx+probiotics^ic^, BG-vehicle+control^ic^ and BG-vehicle+probiotic^ic^ mice. **C.** Spleen weights significantly differed across groups, one-way ANOVA, group effect p<0.001. Bonferroni tests show that probiotic-treated BG-DTx mice phenotypically match control mice: BG-DTx+control^ic^ *vs*. BG-DTx+probiotics^ic^, ∗p=0.006, while BG-DTx+probiotics^ic^ *vs.* BG- vehicle+control^ic^, p>0.9. **D-E.** Germinal center sizes are rescued by probiotics^ic^. **D:** Representative examples using HE-staining. **E:** Same as ***C*** but for germinal center sizes, p<0.0001. Bonferroni tests show that probiotic-treated BG-DTx mice phenotypically match control mice: BG-DTx+control^ic^ *vs*. BG-DTx+probiotics^ic^, ∗p=0.0002, while BG-DTx+probiotics^ic^ *vs.* BG-vehicle+control^ic^, p>0.9. **F-G.** B cell counts are rescued by probiotics^ic^. **F:** Representative IgD-immunostaining of spleen section from the experimental groups. **G:** Same as ***E*** but for IgD+ counts, p<0.001, Bonferroni tests show that probiotic-treated BG-DTx mice phenotypically match control mice: BG-DTx+control^ic^ *vs*. BG-DTx+probiotics^ic^, ∗p=0.0001; BG-DTx+probiotics^ic^ *vs.* BG-vehicle+control^ic^ p>0.5. **H.** Proliferation markers are rescued by probiotics^ic^. Same as in ***G*** but for Ki67 signal, p<0.001. Bonferroni tests show that probiotic-treated BG-DTx mice phenotypically match control mice: BG-DTx+control^ic^ *vs*. BG-DTx+probiotics^ic^, ∗p<0.0001; BG- DTx+probiotics^ic^ *vs.* BG-vehicle+control^ic^, p>0.5. **I.** Follicular dendritic cells (FDC) light zones are rescued by probiotics^ic^. One-way ANOVA, p<0.001. Bonferroni tests show that probiotic-treated BG-DTx mice phenotypically match control mice, BG-DTx+control^ic^ *vs*. BG-DTx+probiotics^ic^, ∗p<0.001. **J.** Probiotics^ic^ (or control^ic^) in BG-DTx (3d), then combined with 7d of gavage with pathogen *S. xylosus*. **K.** Probiotics strongly suppressed mortality upon *S. xylosus* infection in BG-DTx mice, Log Rank (Mantel-Cox), ∗p<0.001. **L.** Same analyses for BG-vehicle mice, Log Rank (Mantel-Cox), p=0.216. **M.** Probiotics suppressed *E. coli* proliferation. After gavage of EcAZ-2 to BG-DTx mice, probiotics^ic^ reduced fecal counts to near undetectable levels, one-way ANOVA, group p<0.001. Bonferroni BG-DTx+control^ic^ *vs*. BG-DTx+probiotics^ic^, ∗p<0.001. **N-Q.** Mucin ingestion rescues immunological function in mice lacking Brunner’s glands. **N:** Schematics of Brunner’s gland ablated mice combined with 10% mucin in drinking water (3d), followed by 7d of daily gavage with the pathogen *S. xylosus* (combined with continued mucin availability). **O:** Mucin ingestion suppressed mortality upon *S. xylosus* infection in BG-DTx mice, Log Rank (Mantel-Cox), ∗p<0.001. **P.** As in ***O***, mucin effect on BG-vehicle mice, Log Rank (Mantel-Cox), p=0.199. **Q.** Mucin suppressed *E. coli* proliferation. After gavage of EcAZ-2 to BG-DTx mice, mucin ingestion reduced fecal counts to undetectable levels, one-way ANOVA, p<0.001. Bonferroni BGx-DTx+control vs. BGx-DTx+mucin, ∗p=0.001.

Since we initially hypothesized that BG may play a key role in mucosal-microbiome interactions, we reasoned that administering mucin solutions to animals would produce effects like those induced by probiotics. Indeed, providing a mucin solution to BG-DTx mice in drinking bottles decreased mortality following gut infection with the pathogen *S. xylosus* (Figure 5N-P). Moreover, mucin solutions prevented bacterial proliferation after seeding the gut of BG-DTx mice with the *E. coli* strain EcAZ-2 (Figure 5Q). In sum, dysbiosis, and, specifically, low probiotic abundance due to mucus suppression, appears as the necessary and sufficient condition for the occurrence of the immunodeficient phenotype associated with BG ablation.

### Probiotic administration normalizes immune signatures in animals lacking the glands of Brunner

Based on the above, we performed additional studies to characterize the immune signatures of BG-vehicle (controls), BG-DTx, and BG-DTx+probiotics mice. This was first accomplished by performing CyTOF analyses based on bone marrow, spleen, and mesenteric lymph node samples (Figure S6A).

We found that BG ablation caused marked reductions in B cell counts in mesenteric lymph nodes, an effect that was entirely normalized by probiotic administration (Figures 6A-B and S6B). Similar effects were observed for spleen and bone marrow samples, although in those cases B cell suppression was not fully reversed by probiotic administration (Figure 6A). Qualitatively similar effects were observed when analyses were performed separately for mature *vs.* immature B cells (Figures 6C-D). In addition to B cells, comparable effects were observed for other immune cell types including dendritic cells (Figure 6E), natural killer cells (6F), and Ly6C^Low^ (“patrolling”) monocytes (6G). Effects were less conspicuous for other immune cell types (Figures S6C-F), with the notable exception of macrophages: while counts increased markedly in BG-DTx animals, they were brought to near-normal levels after probiotic administration (Figure S6G).

**Figure 6.**
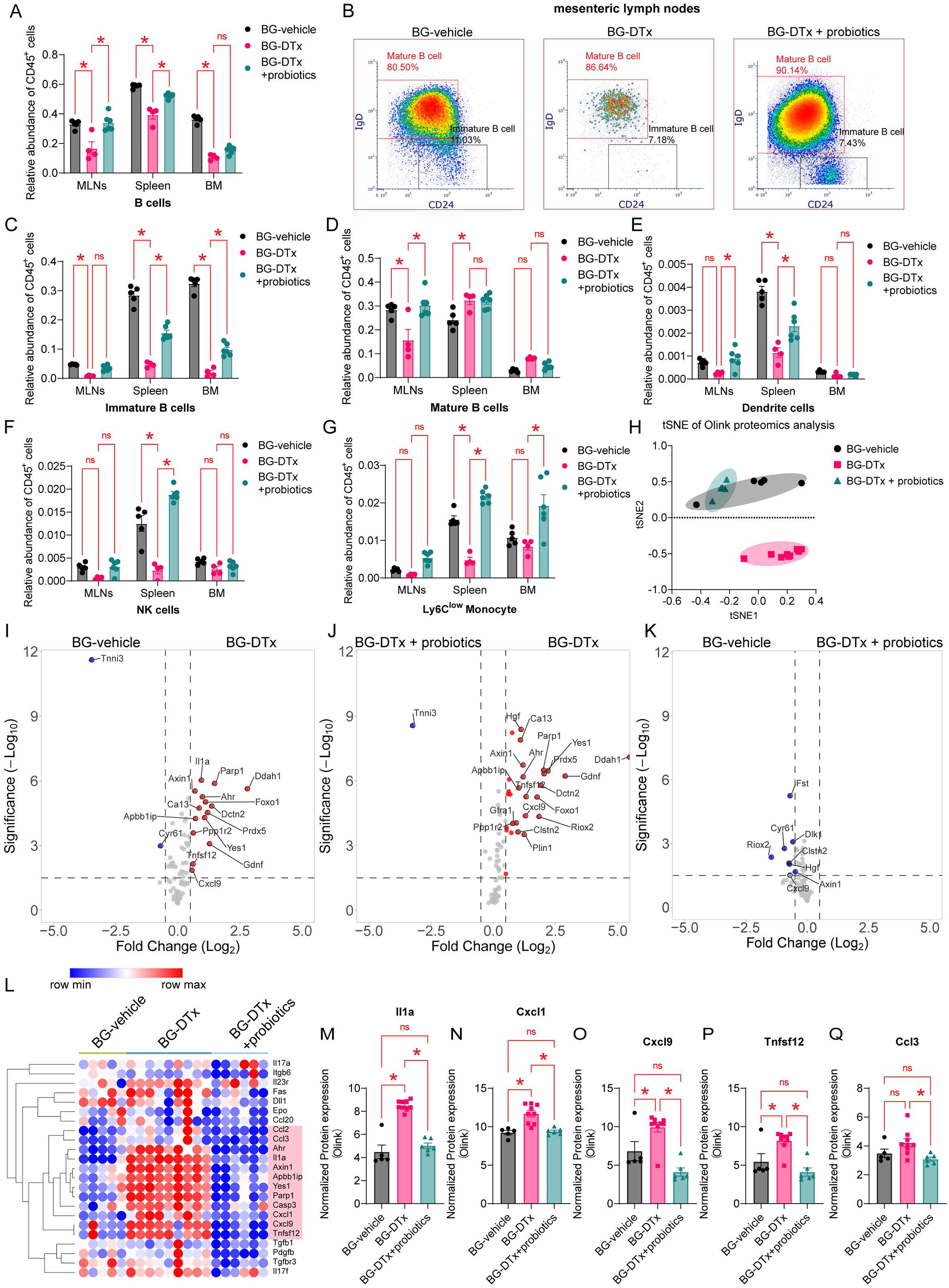
Immune signatures are rescued by probiotic treatment in animals lacking the glands of Brunner. **A-C.** Probiotics ingestion recover immune parameters in animals lacking Brunner’s glands. **A:** CyTOF-based relative abundances of B cells from total pool of live CD45^+^ cells in mesenteric lymph nodes, spleen, and bone marrow samples from BG-vehicle, BG-DTx, and BG- DTx+probiotics mice on a Glp1r-ires-cre×ROSA26-iDTR background. Two-way RM-ANOVA, group and organ x group effects p<0.001. For all Bonferroni post-hoc pairwise comparisons shown, ∗p<0.0001. The detailed gating strategy is shown in Figure S6A. **B.** In mesenteric lymph nodes, B cell suppression is observed in BG-DTx mice, an effect partially recovered by probiotics. **C.** Relative abundance of immature B cells from total pool of CD45^+^ cells in mesenteric lymph nodes, spleen, and bone marrow, 2-way RM ANOVA, group, and organ × group effects p<0.001. For all Bonferroni post-hoc pairwise comparisons shown, ∗p<0.015. **D.** As in ***C*** but for mature B cells, p<0.001; For all pairwise comparisons shown, ∗p<0.001. **E.** As in ***C*** but for Dendritic cells (DCs), p<0.001; For all Bonferroni post-hoc pairwise comparisons shown, ∗p<0.03. **F.** As in ***C*** but for Natural killer (NK) cells, p<0.001; For all Bonferroni post-hoc pairwise comparisons shown, ∗p<0.0001. **G.** As in ***C*** but for Ly6C^low^ monocytes, p=0.003; For all Bonferroni post-hoc pairwise comparisons shown, ∗p<0.0001. **H.** Unsupervised clustering (as per tSNE) for Olink proteomic analyses of blood samples from the three treatment groups. Note BG-vehicle and BG-DTx+Probiotics samples originated one cluster that was separate from the BG-DTx cluster. **I.** Volcano plot contrasting Olink-based blood proteomics profiling of BG-vehicle *vs*. BG-DTx groups. **J.** Volcano plot contrasting BG-DTx *vs*. BG-DTx+Probiotics groups. **K.** Volcano plot contrasting BG-vehicle *vs*. BG-DTx+Probiotics. Note rescue induced by probiotic treatment. **L.** Immune profile of cytokines was markedly altered in BG-DTx mice, an effect brought to near-normal levels by probiotic treatment. **M-Q.** Levels of five inflammation-related cytokines, *Il1a* (***M***), *Cxcl1* (***N***), *Cxcl9* (***O***), *Tnfsf12* (***P***), and *Ccl3* (***Q***) were significantly increased in BG-DTx mice, an effect brought to near-normal levels by probiotic treatment. For all the five cytokines, group effect (one-way ANOVA), p<0.03; Bonferroni pairwise comparisons shown, ∗p<0.03.

Organ-based CyTOF experiments were supplemented with proteomic (Olink) analyses of blood samples from the same animals. t-distributed stochastic neighbor embedding (t-SNE) map visualization of the overall data shows a clear separation between components in BG-DTx *vs*. control and BG-DTx+probiotics mice, both of which clustered together (Figure 6H). Consistently, both volcano plot and cell count analyses showed that BG ablation caused significant increases in levels of several pro-inflammatory and pro-apoptotic cytokines, including IL-1a, Cxcl1, Cxcl9, Tnfsf12, and Ccl3; these effects were however largely reversed by probiotic administration (Figures 6I-Q and S6H-I).

In sum, probiotics administration was sufficient to normalize several immunological parameters in mice lacking glands of Brunner, an effect that involves both cell contents of immune-related tissues and circulating cytokines.

### A neuronal circuit connects the central nucleus of the amygdala to the glands of Brunner via the vagus nerve

Given that vagal parasympathetic fibers innervate and control BG, we hypothesized that the activity of the glands may also be modulated by brain centers upstream of vagal parasympathetic neurons. To verify our hypothesis, we first injected the *Cre*-dependent, polysynaptic, retrograde pseudo-rabies virus strain PRV-CAG-DIO-TK-GFP (“PRV”) into the proximal duodenal submucosa of Glp1r-ires-Cre mice. In addition to expected labeling of vagal parasympathetic neurons in the dorsal vagal complex (“DVC”), we also detected clear labeling of BG origin in regions linked to autonomic control, especially the paraventricular and parasubthalamic hypothalami, the *Locus Coeruleus*, and insular cortex. Interestingly, we observed dense labeling in the medial aspect of the central nucleus of the amygdala (“CeA”), a brain region deeply implicated in emotional regulation^29,30^. To verify that the wiring is this circuit depends on the vagus nerve, we repeated the tracing experiment in Glp1r-ires-Cre mice sustaining bilateral subdiaphragmatic vagotomies. We found that severing the nerve at the abdominal level was sufficient to entirely abolish labeling in DVC and CeA (Figure 7A). When then repeated the experiment after removal of the abdominal celiac ganglia, which severs the splenic nerve^31^. In contrast to the vagotomy studies, labeling in most brain areas, including CeA, was entirely preserved (Figure S7A, which includes additional control studies). Thus, CeA is neuronally connected to the glands of Brunner via vagal, but not spinal/sympathetic, pathways.

**Figure 7.**
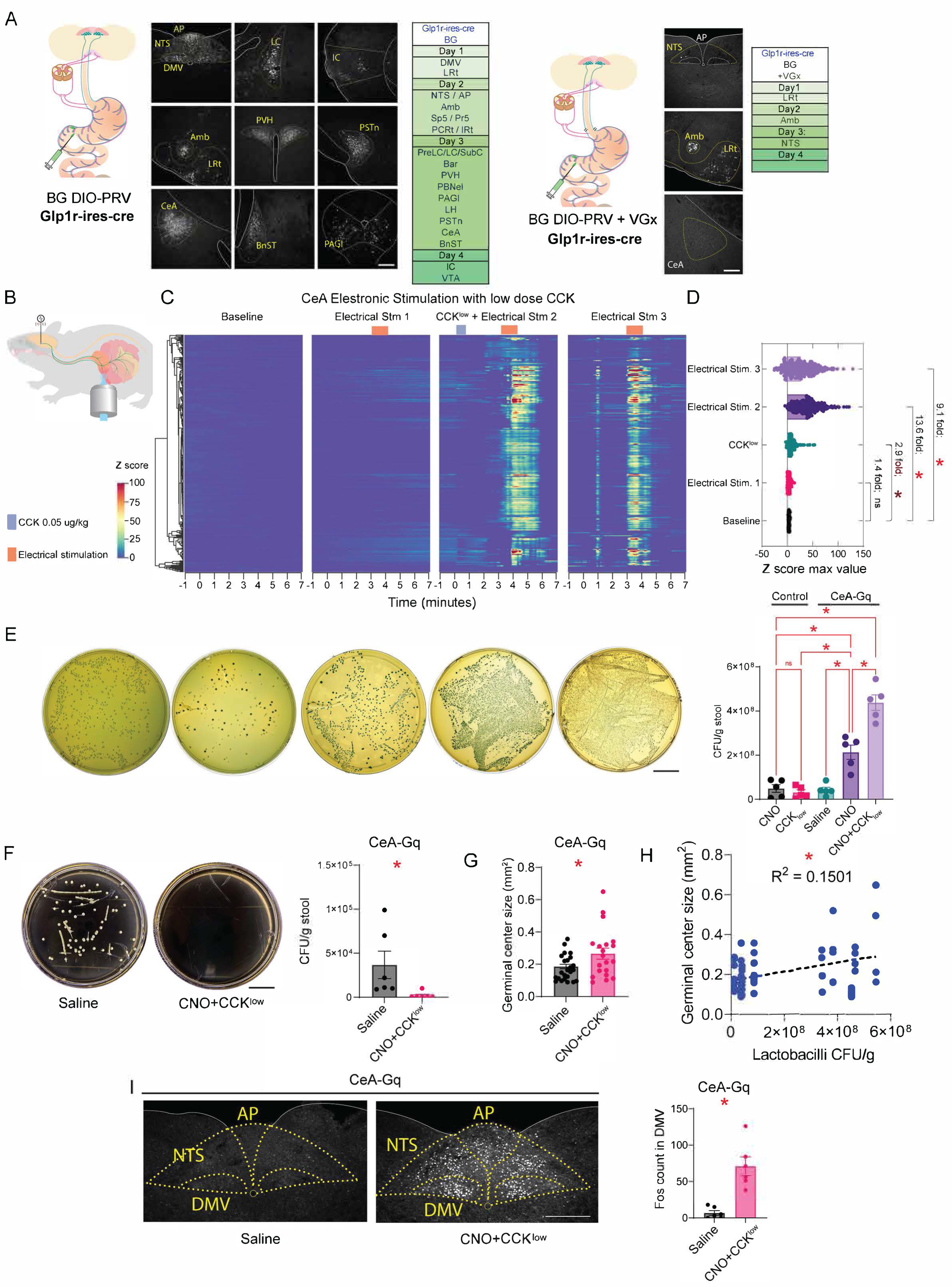
**A.** Injections of the polysynaptic *Cre-*dependent PRV strain into duodenal submucosa of Glp1r- ires-cre resulted mice in dense labeling in well-defined subcortical regions (Left), including the dorsal motor vagus (DMV) and central nucleus of amygdala (CeA). Most labeling, including DMV and CeA, was absent when tracing was performed after subdiaphragmatic vagotomy (Right). Bars=100µm. **B-C.** During intravital imaging, electrical stimulation of CeA in combination with a subthreshold dose of CCK (CCK^low^, 0.05µg/Kg) induced robust, supra-additive calcium transients in Brunner’s glands. **D.** Maximal Z-scores of calcium transients across the different experimental conditions (300 glands analyzed from 3 mice). One-way ANOVA, p<0.0001. Bonferroni post hoc pairwise comparisons: Except for Baseline *vs*. Electrical Stim #1 (p >0.9999), all other comparisons yielded significant effects, ∗p<0.0001. **E.** *Lactobacilli* CFU counts across the experimental different conditions. Chemogenetic CeA stimulation+CCK^low^ induced robust, supra-additive increases in *Lactobacilli* counts. One-way ANOVA, p<0.0001. Bonferroni post-hoc pairwise comparisons: All comparisons involving CNO injections in chemogenetically enabled mice (CeA-Gq-CNO) yielded significant effects, ∗p<0.0015. CNO control injections alone yielded no significant effects, all p>0.9. Bar=2cm. **F.** After gavage with *E. coli* EcAZ-2, chemogenetic CeA stimulation+CCK^low^ reduced fecal counts to undetectable levels compared to control, 2-sample t-test, ∗p=0.0495. Bar=2cm. **G.** Chemogenetic CeA stimulation+CCK^low^ augmented spleen germinal center sizes: 2-sample t-test, ∗p=0.015 (HE- stained sections collected from 4 mice). **H.** Significant linear association between *Lactobacilli* CFU counts and spleen germinal center sizes across conditions (n=56), Pearson ∗p=0.032. **I.** Chemogenetic CeA stimulation+CCK^low^ increased neural activity in dorsal vagal complex (*Nucleus Tractus Solitarius*, NTS, and DMV), as measured by *Fos* reactivity, 2-sample t-test against control, ∗p=0.0007. Bar=200 µm.

### Electrical and chemogenetic activation of the central nucleus of the amygdala activates the glands of Brunner and modulates the intestinal microbiome

We then inquired whether these circuits are functional. We implanted unilaterally the tip of a stimulation electrode in the CeA of Glp1r[GCamp6] mice. We concomitantly performed intra-vital imaging of BG (Figure 7B). At first, we observed no effects of electrical stimulation of CeA on BG activity. We then decided to prime BG baseline activity levels by injecting mice with a sub- threshold dose of CCK that is ineffective on its own (“CCK^Low^”). Under these conditions, electrical stimulation of CeA caused very robust, supra-additive increases in BG calcium transients (Figures 7C-D and Supplemental Movie 3). Consistently, BG transients remained unaltered in animals sustaining splenectomies (Figures S7B).

The ability of CeA to engage activity in BG points to a potential role for the amygdala in modulating the microbiome. To modulate the activity of CeA neurons in awake animals, we performed chemogenetic activation of excitatory design receptors by injecting the construct pAAV-hSyn-hM3D(Gq)-mCherry into the CeA of wild-type mice. Chemogenetic activation was achieved by intraperitoneal injections of the designer drug clozapine-N-oxide (“CNO”). We found that CeA excitation was sufficient to markedly increase *Lactobacilli* counts in excrements of the stimulated mice; consistent with the BG imaging experiments above, robust supra-additive effects were observed upon priming BG with CCK^Low^ (Figure 7E).

We then tested the ability of CeA stimulation to suppress proliferation of exogenous gut bacteria. We again seeded mice with the *E. coli* strain EcAZ-2 via oral gavage. After priming mice with CCK^Low^, we found that CeA chemogenetic excitation with CNO was sufficient to suppress EcAZ- 2 proliferation (Figure 7F). In addition, we found that the same treatment caused a moderate, yet statistically significant, expansion of spleen germinal centers (Figure 7G). Moreover, we again observed a significant, CeA-mediated within-subject correlation between *Lactobacilli* counts and germinal center areas (Figure 7H). This effect points to CeA-driven modulation of peripheral immune-related tissues. Finally, using the early-gene neuronal activity marker *Fos*, we found that CeA chemogenetic excitation with CNO (+CCK^Low^) caused a robust activation of vagal motor neurons in DVC (Figure 7I), suggesting that CeA modulates parasympathetic control over gut secretory glands.

## Discussion

We identified a neuroglandular circuit that controls the secretory activity of the glands of Brunner. We found that this circuit controls gut bacterial composition and that its ablation leads to dysbiosis, deficits in immune signaling, and mortality. The pathology was transmissible via fecal transplant but reversable by probiotics. These findings involving the modulation of intestinal bacteria differ significantly from the classical role attributed to Brunner’s glands, namely, to presumably increase the intrinsic resistance of the duodenum to acid-peptic digestion^21,32^. The results indicate that the glands of Brunner may be the critical cells that regulate the levels of Lactobacilli species in the intestine.

A neuroepithelial circuit for gut mucus secretion has been demonstrated for Goblet cells. Specifically, gut sensory nociceptor fibers engage the CGRP-Ramp1 pathway to induce goblet cell emptying and mucus secretion^33^. Interestingly, genetic disruption of gut nociceptors also leads to severe inflammation and alterations in the intestinal microbiota, including a transmissible dysbiosis^34^. Altogether, and consistent with our main observations, these findings point to a critical role for extrinsic gut nerves in controlling the microbiome via the targeting of mucus-secreting cells. Moreover, recent findings also implicated colonic enteric neurons in the connection between psychological stress and intestinal inflammation^35^. Our present findings add to this emerging picture by revealing a novel neuroglandular circuit via which vagal fibers target submucosal glands of the upper intestine to promote bacterial homeostasis.

Our findings are also directly consistent with the idea that efferent vagal fibers promote immune function and anti-inflammatory signaling^36^. Of particular interest are the reported beneficial effects of vagal cholinergic signaling on spleen function^37^, which parallel our findings linking the glands of Brunner and spleen characteristics. Our proposed circuit may, on one hand, clarify the mechanisms by which the vagus nerve ameliorates inflammatory states via the spleen. But they may also link, via neuron-induced mucosal secretion, microbiome composition to the promotion of hematopoiesis^38^. In fact, the beneficial effects of vagal efferent activity may be particularly relevant for gut mucosal integrity. Non-invasive (auricular branch) vagal nerve stimulation reduced permeability of the small intestine of human subjects pretreated with intravenous administration of stress hormones^39^. This effect is consistent with preclinical studies in which vagal stimulation was shown to both reverse stress-induced changes in intestinal permeability^40^ and attenuate inflammatory bowel disease symptoms^41^. The underlying mechanisms linking vagal cholinergic signaling and intestinal barrier integrity remain however unknown^41^. Thus, our present study suggests that the glands of Brunner, and their mucus secretion, may account for at least part of the protective effects of vagal stimulation on gut mucosal immunity.

In addition to cholinergic signaling, the sensory branch of the vagus nerve appears to also be relevant for vagal modulation of the microbiome. Indeed, we found that cholecystokinin, a pro- digestive gut hormone^42^ whose action depends on vagal sensory ganglia^43^, promotes the expansion of *Lactobacilli* populations via Brunner’s glands. This points to the possibility that the populations of probiotic, digestion-promoting bacteria expand in response to the activity of vagal nutrient-sensing fibers. Moreover, vagal sensory fibers may also sense the intestinal microbial content itself. Probiotic administration improved physiological markers in rodent models of anxiety, an effect that was mediated by vagal sensing^13^. Indeed, vagal sensory ganglia were shown to sense bacterial metabolites, mediating the generation of autonomic responses in response to bacterial signaling^44^. These findings are in line with our observation that mice sustaining lesions to, or bariatric exclusions of, the glands of Brunner developed preferences for fiber-enriched pellets and for probiotic drinking solutions. In other words, the effects of dysbiosis may be sensed via interoceptive pathways^45–47^, enabling the triggering of compensatory behavioral responses.

Finally, we found that the central nucleus of the amygdala, a subcortical area within the temporal lobes, controls the glands of Brunner and modulates the microbiome. Central amygdala neurons are fundamentally implicated in emotional behavior^29,30,48,49^. Specifically, central amygdala plays major roles in conditioned fear and anxiety^29,50,51^, and may thus constitute a critical link between negative psychological states and dysbiosis. In line with this idea, the role of central amygdala in emotional modulation has been connected to vagal signaling. Specifically, gut-borne vagal sensory signals were shown to modulate anxiety states via central amygdala^52^. Consistent with these findings, *L. rhamnosus*, a *Lactobacillus* species – the levels of which we found to be upregulated by cholecystokinin – reduced stress-induced anxiety- and depression-like behaviors by modulating forebrain levels of GABA, the major neurotransmitter of central amygdala, via the vagus nerve^13^. Interestingly, the central amygdala has also been shown to be critical for the execution of predatory hunting^53^. Although the relationship between predation and gut immunity may seem disparate, it has been proposed that the adaptive immune system of vertebrates first evolved in the gut of primitive jawed fishes in response to gastrointestinal infections brought about by the novel predatory lifestyle^54^. Future studies may reveal elusive evolutionary forces that led to the concurrent emergence, in predatory vertebrates, of a central amygdala and gut mucosal immunity.

## Limitations of the Study

An important limitation is the absence of experiments involving lesions of the glands of Brunner in germ-free animals. In fact, both our approaches to lesioning the glands involved invasive surgeries, procedures that would compromise the germ-free status of the subjects. Unfortunately, our single cell transcriptome analyses failed to reveal clear genetic markers that would be unique to Brunner’s glands (*i*.*e*., when considering the whole organism). While this limitation does not impact directly on our conclusions, future intersectional genetic approaches may enable the targeting of Brunner’s glands without requiring invasive interventions.

## Supporting information

Supplentary Figures with legends

## Acknowledgements

NIH-NCCIH R01 AT011697-01 (to IEdA), Food Allergy Science Initiative Consortium (to IEdA), Modern Diet and Physiology Research Center (to IEdA). We thank Dr. Amir Zarrinpar for providing the engineered *E. Coli.* strain EcAZ-2. We thank Geoffrey Kelly and Kai Nie, at the Mount Sinai Human Immune Monitoring Center, for assistance in processing CyTOF and Olink samples. We also thank the Mount Sinai Microscopy CoRE for assisting us with accomplishing intravital imaging of calcium transients.

## Figure Legends

Details on all statistical tests performed are shown in Supplemental Table 3, including type of test, N, degrees of freedom, statistic, and p values. Legends report test and p values when applicable. In all figures, plotted are means ± SEM.

**Supplemental Movie 1**. Related to Figure 1. Intravital imaging of calcium transients in the glands of Brunner. GCaMP6f and tdTomato (red channel) signals from Brunner’s glands. Note tdTomato remains stable and unchanged despite strong calcium transients in the same glands.

**Supplemental Movie 2** Related to Figure 2. During intravital imaging, cholecystokinin induced strong calcium transients in Brunner’s glands.

**Supplemental Movie 3** Related to Figure 7. During intravital imaging, electrical stimulation of the central nucleus of the amygdala induced strong calcium transients in Brunner’s glands.

**Supplemental Table 1**. Details on surgical procedures.

**Supplemental Table 2**. Missing data frequency in Olink proteomics analysis.

**Supplemental Table 3.** Details on all statistical tests performed, including type of test, N, degrees of freedom, statistic, and p values.

## STAR Methods

### RESOURCE AVAILABILITY

#### Technical contact

Further information and requests for reagents should be directed to, and will be fulfilled by, Hao Chang, PhD, and Wenfei Han, M.D., PhD

#### Materials availability

This study did not generate new unique reagents.

#### Data and code availability

- Behavioral, electrophysiological, anatomical, and post-processed intravital imaging data have been deposited at *Mendeley Data* and are publicly available as of the date of publication. DOIs are listed in the key resources table.
- This paper does not report original code.
- Any additional information needed for reanalyzing the dataset reported in this article will be made available by the technical contacts upon request.

### Experimental Model and Subject Details

All experiments presented in this study were conducted according to the animal research guidelines from NIH and were approved by the Institutional Animal Care and Use Committee of Icahn School of Medicine at Mount Sinai.

#### Experimental Animals

A total of 541 adult male mice were used. Strain details and number of animals are as follows:

206 C57BL/6J (Jax Mouse Strain #000664)

15 Glp1r-ires-Cre (Jax Mouse Strain #029283)

33 Glp1r-ires-Cre x Ai148D (Jax Mouse Strain #029283 and #030328)

10 Glp1r-ires-Cre x CD63-emGFP^l/s/l^ (Jax Mouse Strain #029283 and # 036865)

3 Glp1r-ires-Cre x Ai148D x Ai9 (Jax Mouse Strain #029283 and #030328 and #007907) 215 Glp1r-ires-Cre x ROSA26iDTR (Jax Mouse Strain #029283 and #007900)

36 Glp1r-ires-Cre x Ai148D x ROSA26iDTR (Jax Mouse Strain #029283 and #030328 and #007900)

3 Ai9 x Glp1r-ires-Cre (Jax Mouse Strain #007914 and #029283)

5 Glp1r-ires-Cre x CAG-Sun1/sfGFP (Jax Mouse Strain #029283 and #021039) 15 B6J.ChAT-IRES-Cre::Δneo (Jax Mouse Strain #031661)

All mice used in behavioral experiments were individually housed under a 12-hour light/dark cycle. At the time of the behavioral experiments, animals were 8–20 weeks old and weighted approximately 25-28 grams. All animals were used in scientific experiments for the first time. This includes no previous exposure to pharmacological substances or alternative diets. Health status was normal for all animals. Animals used for anatomical tracing studies were group housed and 3 weeks old at the time of injection. Please see details on surgical procedure in Supplemental Table 1.

### Method Details

The following provides details on viral and drug injections, catheterizations, denervation, brain electrode implantation, and intravital imaging preparations for each mouse strain (Also see *Key Resources* table). All surgeries were performed in a Biosafety Level 2-approved laboratory. All mice including surgical mice were housed at the Mount Sinai specific-pathogen-free (SPF) Animal Facility. The surgical mice were monitored daily for body weight and food intake. In all cases, preoperative analgesia: 0.05mg/Kg Buprenorphine (*s*.*c*.); anesthesia induced by 3% isoflurane and maintained by 1.5% ∼2% isoflurane; postoperative analgesia: 0.05mg/Kg Buprenorphine (s.c.) twice a day for three consecutive days. The surgical areas were shaved and cleaned with iodine soap and wiped with 70% isopropyl alcohol. All incisions were thoroughly disinfected with a layer of Baytril ointment. All surgeries were performed under stereomicroscopes, with animals placed on a heated pad (CMA 450; Harvard Apparatus, Holliston, MA). After surgery, animals were allowed to recover under infrared heat until they chose to reside in the unheated side of the cage.

### Lesions of the glands of Brunner

#### Surgical resection of Brunner’ glands (BG-resection)

The abdomen of an 8-hour food-restricted animal was shaved and cleaned. A midline incision was made into the abdomen. The stomach was exteriorized through the midline incision, and the pyloric antrum was loosely stitched to the left rectus abdominis to maximize the view of the proximal duodenum. Wet surgical gauze was applied to isolate the duodenum and the duodenum bulb held with cotton tips. The scalpel tip was carefully manipulated to avoid the mesenteric blood vessels. A longitudinal incision was made on the ventral side of the duodenum bulb (to the distal direction, 5mm from the duodenum-pyloric sphincter junction). The lumen was opened with fine tweezers, and the lumen contents gently flushed with saline. The submucosal tissue underneath the pyloric sphincter was resected with an electric cauterizer. The duodenal incision was sutured with an 8-0 absorbable vicryl suture (Ethicon V548G, USA), the loose suture attached to the stomach removed, and the rectus abdominis/skin incisions were closed with sterile sutures.

#### Duodenum resection (Duo-lesion)

This is a control procedure for the Brunner’s gland resection described above. The mouse duodenum was exposed as described above. The descending part of the duodenum (approximately at the location of the sphincter of Oddi) was held with cotton tips. The scalpel tip was carefully manipulated to avoid mesenteric blood vessels. A longitudinal incision was made on the ventral side of the duodenum (∼5mm long and at least 10mm distal to the duodenal- pyloric junction, thereby avoiding the glands of Brunner). The lumen was opened with fine tweezers and slightly flushed with saline. The submucosal tissue underneath the pyloric sphincter was resected with the electric cauterizer. The duodenal incision was sutured with an 8-0 absorbable vicryl suture (Ethicon V548G, USA), the loose suture attached to the stomach removed, and the rectus abdominis/skin incisions were closed with sterile sutures.

#### Sham surgery (negative control)

The mouse duodenum bulb was exposed, and a ∼5mm longitudinal incision was cut as described above. The duodenal incision was sutured with an 8-0 absorbable vicryl suture (Ethicon V548G, USA), the loose suture attached to the stomach removed, and the rectus abdominis/skin incisions were closed with sterile sutures.

#### Brunner DTx / Saline injection (BG-DTx and BG-vehicle)

The abdomen of an 8-hour food-restricted animal was shaved and cleaned. A midline incision was made into the abdomen. The stomach was exteriorized through the midline incision, and the pyloric antrum was loosely stitched to the left rectus abdominis to maximize the view of proximal duodenum. 0.2mg/mL diphtheria toxin (Sigma D0564), or Saline, was loaded into a Nanofil^TM^ 36G beveled needle (WPI, Sarasota, FL) and Nanofil^TM^ tubing (WPI, Sarasota, FL) was connected to a Nanofil^TM^ 10µl syringe (WPI, Sarasota, FL) mounted on a Pump 11 Elite Nanomite (Harvard Apparatus, Holliston, MA). Eight 25nL injections (total volume of 200 nL) were made at 10nL/s into evenly distributed punctures targeting the submucosal layer of the duodenum bulb (within 3 mm approximate to the pyloric sphincter). The needle tip was carefully manipulated to avoid mesenteric blood vessels. After the infusions were completed, the needle was left in place for 5s before extraction to ensure complete absorption. After removing the loose suture attached to the stomach, sterile sutures were then applied to the skin.

#### Duodenum DTx injection in myenteric layer (Duo-DTx)

This is a control procedure for the Brunner’s gland ablation described above. The duodenal bulb was exposed as described above. 0.2mg/mL diphtheria toxin (Sigma D0564), or saline, was loaded into a Nanofil^TM^ 36G beveled needle (WPI, Sarasota, FL), and Nanofil^TM^ tubing (WPI, Sarasota, FL) connected to a Nanofil^TM^ 10µl syringe (WPI, Sarasota, FL) and mounted on a Pump 11 Elite Nanomite (Harvard Apparatus, Holliston, MA). Eight 25nL injections (total volume of 200 nL) were made at 10nL/s into evenly distributed punctures targeting the myenteric layer of the duodenum 10mm below the bulb. The needle tip was carefully manipulated to avoid mesenteric blood vessels. After the infusions were completed, the needle was left in place for 5s before extraction to ensure complete absorption. After removing the loose suture attached to the stomach, sterile sutures were then applied to the skin.

### Bariatric models

#### Brunner’s glands-excluding surgery (BG- bypass)

In this bypass model, the glands of Brunner were excluded from the alimentary tract. The abdomen of an 8-hour food-restricted animal was shaved and cleaned. A midline incision was made into the abdomen. The stomach was exteriorized through the midline incision, and the pyloric antrum was loosely stitched to the left rectus abdominis to maximize the view of proximal duodenum. The duodenal bulb was held with cotton tips. 5-0 silk suture was carefully applied around the pyloric sphincter with a curved blunt needle. We verified that no mesenteric blood vessels or accompanying nerves were located on the inside of the suture noose before tightly ligating the pyloric sphincter. The pyloric sphincter was transversely cut, and the lower jejunum exposed ∼6-8cm distal to pylorus, and a ∼3mm longitudinal incision was made on the ventral side. The gastric pyloric canal was anastomosed to the lower jejunum with an 8-0 absorbable vicryl suture (Ethicon V548G, USA). After removing the loose suture attached to the stomach, sterile sutures were then applied to the skin.

#### Brunner’s glands-preserving surgery (BG+ bypass)

In this bypass model, the glands of Brunner were maintained in the alimentary tract. The abdomen of an 8-hour food-restricted animal was shaved and cleaned. A midline incision was made into the abdomen. The stomach was exteriorized through the midline incision, and the pyloric antrum was loosely stitched to the left rectus abdominis to maximize the view of proximal duodenum. The duodenal bulb was held with cotton tips. 5-0 silk suture was carefully applied around the pyloric sphincter with a curved blunt needle, ∼5-6mm distal to the pyloric sphincter. We verified that no mesenteric blood vessels or accompanying nerves were located on the inside of the suture noose before tightly ligating the duodenum. The pyloric sphincter was transversely cut, and the lower jejunum exposed ∼6-8cmdistal to pylorus, and a ∼3mm longitudinal incision was made on the ventral side. The duodenal bulb was anastomosed to the lower jejunum with an 8-0 absorbable vicryl suture (Ethicon V548G, USA). After removing the loose suture attached to the stomach, sterile sutures were then applied to the skin.

#### Sham surgery

The duodenum and lower jejunum were exposed as described above. A ∼3mm longitudinal incision was made on the ventral sides of both the duodenum bulb and lower jejunum. The incisions were separately sutured, and the abdominal/skin incisions stitched with sterile sutures.

### Vagus and Splenic Nerve lesions

#### Nodose CCKSAP lesion

Mice injected with unconjugated saporin (Blank-SAP) or cholecystokinin-conjugated saporin (CCK-SAP) were 3 weeks old at the time of surgery. Nodose ganglia injections were performed as described previously^55^. Briefly, a ventral midline incision was made along the length of the neck; submandibular glands were retracted along with sternohyoid and omohyoid muscles to expose the trachea and the carotid artery. The vagus nerve was separated from the carotid artery with the Spinal Cord Hook (FST, Foster City, CA) until the nodose ganglion became visibly accessible. Viral or chemical aliquots were loaded into a Nanofil^TM^ 36G beveled needle (WPI, Sarasota, FL) and Silflex^TM^ tubing (WPI, Sarasota, FL). For each nodose ganglion, a total of 500nL volume of Blank-SAP or CCK-SAP was delivered at 50nL/min using a Nanofil^TM^ 10µl syringe (WPI, Sarasota, FL) mounted on a Pump 11 Elite Nanomite (Harvard Apparatus, Holliston, MA). Sterile suture was then applied to the skin.

#### Subdiaphragmatic vagotomy

3-week-old animals were 8-hour food restricted before the surgery. The abdomen was shaved and cleaned. A midline incision was made into the abdomen. The liver and stomach were retracted aside to expose the esophagus. The branches of the vagus nerves innervating the stomach were carefully separated from both the esophagus and the left gastric artery, and bilaterally severed with an electrical cauterizer. Sterile suture was then applied to muscle and skin.

#### Splenectomies

Animals at the time of splenectomies were 3 weeks old. The abdomen was shaved and cleaned. A midline incision was made into the abdomen. The stomach was exteriorized through a midline incision. The stomach fundus was pulled towards the liver to expose the celiac and superior mesenteric ganglia attached respectively to the celiac and superior mesenteric arteries. The greater, lesser, and least splanchnic nerves carrying the 5th through 11th thoracic sensory and sympathetic innervations were identified, carefully separated away from the celiac artery and aorta, and dissected with miniature forceps and spring scissors (FST, Foster City, CA). Sterile sutures were then applied to the skin.

### Brain stereotactic surgeries

#### Stereotaxic viral injection and probe implantation

Animals at the time of surgery were 6-week-old. Injections were performed with a Hamilton 1.0µL Neuros Model 7001KH syringe, at a rate of 20nL/min. In what follows, and for each mouse strain, we first list the viral construct injected or device implanted, then the relevant stereotaxic coordinates are described. Stereotaxic coordinates are shown with respect to *bregma*, according to a standardized atlas of the mouse brain^56^.

<u>Mouse strain **ChAT-ires-Cre**</u>

<u>Injection Location</u> *DMV*

<u>Viral construct</u> AAV1 hSyn FLEx mGFP-2A-Synaptophysin-mRuby (300nL unilateral) Or AAV9-Ef1a-DIO EYFP (300nL each, bilateral)

<u>Coordinates:</u> AP: −7.5mm, ML: ±0.3mm, DV −5.5 ∼-5.3mm.

<u>Mouse strain **Glp1r-ires-Cre x Ai148D**</u>

<u>Injection Location</u> CeM

<u>Viral construct</u> AAV5-hSyn-hM3D(Gq)-mCherry (300nL each, bilateral) Injection Coordinates: AP:-1.0mm, ML: ±2.5mm, DV −4.8mm – 5.2mm.

<u>Mouse strain</u> **<u>B6</u>**

<u>Injection Location</u> *CeM*

<u>Viral construct</u> AAV5-hSyn-hM3D(Gq)-mCherry

<u>Injection Coordinates</u>: AP:-1.0mm, ML: ±2.5mm, DV −4.8mm – 5.2mm. 0.3µL/side.

<u>Mouse strain Glp1r-ires-Cre x Ai148D</u>

<u>Implant Location</u> *CeM*

Stereotaxic electrode implantation

<u>Electrode type: B</u> twisted wire 2-channel electrode (Protech international)

<u>Electrode Coordinates:</u> AP:-1.0mm, ML: ±2.5mm, DV −5.0mm.

### Intravital window for confocal microscopy

For intravital confocal imaging of calcium transients in Brunner’s gland and pancreas (control), we generated Glp1r-ires-Cre×Ai148(TIT2L-GC6f-ICL-tTA2)-D mice (*Cre*-driven GCaMP6f expression) and Glp1r-ires-Cre×Ai148(TIT2L-GC6f-ICL-tTA2)-D×Ai9 mice (*Cre*-driven GCaMP6f compounded with tdTomato expression). These strain nomenclatures were abbreviated as Glp1r[GCamp6] in main text. When 6-week-old, 8-hour food-restricted animals underwent the surgical procedures as below, followed by proximal duodenal/ pancreas abdominal glass window placement for intravital confocal microscopy, as previously described^31,57^. A purse string suture was placed on the incision edge between the rectus abdominis and the abdominal skin. The stomach was exteriorized through the midline incision. The pyloric antrum was loosely stitched to the left rectus abdominis to maximize the field of view over the proximal duodenum and pancreas through the cover glass. Then, the purse string suture was tightened to the groove made on a customized titanium ring (ID ø 14mm, OD ø 32mm, 1mm width, manufactured by Send Cut Send, USA). Drops of Low Toxicity Silicone Adhesive (KWIK-SIL, WPI, Sarasota, FL) were carefully applied to the outer surface of the proximal duodenum and pancreas before being attached to a round coverslip (ø15 mm, Electron Microscopy Sciences, Hatfield, PA) which was previously glued (Ultra Gel Control, Loctite, Hartford, CT) to a second, larger titanium ring (ID ø 14mm, OD ø 46mm, 1mm width manufactured by Send Cut Send, USA). Animals were maintained under continuous 1.5% isoflurane anesthesia. Calcium imaging was performed using a Leica SP8 confocal microscope equipped with a 10X objective lens along with a 0.75X amplification hardware setting. Scanning pixels were 400×400 per frame, and the pinhole was used at maximum value. Upon visualizing GCaMP6f-expressing Brunner’s glands, continuous imaging series were acquired at ∼3.2 frames/sec, FOV=2.25mm^2^, 400X400 frame scanning speed.

### Intravital imaging combined with cholecystokinin infusions

Before intravital window implantation, a MicroRenathane tubing (0.025“, Braintree Scientific, Braintree, MA) was inserted into the peritoneal cavity, guided by a 23-gauge needle, and tightened to the skin with a purse string around the tubing. Cholecystokinin (CCK8) or saline infusates were freshly prepared. After a 12 minutes-long baseline and Saline^ip^ image acquisition period, 0.1mL of 10µg/Kg CCK8 was infused via the i.p. catheter at 0.1mL/min, after which post-injection series were acquired. Imaging was acquired continuously through the pre-, injection, and post-injection periods.

### Intravital imaging combined with cholecystokinin infusions and vagal lesions

To assess the effects of sensory vagal denervation on the above, nodose ganglia were injected with CCK-SAP or Blank-SAP 2 days before intravital window implantation. Subdiaphragmatic vagotomies were performed immediately before intravital window implantation. Intraperitoneal catheterization and CCK/saline injections were as described above.

### Intravital imaging combined with vagus nerve electrical stimulation

Glp1r-ires-Cre×Ai148(TIT2L-GC6f-ICL-tTA2)-D mice were used in these experiments. Before intravital window implantation, the right (dorsal) subdiaphragmatic vagus nerve trunk was separated from the esophagus with a Spinal Cord Hook. Minimally traumatic elastic cuff electrodes (Micro Cuff Sling, 200 µm/3pol/2,5mm/cable entry top, CorTec GmbH) were gently placed under the nerve. The cuff electrode cable ends were soldered to a male miniature pin connector (520200, A-M Systems) and connected to a Grass S48 Pulse Stimulator (A-M Systems). After a 12 minutes-long baseline image acquisition period, electric pulses at 1Hz were triggered via TTL signals at maximum value of 5-20V/minute followed 1-min off rest period. This cycle was repeated multiple times. Then, a 24 minutes-long post simulation image series was recorded.

### Intravital imaging combined with electrical stimulation of the central nucleus of the amygdala

Glp1r-ires-Cre×Ai148(TIT2L-GC6f-ICL-tTA2)-D mice were used in these experiments. As described above, brain stimulation electrodes (Protech international; Tungsten 2-Channel Electrode (B Twisted Wire, MS303T/3-B/SPC) implantation was performed 7 days before intravital window implantation. On the experimental day, both the intravital window and an i.p. catheter were implanted. The stimulation electrode was connected to a Grass S48 Pulse Stimulator (A-M Systems). A cycle of 20 Hz electrical pulses were delivered to central amygdala, triggered via TTL signals at the maximum voltage of 0.5-3V. Experimental design during calcium transients imaging were as follows: *1)* one cycle of electric stimulation (ES1); *2)* i.p. infusion of 200μL of subthreshold dose of CCK (0.05 μg/Kg); *3)* one second electrical stimulation cycle (ES2); *4)* a third cycle of electrical stimulation (ES3); and *5)* one final cycle of electrical stimulation (ES4). Each stimulation cycle lasted one minute, with a 10–20-minute intervals in between. 10 minutes-long baselines before simulation and 10 minutes-long post-stimulations resting periods were scanned for normalization and comparison.

### Gut permeability test

Mice fasted for four hours before 4kDa FITC-dextran, 25 mg/ml (Chrondex #4013), was administered orally (20 mL/kg). Three hours after dosing, mice were anesthetized using isoflurane (induction 5%, maintenance 1.5% isoflurane, 0.7 L/min N2O, 0.3 L/min O2), and retro-orbital blood was collected and stored in heparin-coated tubes. Animals were then euthanized. The samples were centrifuged (4 °C, 7 min, 8000 g) and plasma collected in clear Eppendorf tubes (Fisherbrand™ Premium). Plasma from PBS-administered control mice were used for defining the standard curve. FITC-dextran concentrations in plasma were analyzed in duplicates using a spectrophotometer (SpectraMax 340PC Microplate reader, Molecular Devices, San Jose, CA, USA), under excitation λ=490nm and emission λ=520nm.

### Intraperitoneal Glucose Tolerance Tests

A bolus of a 20% glucose solution (2g glucose/kg) was intraperitoneally injected to 16-hour food restricted animals. Blood glucose levels were measured using the OneTouch Ultra2 Blood Glucose Monitoring System at 0,15,30,60, and 120 minutes post-glucose injections, through placing a small drop of tail blood on an unused test strip (OneTouch Ultra Test Strips). At the end of the experimental session mice were returned to a clean cage with water and food and monitored.

### Oral glucose tolerance tests

Same procedure as above except that mice were orally gavaged with the glucose solution.

### Behavioral tests

#### Food preference test

Mice were single housed for at least a week before testing. During the test, two different types of food pellets, each ∼10g in total, were evenly distributed and left at the cage for 24 hours. Then the amounts remaining of each type of food were weighted, returned to cage, and this was repeated for three days at the exact same time of the day. Daily averages of food intake for each type of food were then calculated. We compared intake between four pairs of food types containing different contents of fat, sugar, protein, and fiber. All different types of food were purchased from Research Diets, Inc. Comparisons were as follows: 1. High-fat (D12451) *vs*. fiber-free high-fat (D13121101); 2. High-fat (D12451) *vs*. regular chow (D12450B); 3. fiber-free high-fat (D13121101) *vs*. fiber-free high-protein (D17120803); 4. fiber-free high-protein (D17120803) vs. high-protein (D22072013).

#### Probiotics preference test

Mice were deprived of water overnight. Two water bottles containing two types of aromatic solutions (banana *vs*. apple, 1:1000) were weighted and provided to the animals for 1hour *ad libitum*. The amount of solution remaining was weighted after the test. Then, for each mouse, the less preferred flavor was identified, and mixed to a probiotics cocktail (300mg/10ml water; probiotics were purchased from RenewLife, lnc). These mixes were provided to condition the mice for three days. In each training day, mice were deprived of water overnight, after which probiotics plus (less preferred) flavor was provided continuously. After three days of conditioning, mice were deprived of water overnight and provided with the two types of flavors (without any probiotics) for one hour. Flavor preferences before and after training were calculated.

#### Open field tests

Open field tests made use of automated video analyzes (EthoVision XT11.5, Noldus). Animals were placed on a novel Plexiglas arena (Med Associates, 25 cm × 30 cm). The total area was divided into a central (8.3 × 10 cm) and a marginal subarea. Immediately above the central subarea, a 150-W lamp was activated to induce natural aversion to this location. Animals were tested once in this arena. The sessions were digitally recorded, and animal activity immediately analyzed using automated video analysis algorithms (EthoVision XT11.5, Noldus).

### Bacterial manipulations

#### Assessment of intestinal and fecal levels of Lactobacilli

B6 mice were administered sterile 0.9% saline (vehicle) or 10μg/Kg CCK solutions, intraperitoneally, daily, for seven days. Small intestinal contents, large intestinal contents, and feces were collected and diluted into sterile ddH_2_O. Small intestinal contents were collected from distal ileum, and large intestinal contents were collected from proximal colon. In both cases, samples were collected from 10mm-long intestinal segments, and fecal samples (0.1mg) were diluted at 1:10000 in sterile water and applied to MRS plates for CFU measurements.

#### Oral gavage of probiotics

These experiments employed the chloramphenicol-resistant strain *Lactobacillus rhamnosus (Hansen) Collins* (ATCC, 27773). B6 mice were administered sterile 0.9% saline solution (vehicle) or 10μg/Kg CCK solutions daily, for 3 days. On the third day, mice were administered around 10^8^ CFUs of *Lactobacillus rhamnosus* by oral gavage. Mice continued to be administered saline or CCK daily for another two days, after which fecal samples were collected. Samples (100µl, from a solution of 1mg feces, diluted at 1:1000 in sterile water) were applied to MRS plates previously treated with 20mg/L chloramphenicol, and clones were counted.

#### Intra-cecum infusions of probiotics

Animals at the time of experiments were 6-week-old. The abdomen and dorsal back skin were shaved and cleaned. A small incision to the dorsal neck, and a midline incision into the abdomen, were made. The cecum was exteriorized through the midline incision. A purse string suture was placed on the distal caecal wall, proximal to the caecal lymphoid patch, into which the tip of MicroRenathane tubing (0.025“, Braintree Scientific, Braintree, MA) was inserted. A purse string was tightened around the tubing, which was then tunneled subcutaneously to the dorsum via a small hole made into the abdominal muscle. A small incision to the dorsum between the shoulder plates was then made to allow for catheter exteriorization. The abdominal incision was then closed with sterile suture. All infusates (Renew Life Extra Care Probiotic, 6mg/mice/day) were freshly prepared in 0.1mL filtered ddH2O and infused at 0.1mL/minute.

#### Bacterial pathogen colonization experiments

These experiments employed either *Staphylococcus xylosus* (ATCC, #29966) or *E. coli* EcAZ-2 (generously provided by Dr. Amir Zarrinpar). In *S. xylosus* experiments, animals lacking Brunner’s glands, or their appropriate controls, were treated with probiotics (intra-cecum infusions, see above) or mucin (in drinking water) for three days. Then, mice were orally gavaged ∼10^8^CFUs bacteria daily, for seven days. Mortality was recorded daily. For *E. coli* EcAZ-2 experiments, animals lacking Brunner’s glands, or their appropriate controls, were treated with probiotics (intra-cecum infusions, see above) or mucin (in drinking water) for three days. Then, mice were orally gavaged ∼10^10^CFUs bacteria one single time. Seven days later, fresh stools were collected, cultured on LB agar plates with 50 µg/mL Kanamycin, and resulting clones counted.

#### Antibiotic treatments and fecal microbiota transplantation

The broad-spectrum antibiotic treatment for mouse models was described before^44^. Specifically, a broad-spectrum antibiotic cocktail (0.25g Vancomycin, 0.25g metronidazole, 0.5g ampicillin, and 0.5g neomycin) were dissolved into 500 mL filtered water. To mask the bitter taste of this antibiotic solution, 5g of artificial sweetener (Splenda) was added. The resulting solutions were filtered with SteriCup, 0.22μm. The experiment used C57BL/6 mice maintained in the Mount Sinai SPF facility. All the C57BL/6 mice were given *ad libitum* access to the antibiotic mixture for 7 consecutive days in drinking water. Antibiotic treated mice were then orally gavaged daily with a freshly prepared solution from fecal samples of Brunner’s gland-lesioned (BGx-DTx) or control (BG-vehicle) mice, at a concentration of 1g/mL, daily, for the next 7 days. Mortality was recorded daily.

#### Central Amygdala chemogenetic stimulation

Stereotaxic viral injections were performed 2 weeks before intravital window implantation. During the experiments, the chemogenetic design drug clozapine-N-oxide (CNO, 5mg/kg) and subthreshold CCK (CCK^Low^, 0.05 µg/kg) were injected intraperitoneally daily. After 7 treatment days, fresh fecal samples were collected and added to MRS plates. For chemogenetic experiments involving EcAZ-2 colonization, mice were treated with CNO and CCK^Low^ for three days, after which they were gavaged once with 10^10^ CFU EcAZ-2. Mice continued to be injected with CNO and CCK^Low^ for additional three days. Seven days after gavage, fresh stools were collected, cultured on MRS plates, and resulting clones counted.

### Analyses of microbiome composition

#### Blood and intestine sample collection

Animals received preoperative analgesia (0.05 mg/Kg Buprenorphine s.c.) and maintained under deep anesthesia with 3% isoflurane until euthanasia. The abdomen was shaved and cleaned with iodine soap and wiped with 70% isopropyl alcohol. The abdominal skin was covered with a sterile surgical drape before a midline incision was made. The intestine was pulled onto the surgical drape to expose the inferior vena cava for blood collection. After cutting the duodenum away from pylori, the complete intestine was pulled out from the abdomen cavity and gently stretched over a sterile surgical drape. Different intestine segments were then identified and separated for further analysis.

#### Fresh fecal sample collection

Freely behaving animals were transferred to a sterile cage without bedding. Around 5 fecal pellets were collected within 15 mins. Feces were stored in −80°C (up to one month) before DNA extraction and sequencing.

#### Bacteria DNA extraction and targeted library preparation

Bacteria DNA from small and large intestinal contents, and from feces, were extracted using the ZymoBIOMICS®-96 MagBead DNA Kit (Zymo Research, Irvine, CA). Bacterial 16S ribosomal RNA gene targeted sequencing was performed using the Quick-16S™ NGS Library Prep Kit (Zymo Research, Irvine, CA). Bacterial 16S primers amplified the V3-V4 region of the 16S rRNA gene. These primers were custom designed by Zymo Research to provide the best coverage of the 16S gene while maintaining high sensitivity. The sequencing library was prepared using an innovative library preparation process in which PCR reactions were performed in real-time PCR machines to control cycles and therefore limit PCR chimera formation. Final PCR products were quantified with qPCR fluorescence readings and pooled together based on equal molarity. The final pooled library was cleaned with the Select-a-Size DNA Clean & Concentrator™ (Zymo Research, Irvine, CA), and then quantified with TapeStation® (Agilent Technologies, Santa Clara, CA) and Qubit® (Thermo Fisher Scientific, Waltham, WA).

#### 16S amplicon sequencing

The final library was sequenced on an Illumina® MiSeq™ with a v3 reagent kit (600 cycles). The sequencing was performed with 10% PhiX spike-in. All the mice including surgical mice were housed in the SPF-level Mount Sinai animal facility before sample collections for bacterial DNA extraction and 16S amplicon sequencing.

#### Bioinformatics Analysis

Unique amplicon sequences variants were inferred from raw reads using the DADA2 pipeline^58^. Potential sequencing errors and chimeric sequences were removed with the Dada2 pipeline. Taxonomy assignment was performed using Uclust from Qiime v.1.9.1 applied to the Zymo Research Database, a 16S database that was internally designed and curated, as reference. Composition visualization, alpha-diversity, and beta-diversity analyses were performed with Qiime v.1.9.1^59^. If applicable, taxonomy with significant abundance among different groups were identified by LEfSe^60^ using default settings. Additional analyses including heatmaps, Taxa2ASV Decomposer, and PCoA plots were performed with Zymo Research proprietary scripts. As a confirmation, and to remove the influence of different reference databases, amplicon sequences were also analyzed using One Codex pipeline and its Targeted Loci Database. Significant enriched species, genera, and families were confirmed through Targeted Loci Database analysis results.

#### Bacterial culturing and identification

100µl dilutions of small or large intestinal contents, or fecal samples, in sterile water were applied to MRS (deMan, Rogosa, Sharpe, Millipore, #110660) plates containing anaerobic cultivation sets (Thermo Scientific, AnaeroPack™) and maintained at 37°C. For the drug-resistant bacterium *Lactobacillus rhamnosus* (ATCC, #27773), the sterile dilutions of bacterial or fecal samples were placed on MRS agar plates with 20mg/L chloramphenicol at 37°C containing anaerobic cultivation sets (Thermo Scientific, AnaeroPack™). For *Staphylococcus xylosus* (ATCC, # 29966), 100µl of sterile dilutions of freeze-dried bacteria or blood samples were plated on brain heart infusion (BHI) plates or 20ml of BHI culture medium at 37°C. For the *E. coli* strain EcAZ-2, sterile dilutions of bacterial or fecal samples were placed on LB agar plates (for clone counting), or incubated in LB medium with 50 µg/mL Kanamycin (for use *in vivo*) at 37°C. For identification of bacterial strains, colonies were washed out from MRS plate using sterile PBS solution and DNA extracted for 16S rRNA sequencing.

#### Vesicles and membrane secretion analysis

Glp1r-ires-Cre×CD63-emGFP^l/s/l^ mice were used to evaluate vesicle translocation and distribution before and after CCK treatment. The tetraspanin CD63 resides in endosomes, lysosomes, secretory vesicles, and plasma membranes and is transported between these compartments. Whenever robust secretion occurs from Brunner’s glands, CD63-emGFP aggregates at the apical membrane region. Double transgenic mice were deprived of food overnight. On the next day, saline, or 10μg/Kg CCK was injected into two groups of mice. After 1h, animals were perfused with filtered saline, followed by 4% paraformaldehyde. Following perfusion, Brunner’s glands segments were placed in 4% paraformaldehyde for 24 hours and then moved to a 20% sucrose solution in 0.02M potassium phosphate buffer (KPBS, pH 7.4) for 48hs. 5μm sections were obtained using a cryotome (Thermo Fisher) and mounted for confocal imaging.

#### Pseudorabies virus tracing

The abdomen of 8-hour food restricted animals was shaved and cleaned. A midline incision was made into the abdomen. Immediately after subdiaphragmatic vagotomy or celiactomy^31^ when appropriate, stomachs were exteriorized through the midline incision, and the pyloric antrum was loosely stitched to the left rectus abdominis, to maximize the view of the proximal duodenum. PRV-Introvent-GFP obtained from CNNV, ^61^ were loaded into a Nanofil^TM^ 36G beveled needle (WPI, Sarasota, FL) and Silflex^TM^ tubing (WPI, Sarasota, FL), connected to a Nanofil^TM^ 10µl syringe (WPI, Sarasota, FL), and mounted on a Pump 11 Elite Nanomite (Harvard Apparatus, Holliston, MA). A 100nL injection was made at 10nL/s into the submucosal layer of duodenum bulb (adjacent to pyloric sphincter) or into the descending part of duodenum in control cases (adjacent to the sphincter of Oddi). The needle tip was carefully guided to avoid mesenteric blood vessels. After completing the infusions, needle was left in place for 5s before removal to ensure full absorption. After removing the loosing suture attached to the stomach, sterile suture was then applied to the skin.

### Histology analysis

#### Histological procedures

Mice were deeply anesthetized with a ketamine/xylazine mix (400mg ketamine + 20mg xylazine/kgBW I.P.). All animals were perfused with filtered saline, followed by 4% paraformaldehyde. Following perfusion, brains or intestinal segments were placed in 4% paraformaldehyde for 24 hours and then moved to a 20% sucrose solution in 0.02M potassium phosphate buffer (KPBS, pH 7.4) for 2 days. Brains were then frozen and cut into four series of 40μm sections with a sliding microtome equipped with a freezing stage. To identify electrode locations, relevant sections were identified and mounted on slides. Sections were then photographed under bright field and fluorescence. For Pseudorabies virus visualization, 3,4 or 5 days after viral injection, mice were perfused, and brains cut at 40 µm.

#### HE staining

All animals were perfused with filtered saline and 4% paraformaldehyde. Intestinal tissues and spleen tissues were post-fixed, embedded in OCT and cryo-sectioned at 5µm. Tissue sections were stained with H&E and imaged using Zeiss Axio Imager using 10X and 40X objectives.

#### PAS staining-based void scores

To preserve Brunner’s glands, goblet cells and intestinal mucus layers for posterior analyses *ex vivo*, immediately after excision, tissues were submerged in Carnoy’s solution (60% ethanol, 30% chloroform and 10% glacial acetic acid) at 4°C for 2hrs and then placed into 100% ethanol, as described before ^17,62^. Fixed tissues were embedded in OCT and cryo-sectioned at 5µm. Tissue was stained with the Periodic Acid Schiff (PAS) Stain Kit (Abcam, #ab150680) and imaged with Zeiss Axio Imager using 10X and 40X objectives.

##### Calculation of void scores using Deep Learning

Unlike goblet cells, Brunner’s glands are most often in an intermediate state between being fully devoid of, and fully filled with, mucus. Accordingly, we generated a “void score” to approximately quantify mucin storage in each gland. We used the automated segmentation algorithm “Segment anything model” (SAM)^63^ to quantify mucin content. When using a rectangle to label an individual gland, the algorithm defines its outer boundary. Now, if the given gland contains a relatively small amount of mucin, the algorithm will define its inner boundary. The quotient of the area associated with the inner boundary over the area associated with the outer boundary was defined as the gland’s “void score”, representing mucin storage in this gland. Whenever the void score was < 0.05, the gland was considered “void”.

#### Immunohistochemical staining

To visualize Fos via immunofluorescence, sections were incubated with a Rabbit Anti-c-Fos antibody (Rabbit Anti-c-Fos ab190289, 1:1000), followed by DyLight™ 649-conjugated goat anti-rabbit (IgG (H+L) DI-1649-1.5, Vector Laboratories, 1:200). Fos expression was analyzed and quantified as follows: Coronal sections collected at ∼160μm intervals were photographed at 10× magnification and montaged with ImageJ to preserve anatomical landmarks. Fos+ neurons in each section were detected and counted using ImageJ and expressed as the cumulative sum of Fos+ neurons within the relevant regions for each animal.

To visualize B cells, cell proliferation, and FDC light zones via immunofluorescence, spleen sections were incubated with Alexa647-conjugated anti-mouse IgD antibody (Biolegend, #405707, 1:200, Clone: 11-26c.2a), Brilliant Violet 421-conjuncted anti-mouse CD21/CD35 antibody (Biolegend, # 123421, 1:200, Clone: 7E9), and Alexa488-conjuncted anti- mouse/human Ki-67 antibody (Biolegend, # 151204, 1:200, Clone: 11F6) at 4°C overnight. Slices were washed three times using KPBS solution and imaged using Zeiss LSM980 Airyscan 2 with a 10X objective.

To visualize goblet cells via immunofluorescence, slices were incubated with Rabbit Anti-MUC2 antibody (Abcam, #ab272692, 1:1000), followed by DyLight™ 649-conjugated affinipure goat anti-rabbit (IgG (H+L) DI-1649-1.5, Vector Laboratories, 1:200). Slices were washed three times using KPBS solution and imaged using Leica STED 3X with a 10X objective.

#### RNAscope

Animals received preoperative analgesia (0.05 mg/Kg Buprenorphine s.c.) and were induced into deep anesthesia with 2% Isoflurane. Animals were transcardially perfused with ice cold Krebs-Henseleit Buffer Modified (K3753-10L, Sigma-Aldrich). Nodose ganglia were exposed and dissected as described previously^55^. The proximal 3mm end of the duodenum (to include the duodenal bulb, within which the glands of Brunner are located) were harvested. All fresh collected tissue were immediately placed into dry ice-cooled cryomolds and embedded with OCT before freezing under −80°C. Tissues were then cryo-sectioned at 10µm and mounted onto Superfrost Plus slides and kept at −80°C. After 1-hour brief fixation in 4% PFA at 4°C, slides were rinsed in PBS and dehydrated with ethanol. The RNAscope procedure was achieved with the RNAscope Multiplex Fluorescent Reagent Kit v2 with TSA Vivid Dyes (Cat No. 323270) and RNAscope® Probes of Mm-Glp1r (418851 RNAscope®), Mm-Cckar-C2 (313751-C2 RNAscope®), 2.5 duplex mouse positive control probe (321651 RNAscope®) and 2-plex negative control probe (320751 RNAscope®).

#### Single nuclei sequencing

Duodenal segments containing Brunner’s glands were rapidly extracted after perfusion with ice-cold 1X modified Krebs-Henseleit buffer (Sigma, K3753-10X1L). Samples were stored in −80°C for up to one month. Fresh-frozen tissues were disaggregated in 1mL of a nuclear extraction buffer^64^ with gentle slicing with a Tungsten Carbide Straight 11.5 cm Fine Scissor (Fine Science Tools, #14558-11) for 10 minutes on ice. Large debris were removed with a 40μm strainer (Falcon, # 352340). The left homogenate was spun at 1000xg for 8 min at 4°C, pellets washed once, and resuspended in 1 ml 1% BSA in PBS. The samples were diluted to 1000 nuclei/μl estimated from counting using Trypan blue staining and performed on a hemocytometer. Nuclei (∼2000 per sample) were loaded onto a 10X Genomics Chromium platform to generate cDNAs carrying cell- and transcript-specific barcodes. Sequencing libraries were constructed using the Chromium Single Cell 3’ Library & Gel Bead Kit 3. Libraries were sequenced on an Illumina NextSeq using paired-end sequencing.

### CyTOF

#### Sample preparation

Animals received preoperative analgesia (0.05 mg/Kg Buprenorphine s.c.) and were induced into deep anesthesia with 2% isoflurane. The abdomen skin was shaved, and a midline incision was made into the abdomen. The spleen and mesenteric lymph nodes (MLNs) were collected. The inferior vena cava was exposed for blood collection. Femurs and humerus were dissected and cut open by removing both epiphyses. Bone marrow (BM) was flushed out and collected. MLNs)/spleen/ BM samples were then placed into 1.5ml tube with RPMI 1640 Media (Gibico, #21870092) + 10% FBS (Thermo Scientific, #A5256701). Tissues were gently sectioned using a Tungsten Carbide Straight 11.5 cm Fine Scissor (Fine Science Tools, #14558-11). Then, a 70μm cell strainer (Fisherbrand, #22-363-548) was placed in a 50-mL tube (Thermo Scientific, #339652), and rinsed with 2mL of RPMI 1640 Media + 10% FBS. Spleen samples were placed in the cell strainer and grinded with a syringe plunger. 10mL of RPMI 1640 Media + 10% FBS was flushed through the strainer. The sample was then centrifuged at 500-600xg for 5min at 4°C, and the supernatant discarded. The cell pellet was resuspended in 2–5mL cold 1x RBC Lysis buffer (Bioscience, #00-4300-54). The suspension was then incubated for 5 minutes at room temperature. The cell suspension was washed with 10–20mL cold RPMI 1640 Media + 10% FBS (ten-fold of RBC lysis buffer). Finally, the cells were 500–600xg centrifuged for 5min at 4°C, and the supernatant discarded. The pellet was resuspended with 2mL of RPMI 1640 Media + 10% FBS and kept on ice until staining. Samples collected from MLNs/Spleen/BM were stained using the mouse antibody panel listed in the Key Resources Table. Cells were stained with an iridium intercalator overnight prior to CyTOF acquisition. Samples were washed twice with water, resuspended in normalization beads (Fluidigm), and filtered through a cell strainer.

#### Sample Processing

Samples were delivered to the processing facility (Mount Sinai HIMC) in fresh media. Cell counts were performed on a Nexcelom Cellaca Automated Cell Counter (Nexcelom Biosciences, Lawrence, MA, USA), and cell viability was measured utilizing Acridine Orange/Propidium Iodide viability staining reagent (Nexcelom). After washing cells in Cell Staining Buffer (CSB) (Fluidigm Corporation) Fc receptor blocking (Biolegend Inc., San Diego, CA, USA) and Rhodium-103 viability staining (Fluidigm) were performed simultaneously with surface markers for 30min at room temperature. Cells were subsequently washed twice in CSB and then fixed and permeabilized using the eBioscience Foxp3 / Transcription Factor Staining Buffer Set. After 30min incubation on ice with the fix/perm buffer, palladium barcoding was performed on each sample utilizing the Fluidigm Cell-ID 20-Plex Pd Barcoding Kit (Fluidigm) following manufacturer’s instructions. After a 30min incubation at room temperature, samples were washed twice in CSB and pooled. Samples were washed in eBioscience Permeabilization wash and intracellular CyTOF staining was performed. Heparin blocking (100 units/mL) was utilized to prevent non-specific binding of intracellular antibodies to eosinophils^65^. After 30min incubation on ice, cells were washed twice in Permeabilization wash and final fixation was performed. Samples were fixed in 1mL 4.4% paraformaldehyde (Electron Microscopy Sciences Inc., Hatfield, PA, USA).125nM Iridium-193 (Fluidigm) and 2nM Osmium tetroxide (EMS) cell labeling was performed simultaneously with sample fixation^66^. After 30min incubation at room temperature, samples were washed twice with CSB and stored in FBS + 10%DMSO at −80°C until acquisition.

#### Data Acquisition/Processing

Prior to data acquisition, samples were washed in Cell Acquisition Solution (Fluidigm) and resuspended at a concentration of 1 million cells per ml in Cell Acquisition Solution containing a 1:20 dilution of EQ Normalization beads (Fluidigm). The samples were then acquired on a Helios Mass Cytometer equipped with a wide bore sample injector at an event rate of <400 events per second. After acquisition, repeat acquisitions of the same sample were concatenated and normalized using the Fluidigm software. The FCS file was further cleaned using the Human Immune Monitoring Center at Mt. Sinai’s internal pipeline. The pipeline removed any aberrant acquisition time-windows of 3 seconds where the cell sampling event rate was too high or too low (2 standard deviations from the mean). EQ normalization beads that were spiked into every acquisition and used for normalization were removed, along with events that had low DNA signal intensity. The pipeline also was used to demultiplex the cleaned and pooled FCS files into constituent single sample files. The cosine similarity of every cell’s Pd barcoding channels to every possible barcode used in a batch was calculated and then was assigned to its highest similarity barcode. Once the cell had been assigned to a sample barcode, the difference between its highest and second highest similarity scores was calculated and used as a signal-to-noise metric. Any cells with low signal-to-noise were flagged as multiplets and removed from that sample. Finally, acquisition multiplets were removed based on the Gaussian parameters Residual and Offset acquired by the Helios mass cytometer.

#### Data analysis

Samples were normalized and debarcoded using the *Premessa* package in R. Cell populations were manually gated using FCS Express (https://www.standardbio.com/products/software/fcs-express, gating strategy, Figure S6A). Cell frequencies were calculated as the fraction of CD45^+^ cells, which were quantified as a percentage of singlet cells. Relative abundances of major cell types such as total B cells, T cells, immature B cells, mature B cells, CD4 T cells, CD8 T cells, dendrite cells, macrophages and monocytes were calculated.

#### Olink proteomics analysis

Blood collection from inferior vena cava was described as above. Serum was separated from centrifuged blood. Olink data were generated from serum samples submitted to Olink Proteomics for analysis using the Olink Target 96 Mouse Exploratory panel assay of 92 analytes (Supplementary Table 2). Out of 92 proteins, 69 were detected in >75% of samples and used in analysis. Data are presented as normalized protein expression values (NPX, Olink Proteomics arbitrary unit on log2 scale). See Supplemental Table 2 for Olink missing values.

#### In vivo ultrasound studies

Mice were maintained anesthetized with inhaled 1.5% isoflurane. Animals were kept in supine position on a 37°C heating platform, and heart and breath rates were monitored throughout. All data videos were recorded under approximately the same anesthesia, heart, and breathing parameters. To measure the sizes of stomach under different experimental conditions, videos were recorded capturing the whole stomach along both the esophagus-directed and duodenum-directed axis. Images were analyzed with Visual Sonics Vevo 2100 high-resolution ultrasound imaging system. To track and evaluate the movement of the duodenum, the analysis considered breathing periods to avoid movement confounds.

#### Quantification and Statistical Analyses

The exact value of all N (always number of animals), df, T/F, and p values are reported in Supplemental Table 3 containing details on the statistics in the interest of space in text/legends. Figure legends report p values whenever relevant.

Data analyses, excluding electromyogram/electrophysiological/calcium imaging data, were performed using SPSS (v.21.0, IBM Predictive Software), Ethovision XT 11.5 (Noldus), GraphPad Prism 7 (GraphPad) and Matlab (v.17b, MathWorks). Before experiments, groups of animals assigned to the different experimental conditions were formed by naïve littermates, so that no randomization or other *a priori* criteria were adopted for group assignments. Experimental manipulations were analyzed according to within-subject repeated-measures designs whenever appropriate. Order of experimental conditions was randomly assigned across subjects. Samples sizes were chosen based on our previous studies employing similar intravital imaging, electrophysiological, behavioral, and neuronal ablation approaches. Samples sizes adopted in our current study were sufficient for detecting robust effect sizes while complying with guidelines from the Institutional Animal Care and Use Committee enforcing minimal and ethical animal usage. Experimenters were not blind to experimental conditions. Only animals carrying signs of distress/infection/bleeding/anorexia after the surgical procedures were excluded. Data from all animals used in the experiments were included in the final analyses and plots.

#### Mouse behavioral data analysis

For all behavioral studies, including those resulting from cell type-targeted lesions, or chemogenetic experiments, analyses made use of standard linear models (Pearson correlation), as well as one- or two-way (repeated measures) ANOVAs and post-hoc t-tests tests whenever relevant, for correcting for multiple comparisons. All data were reported as mean±SEM. In all cases sample sizes (N) denote number of animals used. All p-values associated with the t-tests performed correspond to two-tailed tests, and all post-hoc tests were corrected for multiple comparisons by employing Bonferroni correction. To assess potentially spurious results associated with non-normality, all significant effects were confirmed by rerunning the tests using the appropriate non-parametric test. All data are individually plotted (Prism 7, GraphPad), and the corresponding bar plot of the precision measures (mean ± SEM) were overlaid on the figure. The exact value of all N, df, T/F, and p values are reported in Supplemental Table 3 containing details on the statistics. Effects were considered statistically significant whenever the corresponding statistic was associated with a p-value (Bonferroni-corrected when appropriate) strictly less than 0.05.

#### Calcium imaging data analysis

The raw data from calcium transients were first denoised using Noise2Void, a self-supervised deep-learning method for spatiotemporal enhancement of calcium imaging data^67^. Unlike Noise2Noise-based DeepCAD^68^, Noise2Void does not need paired images. Specifically, the first 2000 frames of the whole stack were transferred into patches with a (32, 64, 64) format, 70% of which were used for training and 30% for testing. Overall number of training epochs was ∼150- 200, allowing for obtaining the best model. The learning rate started at 0.0004; if the loss-of-function value did not decrease within 10 epochs, a new learning rate was defined as 0.5 x current rate. The whole stack was then transferred to a denoised dataset using the best model. The data were then motion-corrected using NoRMCorre^69^. We then generated the maximum intensity projection of the time-lapse images and utilized Cellpose, a deep learning-based segmentation method or a Mask R-CNN plus PointRend^70^, to segment individual glands. The output files corresponding to each region of interest (ROIs) of individual glands were stored as an ImageJ ROI file. Using the motion-corrected imaging data and segmentation ROI files, the mean values of both ROI GCaMP signals (ROI-Signal_Mean_) and of the surrounding background (Background-Signal_Mean_) for each individual gland were then calculated, following the strategy of the CaImAn software package^69^. The calcium signal of each gland was then defined as ROI- Signal_Mean_ – Background-Signal_Mean_. We then denoised each gland-associated raw data using a Gaussian filter. Next, for each gland, calcium signal z-scores were computed using the mean values and standard deviations obtained from 6min-long baseline periods without any stimulation. The *Python*-based packages used for data analysis included NumPy (version 1.22.0), SciPy (version 1.7.1), Pandas (version 1.4.1), Matplotlib (version 3.4.3), and Seaborn (version 0.11.2).

#### Single nuclei sequencing data analysis

The unique molecular identifier (UMI) count matrices obtained from Cell Ranger (v. 7.1.0) processing for cell barcode aggregation, genome alignment, and unique molecular identifier transcript quantification was cleaned from cell free mRNA contamination with SoupX (v. 1.6.1) package^71^ for R programming environment. The filtered matrices were analyzed with the Scanpy (v. 1.9.1) package^72^ for a Python programming environment. For downstream analysis, low-quality cells were removed based on the following criteria. (i) Cells with relative high percentage of mitochondrial genes were removed. (ii) Cells with fewer than 120 unique genes were removed. (iii) Cells with doublet scores, which were estimated using Scrublet package higher than 0.4, were removed. Remaining cells were normalized to 10,000 reads per cell for downstream comparison. The top 3,000 variable genes were preselected and used for dimensionality reduction by PCA. Next, cell clusters were detected by constructing a shared nearest neighbor graph. The detected cell clusters were visualized by performing the PAGA combined UMAP algorithm. After fixing the clusters, marker genes were identified using Wilcoxon rank test. Panels of *dotplots* and *umaps* were created with Scanpy and matplotlib packages. The cell ontology classes were annotated using PanglaoDB (https://panglaodb.se/index.html).

## Notes

### Competing Interest Statement

The authors have declared no competing interest.

### Summary of Updates

I have a mistake in abstract part. We lost one sentence for the first submission. So we chagned it.

