## Supplementary material for "An Amygdalar-Vagal-Glandular Circuit Controls the Intestinal Microbiome": Supplentary Figures with legends

### Figure S1

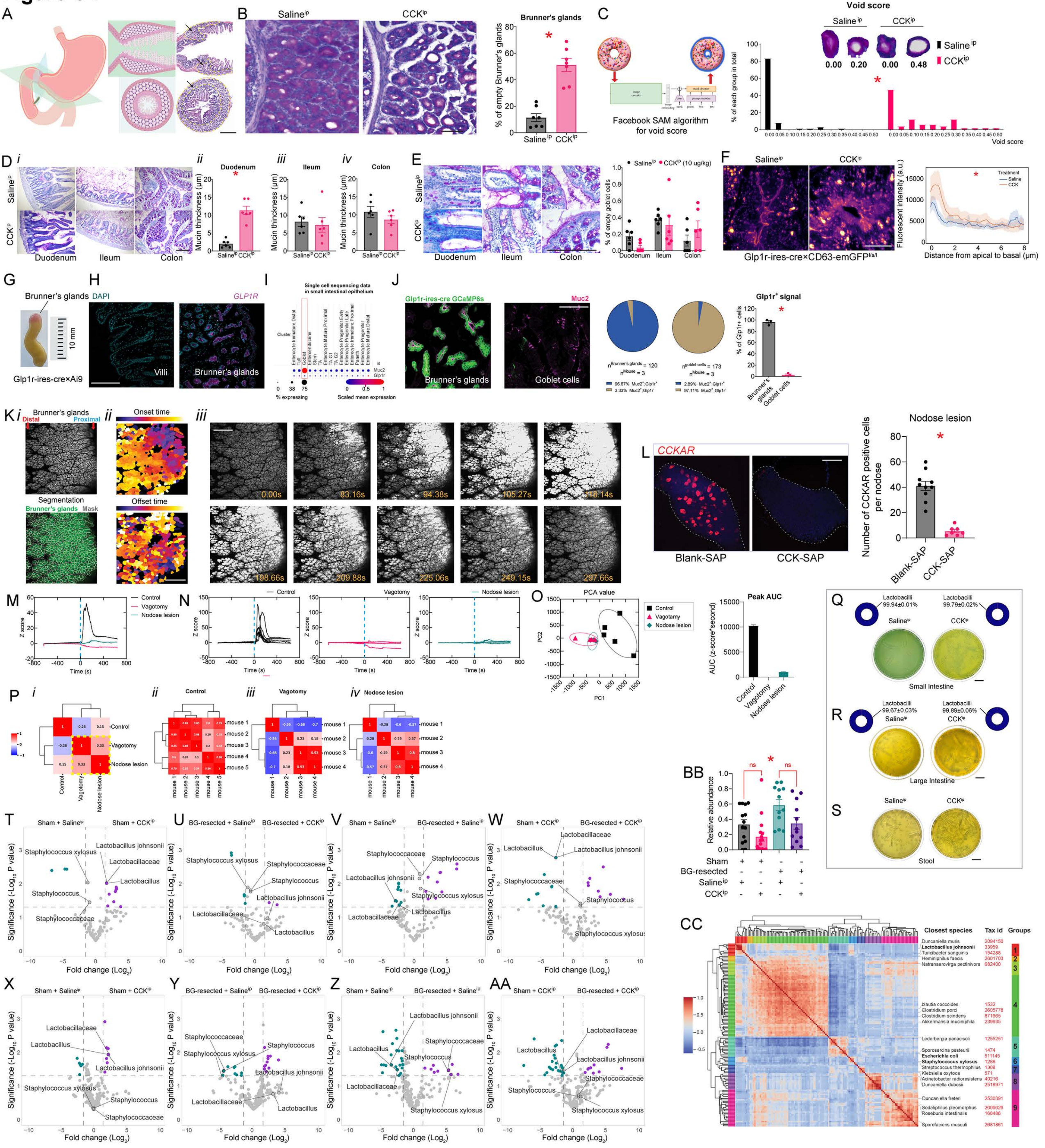

A
B
Saline^ip
CCK^ip
Brunner's glands
% of empty Brunner's glands
C
Facebook SAM algorithm for void score
Void score
% of each group in total
D
Duodenum
Ileum
Colon
Mucin thickness (μm)
E
% of empty goblet cells
F
Glp1r-ires-cre×CD63-emGFP
Fluorescent intensity (a.u.)
Distance from apical to basal (μm)
G
Brunner's glands
10 mm
Glp1r-ires-cre×Ai9
H
DAPI
GLP1R
Villi
I
Single cell sequencing data in small intestinal epithelium
J
Glp1r-ires-cre GCaMP6s
Muc2
Goblet cells
Glp1r+ signal
% of Glp1r+ cells
K
Distal
Proximal
Segmentation
Onset time
Offset time
L
CCKAR
Blank-SAP
CCK-SAP
Nodose lesion
Number of CCKAR positive cells per nodose
M
Control
Vagotomy
Nodose lesion
Z score
Time (s)
N
O
PCA value
Peak AUC
AUC (z-score*second)
P
Q
Lactobacilli
99.94±0.01%
99.79±0.02%
Small Intestine
R
99.67±0.03%
99.89±0.06%
Large Intestine
S
Stool
BB
Relative abundance
Sham
BG-resected
T
U
V
W
X
Y
Z
AA
Significance (-Log10 P value)
Fold change (Log2)
Staphylococcus xylosus
Lactobacillus johnsonii
Lactobacillaceae
Lactobacillus
Staphylococcus
Staphylococcaceae
CC
Closest species
Tax id
Groups

**Figure S1. Related to Figure 1**

**A.** The glands of Brunner (BG) are located within the submucosa of the proximal duodenum. Scale bar=1mm. **B.** 1h after an intraperitoneal CCK injection ($CCK^{ip}$, 10µg/Kg), PAS staining reveals robust BG voiding and mucin secretion: Percentage of empty glands (void scores >0.05), 2-sample t-test, ∗p<0.001. Bar=100µm. **C.** Left: The Deep Learning model "Segment-Anything" was used to calculate the "void score" of a given gland by adaptively detecting both the outside boundary and the internal (void) region (See STAR Methods for details). Right: Histogram showing the distribution of BG void scores in saline *vs*. CCK injected mice. A total of 110 glands from 7 mice in each group. 2-sample t-test, ∗p< 0.0001. **D.** (***i***) Mucin thickness as assessed with PAS-staining of samples from duodenum, ileum, and colon of vehicle- *vs.* $CCK^{ip}$-treated mice. Bar=200µm. Mucin thickness on the surface of intestinal villi after $CCK^{ip}$ *vs.* vehicle: Duodenum: 2-sample t-test, ∗p<0.001 (***ii***); for Ileum (***iii***) and Colon (***iv***), p>0.25. **E.** Left: PAS staining of duodenum, ileum, and colon epithelia in saline- *vs.* CCK-treated mice. Right: percentage of void goblet cells in duodenum, ileum, and colon epithelia in vehicle- *vs.* $CCK^{ip}$-treated mice, two-way mixed RM-ANOVA, main effects of injection, p=0.7742, Injection x segment, p=0.1473. Bar= 200µm. **F.** Glp1r-ires-cre×CD63-$emGFP^{l/s/l}$ mice were injected with vehicle or $CCK^{ip}$ 1h before perfusion. Left: Differential distribution of CD63-emGFP fluorescent intensities indicate greater translocation of vesicles to membrane after $CCK^{ip}$ *vs.* $Saline^{ip}$ injections. Right: fluorescent intensity distribution from apical membrane to ~8µm towards basal membrane, ANOVA effects of injection. ∗p<0.0001. Bar=10µm. **G.** In Glp1r-ires-cre×Ai9 mice, Brunner's glands were easily visualized in the proximal duodenum, spanning 2mm length x 3-5mm width regions. Bar=10mm. **H.** *In situ* hybridization for *Glp1r* transcripts in BG and overlaying duodenal villi. Glp1r-reactive cells exclusively located in BG. Bar=200µm. **I.** We confirmed the previous finding using public transcriptome databases. Single cell sequencing data from small intestine epithelium shows that goblet cells are Muc2-positive and Glp1r-negative (GEO; GSE92332, Single Cell Portal). **J.** Confocal imaging of duodenal samples from Glp1r [GCamp6] mice. Around 96.67% of Glp1r positive cells were localized to Brunner's glands, and only 2.89% to Muc2+ (goblet) cells. BG exhibited strong Cre-dependent GCaMP6 signals. In contrast, GCaMP6 signal was absent from mucous (Muc2+) goblet cells: paired t-test, ∗p=0.0013. Bar=100µm. **K.** (***i***): Deep Learning algorithms were used for segmenting individual glands for further analyses (segmented glands in green on right panel).(***ii***): glands closer to the proximal end of duodenum display early onset and protracted deactivation of calcium transients. Bar=200µm. (***iii***) Timelapse of Brunner's glands calcium transients from onset (upper) to deactivation (lower). **L.** CCK-SAP injected into nodose ganglia to ablate gut-innervating vagal sensory neurons. Controls injected with Blank-SAP. CCKAR-positive neurons were completely ablated after CCK-SAP (counts from *in situ* hybridization), 2-sample t-test, ∗p<0.0001. Bar=100µm. **M.** Average calcium transient Z-scores before *vs.* after CCK infusions in control, vagotomized, and nodose-lesioned (CCK-SAP) mice. Blue line=$CCK^{ip}$. **N.** As in ***M*****,** this time showing data from all individual mice: Sham intact (Left), vagotomized (Middle) and CCK-SAP (Right). Blue lines= $CCK^{ip}$. **O.** Left: Principal Component nalyses (PCA) applied to z-scores from the three groups. Values from vagotomized and nodose-lesioned mice clustered together but were fully separate from the intact mice cluster. Right: The corresponding total AUC values were calculated for the three groups. **P.** (*i*) Between-group Pearson correlation matrices based on the average calcium transient values from control, vagotomized, and CCK-SAP mice. (*ii-iv*): Same as (*i*), but for individual mice in control (ii), vagotomized (iii), and CCK-SAP (iv) groups. **Q-R.** Confirmation that clones from MRS culture plates are *Lactobacilli*. Representative examples of MRS-based culture plates of small intestine, large intestine, and fecal contents in Saline$^{ip}$ and CCK$^{ip}$ conditions. Upper percentage graph in **Q**: Two MRS plates each from vehicle- *vs.* CCK-treated mice. Small intestine contents were sequenced using 16S protocols, and >99.7% of the clones were identified as *Lactobacilli*. **R**: Same as in ***Q*** but for large intestine: >99.6 of the clones were identified as *Lactobacilli.* **S.** MRS plates from fecal samples. Bar=20 mm. **T.** Volcano plot contrasting relative abundances of bacteria after 16S sequencing of small intestine contents in sham+Saline$^{ip}$ *vs*. sham+CCK$^{ip}$ injection. Cyan represents downregulated, and purple upregulated, species. **U.** As in ***T*** but for BG-resected+Saline$^{ip}$ *vs*. BG-resected+CCK$^{ip}$ groups. **V.** As in ***T-U*** but for sham+Saline$^{ip}$ *vs*. BG-resected+Saline$^{ip}$ groups. **W.** As in ***U-V*** but for sham+CCK$^{ip}$ *vs.* BG-resected+CCK$^{ip}$ groups. **X.** As in ***T*** but for large intestine. **Y.** As in ***U*** but for large intestine. **Z.** As in ***V*** but for large intestine. **AA.** As in ***W*** but for large intestine. **BB.** Same as *Figure 1O* but for *Staphylococci* instead of *Lactobacilli* (small intestine contents). Surgery**,** p<0.0009. **CC.** Pearson correlation matrices and K-mean clustering were calculated across the different bacterial strains associated with significant LEfSe values. In total, 9 groups formed well-defined clusters.

### Figure S2

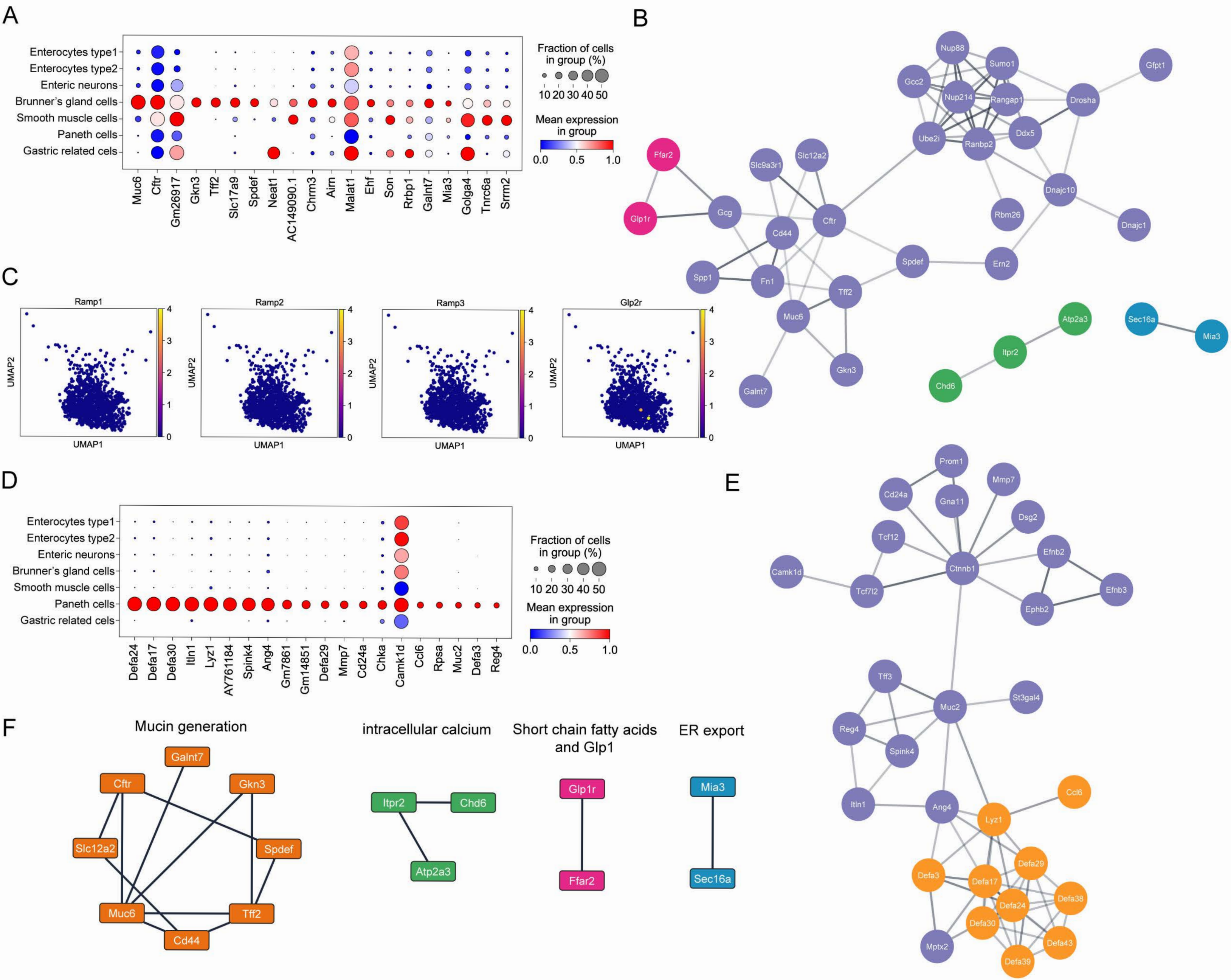

A
Enterocytes type1
Enterocytes type2
Enteric neurons
Brunner's gland cells
Smooth muscle cells
Paneth cells
Gastric related cels
Muc6
Cftr
Gm26917
Gkn3
Tff2
Slc17a9
Spdef
Neat1
AC149090.1
Chrm3
Airn
Malat1
Ehf
Son
Rrbp1
Galnt7
Mia3
Golga4
Tnrc6a
Srrm2
Fraction of cells in group (%)
10 20 30 40 50
Mean expression in group
0.0 0.5 1.0
B
Nup88
Sumo1
Gfpt1
Gcc2
Nup214
Rangap1
Drosha
Ube2i
Ranbp2
Ddx5
Ffar2
Slc9a3r1
Slc12a2
Dnajc10
Glp1r
Gcg
Cftr
Rbm26
Dnajc1
Cd44
Spdef
Ern2
Spp1
Fn1
Tff2
Muc6
Atp2a3
Sec16a
Mia3
Itpr2
Gkn3
Chd6
Galnt7
C
Ramp1
Ramp2
Ramp3
Glp2r
UMAP1
UMAP2
0 1 2 3 4
D
Enterocytes type1
Enterocytes type2
Enteric neurons
Brunner's gland cells
Smooth muscle cells
Paneth cells
Gastric related cels
Defa24
Defa17
Defa30
Itln1
Lyz1
AY761184
Spink4
Ang4
Gm7861
Gm14851
Defa29
Mmp7
Cd24a
Chka
Camk1d
Ccl6
Rpsa
Muc2
Defa3
Reg4
Fraction of cells in group (%)
10 20 30 40 50
Mean expression in group
0.0 0.5 1.0
E
Prom1
Cd24a
Mmp7
Gna11
Dsg2
Tcf12
Efnb2
Camk1d
Ctnnb1
Tcf7l2
Efnb3
Ephb2
Tff3
St3gal4
Muc2
Reg4
Spink4
Ccl6
Itln1
Ang4
Lyz1
Defa29
Defa3
Defa17
Defa38
Defa24
Defa30
Defa43
Mptx2
Defa39
F
Mucin generation
Galnt7
Cftr
Gkn3
Slc12a2
Spdef
Muc6
Tff2
Cd44
intracellular calcium
Itpr2
Chd6
Atp2a3
Short chain fatty acids and Glp1
Glp1r
Ffar2
ER export
Mia3
Sec16a

**Figure S2. Related to Figure 2.**

**A.** Topmost enriched genes in Brunner’s glands (cluster #3). **B.** Protein interaction networks for the topmost enriched genes in Brunner’s glands. **C.** Additional receptor genes not expressd in Brunner’s glands. **D.** Topmost enriched genes in Paneth cells (cluster #5). **E.** Protein interaction networks for the topmost enriched genes in Paneth cells. **F**. Protein-protein interaction networks in Brunner's glands, based on the top 40 most enriched genes.

### Figure S3

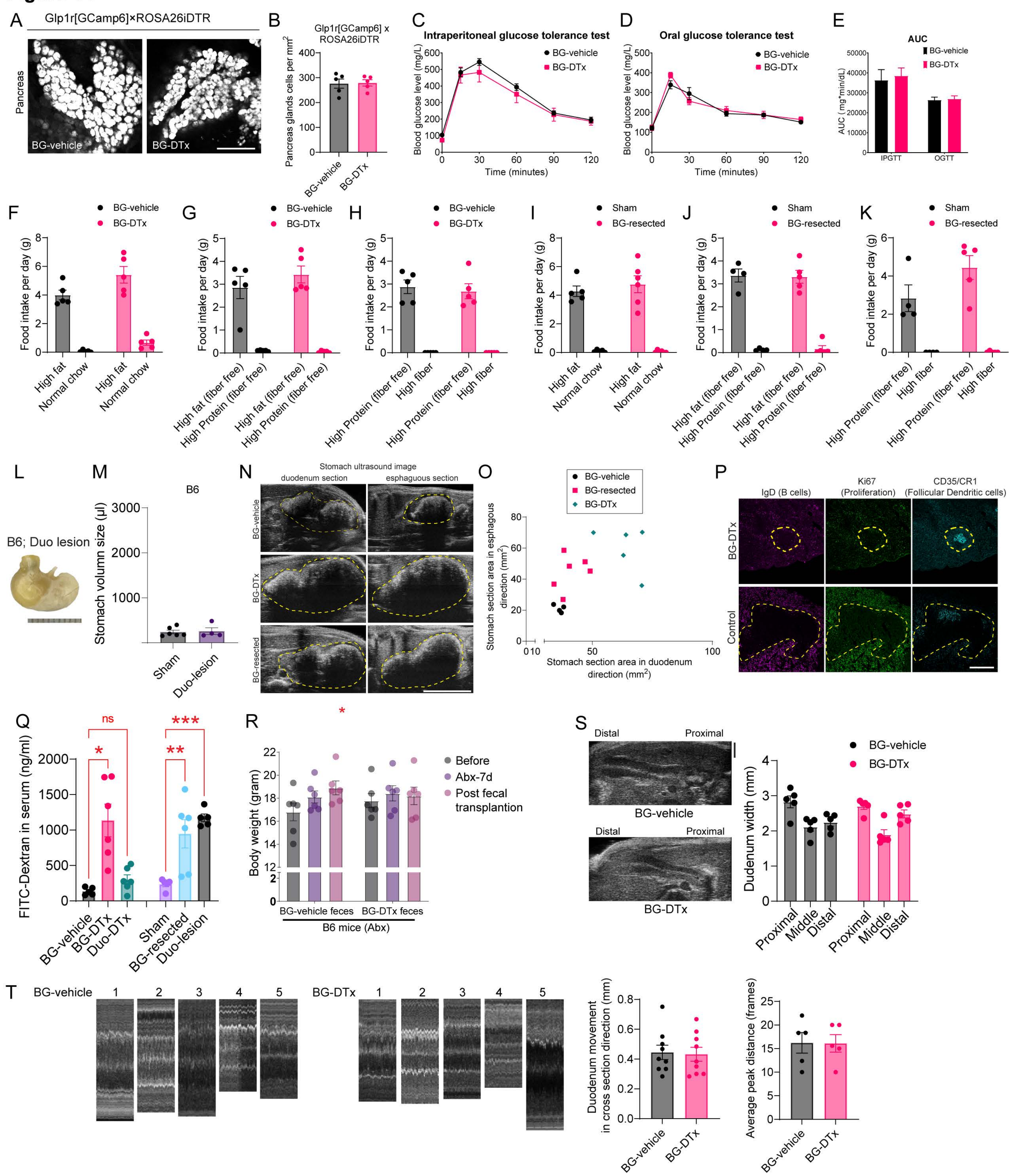

A
Glp1r[GCamp6]×ROSA26iDTR
Pancreas
BG-vehicle
BG-DTx
B
Glp1r[GCamp6] x ROSA26iDTR
Pancreas glands cells per mm²
BG-vehicle
BG-DTx
C
Intraperitoneal glucose tolerance test
Blood glucose level (mg/L)
Time (minutes)
BG-vehicle
BG-DTx
D
Oral glucose tolerance test
Blood glucose level (mg/L)
Time (minutes)
BG-vehicle
BG-DTx
E
AUC
AUC (mg*min/dL)
IPGTT
OGTT
BG-vehicle
BG-DTx
F
Food intake per day (g)
High fat
Normal chow
BG-vehicle
BG-DTx
G
Food intake per day (g)
High fat (fiber free)
High Protein (fiber free)
BG-vehicle
BG-DTx
H
Food intake per day (g)
High Protein (fiber free)
High fiber
BG-vehicle
BG-DTx
I
Food intake per day (g)
High fat
Normal chow
Sham
BG-resected
J
Food intake per day (g)
High fat (fiber free)
High Protein (fiber free)
Sham
BG-resected
K
Food intake per day (g)
High Protein (fiber free)
High fiber
Sham
BG-resected
L
B6; Duo lesion
M
B6
Stomach volumn size (μl)
Sham
Duo-lesion
N
Stomach ultrasound image
duodenum section
esphaguous section
BG-vehicle
BG-DTx
BG-resected
O
BG-vehicle
BG-resected
BG-DTx
Stomach section area in esphagous direction (mm²)
Stomach section area in duodenum direction (mm²)
P
IgD (B cells)
Ki67 (Proliferation)
CD35/CR1 (Follicular Dendritic cells)
BG-DTx
Control
Q
FITC-Dextran in serum (ng/ml)
ns
*
***
**
BG-vehicle
BG-DTx
Duo-DTx
Sham
BG-resected
Duo-lesion
R
*
Body weight (gram)
Before
Abx-7d
Post fecal transplantion
BG-vehicle feces
BG-DTx feces
B6 mice (Abx)
S
Distal
Proximal
BG-vehicle
BG-DTx
Dudenum width (mm)
Proximal
Middle
Distal
T
BG-vehicle
BG-DTx
Duodenum movement in cross section direction (mm)
Average peak distance (frames)

**Figure S3. Related to Figure 3.**

**A-B.** Glp1r[GCamp6]×ROSA26iDTR triple transgenic mice were used to exclude potential pancreatic lesions after DTx microinjections into BG. **A:** Confocal imaging of gland cells in pancreas head. **B:** $Glp1r^+$ pancreas glands cells/$mm^2$ in BG-vehicle *vs.* BG-DTx, 2-sample t-test, p=0.9066. Bar=200 µm. **C-D.** DTx microinjections into BG did not alter blood glucose homeostasis. **C:** Intraperitoneal glucose tolerance test, IPGTT, in BG-vehicle *vs.* BG-DTx mice, 2-way mixed RM-ANOVA, Surgery, p=0.4436, Surgery x sampling time, p=0.8065. **D:** Oral glucose tolerance test, OGTT, Surgery, p=0.6663, Surgery x sampling time, p=0.0942. Bar=200 µm. **E.** AUC values for IPGTT and OGTT. No significant between-group effects detected. Unit is mg∗min/dL. **F.** Daily food preferences for high-fat *vs*. chow pellets in BG-vehicle and BG-DTx mice. No significant between-group effects detected, Two-way mixed RM-ANOVA p=0.5069. **G.** As in ***F*** but for high fat (fiber free) *vs.* high protein (fiber free) pellets, p=0.3487. **H.** As in ***F-G*** but for high protein (fiber free) *vs.* fiber-rich pellets, p=0.6747. **I.** As in ***F*** but for sham *vs.* BG-resected mice, p=0.4965. **J.** As in ***G*** but for sham *vs.* BG-resected mice, p=0.7637. **K.** As in ***H*** but for sham *vs.* BG-resected mice, p=0.1327. **L-M.** In control experiments, resection of duodenal layers underneath Brunner's glands (Duo-lesion) failed to cause gastric distension. **L:** Representative example of the aspect of the stomach in Duo-lesion mice. **M:** Duo-lesion *vs*. Sham 2-sample t-test, p=0.8177. Sham data as in Figure 3J. Bar=10mm. **N-O.** Ultrasound imaging was used to confirm gastric distension *in vivo*. Intact, BG-DTx, and BG-resection mice on a Glp1r-ires-cre×ROSA26-iDTR background were anesthetized under isoflurane. Stomach section sizes in both the duodenal and esophageal orientations were measured. **N**: Representative examples from control, BG-DTx, and BG-resection mice. **O**: BG-DTx stomachs distended in both directions while BG-resection stomachs distended preferentially in the esophageal direction. Bar=5 mm. **P.** Spleen sections stained against IgD, Ki67, CD35/CR1 antibodies. Bar=200 µm. **Q.** Genetic and surgical BG ablation induces "leaky gut". Intestinal permeability was assessed by measuring FITC-dextran in systemic circulation 3hs after intraluminal injection of 4kDa FITC-dextran in BG-vehicle, BG-DTx, and Duo-DTx mice (Glp1r-ires-cre×ROSA26-iDTR background) as well as sham, BG-resection, Duo-lesion mice (B6 background). One-way ANOVA, surgery, p<0.0001. Bonferroni post-hoc planned comparisons, BG-vehicle *vs*. BG-DTx, ∗p=0.0005, and BG-vehicle *vs*. Duo-DTx, p>0.9999. Also, Sham vs. BG-resection, ∗∗p=0.0237, but Sham vs. Duo-lesion, ∗∗∗p=0.0022. Thus, although less impactful than BG lesions, duodenal lesions also increased permeability. **R.** Body weights of B6 mice after antibiotic cocktail (Abx) treatment and BG-DTx fecal transplant. Two-way mixed RM-ANOVA, group, p=0.831; group×time, ∗p=0.011. **S.** BG-vehicle and BG-DTx mice show similar sizes of proximal, middle, and distal duodenal segments. Two-way mixed RM-ANOVA, p=0.1951. **T.** No effects of lesions on duodenal movement patterns. Left: Kymographs of duodenum movements in BG-vehicle and BG-DTx mice. Middle: duodenal cross-sectional movement, 2-sample t-test, p=0.8486. Right: Peak distances of duodenum movement (frames), 2-sample t-test, p=0.9758.

### Figure S4

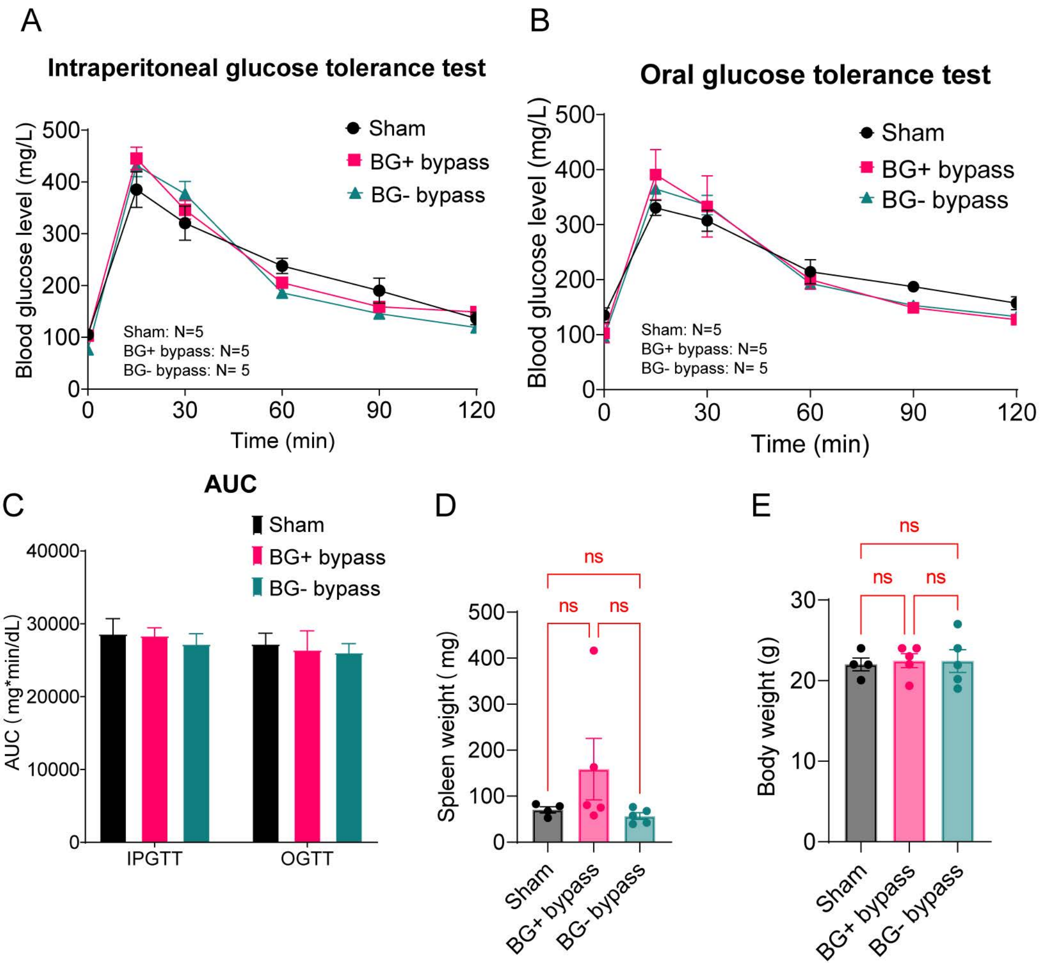
A
Intraperitoneal glucose tolerance test
Blood glucose level (mg/L)
Time (min)
Sham
BG+ bypass
BG- bypass
Sham: N=5
BG+ bypass: N=5
BG- bypass: N= 5
B
Oral glucose tolerance test
Blood glucose level (mg/L)
Time (min)
Sham
BG+ bypass
BG- bypass
Sham: N=5
BG+ bypass: N=5
BG- bypass: N= 5
C
AUC
AUC ( mg*min/dL)
Sham
BG+ bypass
BG- bypass
IPGTT
OGTT
D
Spleen weight ( mg)
ns
Sham
BG+ bypass
BG- bypass
E
Body weight (g)
ns
Sham
BG+ bypass
BG- bypass

**Figure S4. Related to Figure 4.**

**A-C.** Blood glucose tolerance tests in Sham, BG+ and BG- mice. No effects of surgery type detected. **A:** Intraperitoneal glucose tolerance test, IPGTT: 2-way mixed RM-ANOVA, surgery, p=0.679; surgery x sampling time, p=0.007. **B:** Oral glucose tolerance test, OGTT, surgery F[2,12]=0.104, p=0.902; surgery × sampling time, p=0.33. **C.** AUC values for IPGTT and OGTT. Average AUC values for IPGTTs were 28,656±2,047 for Sham, 28,404±1061 for BG+, 27,258±1,384 for BG-. Average AUC values for OGTTs were 27,276±1,436 for Sham, 26,480±2,544 for BG+, and 26,100±1,177 for $BG^{-}$. Units= mg/dL/min. All Bonferroni post hoc tests n.s.. **D.** No effects observed on spleen weights for Sham, BG+ and BG- mice. One-way ANOVA, surgery, p=0.2087. All Bonferroni post hoc tests n.s.. **E.** No effects on body weights for Sham, BG+ and BG- mice. One-way ANOVA, surgery, p=0.9534. All Bonferroni post hoc tests n.s..

### Figure S5

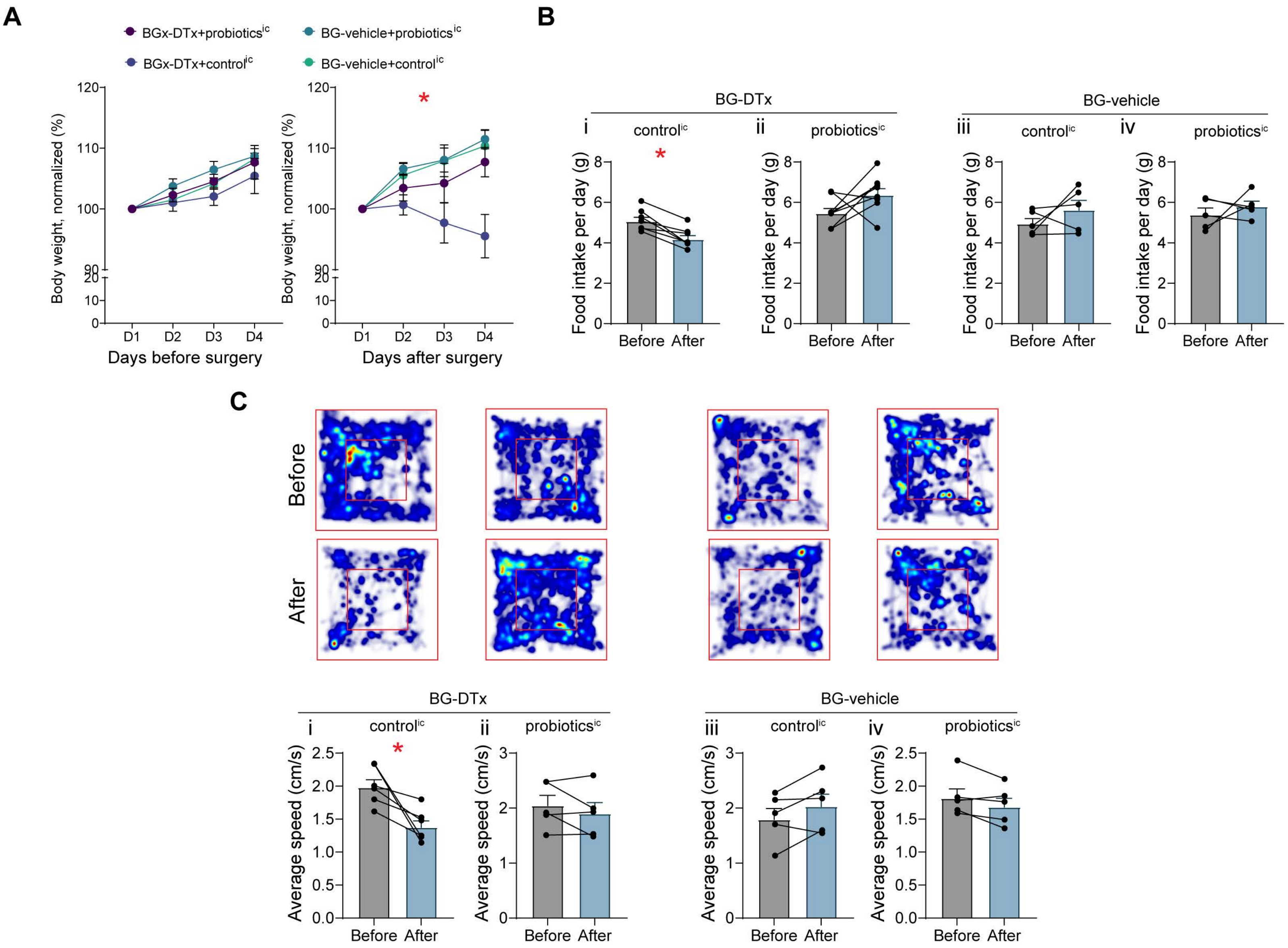

A
BGx-DTx+probiotics^ic
BG-vehicle+probiotics^ic
BGx-DTx+control^ic
BG-vehicle+control^ic
Body weight, normalized (%)
Days before surgery
Days after surgery
D1 D2 D3 D4
B
BG-DTx
BG-vehicle
i control^ic
ii probiotics^ic
iii control^ic
iv probiotics^ic
Food intake per day (g)
Before After
C
Before
After
BG-DTx
BG-vehicle
i control^ic
ii probiotics^ic
iii control^ic
iv probiotics^ic
Average speed (cm/s)
Before After

**Figure S5. Related to Figure 5.**

**A.** Body weights before (left) and after (right) BG ablation followed by intra-cecal probiotic administration, 2-way mixed RM-ANOVA, before surgery, intervention effect, p=0.3932; after surgery: p=0.0144. **B.** Daily food intake before and after surgery for vehicle and probiotics treatments. (i) BG-DTx+control$^{ic}$: paired t-test, ∗p=0.0021. (ii) BG-DTx+probiotics$^{ic}$: p=0.1081. (iii) BG-vehicle+Control$^{ic}$, p=0.3122, (iv) BG-vehicle+probiotics$^{ic}$: p=0.4892. **C.** Open-field tests for vehicle and probiotics treatments. Upper: heatmaps of mice in the 4 treatment groups before and after surgery. Lower: (i) BG-DTx+control$^{ic}$: paired t-test, ∗p=0.0190. (ii) BG-DTx+probiotics$^{ic}$: p=0.3276. (iii) BG-vehicle+control$^{ic}$: p=0.1382, (iv) BG-Vehicle+probiotics$^{ic}$: p=0.1208.

### Figure S6

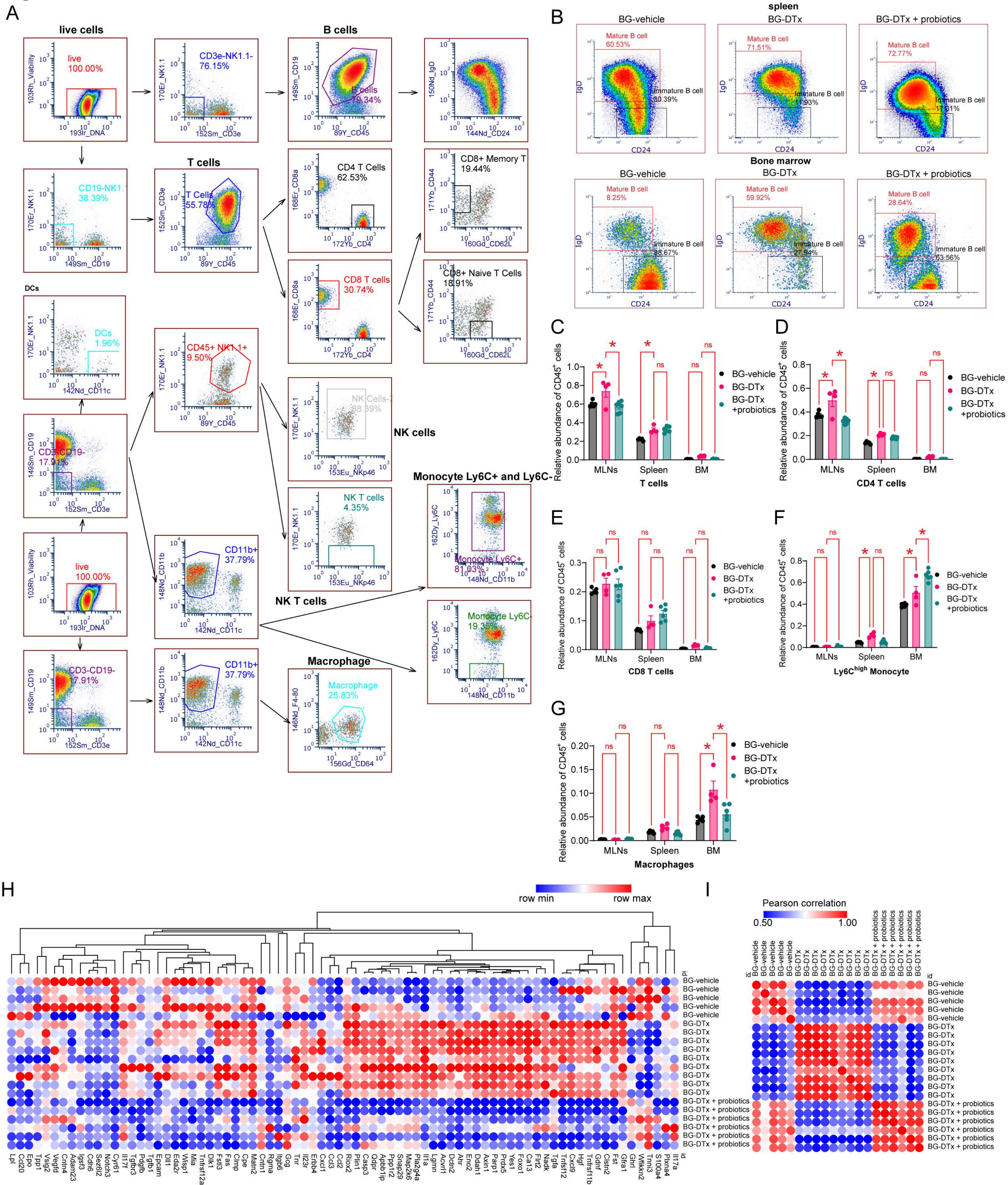
A
live cells
live
100.00%
103Rh_Viability
193Ir_DNA
CD3e-NK1.1-
76.15%
170Er_NK1.1
152Sm_CD3e
B cells
149Sm_CD19
89Y_CD45
B cells
79.34%
150Nd_IgD
144Nd_CD24
T cells
CD19-NK1.1-
38.39%
149Sm_CD19
T Cells
55.78%
152Sm_CD3e
89Y_CD45
CD4 T Cells
62.53%
168Er_CD8a
172Yb_CD4
CD8+ Memory T
19.44%
171Yb_CD44
160Gd_CD62L
CD8 T cells
30.74%
CD8+ Naive T Cells
18.91%
DCs
DCs
1.96%
142Nd_CD11c
CD45+ NK1.1+
9.50%
NK Cells
88.89%
153Eu_NKp46
NK cells
CD3-CD19-
17.91%
NK T cells
4.35%
Monocyte Ly6C+ and Ly6C-
162Dy_Ly6C
Monocyte Ly6C+
81.03%
148Nd_CD11b
CD11b+
37.79%
NK T cells
Monocyte Ly6C-
19.35%
Macrophage
Macrophage
25.83%
146Nd_F4-80
156Gd_CD64
B
spleen
BG-vehicle
BG-DTx
BG-DTx + probiotics
Mature B cell
60.53%
Immature B cell
30.39%
Mature B cell
71.51%
Immature B cell
11.93%
Mature B cell
72.77%
Immature B cell
17.61%
IgD
CD24
Bone marrow
Mature B cell
8.25%
Immature B cell
88.67%
Mature B cell
59.92%
Immature B cell
27.94%
Mature B cell
28.64%
Immature B cell
63.56%
C
Relative abundance of CD45+ cells
MLNs
Spleen
BM
T cells
BG-vehicle
BG-DTx
BG-DTx +probiotics
ns
D
CD4 T cells
E
CD8 T cells
F
Ly6Chigh Monocyte
G
Macrophages
H
row min
row max
I
Pearson correlation
0.50
1.00
id
BG-vehicle
BG-DTx
BG-DTx + probiotics

**Figure S6. Related to Figure 6.**

**A.** Depiction of the manual cell subpopulation gating strategy. **B.** In spleen and bone marrow, B cell subpopulations characteristics are changed in BG-DTx mice, an effect partially rescued by probiotics. **C-G.** Relative abundance of immune cells from the total pool of live CD45$^{+}$ cells in mesenteric lymph node, spleen, and bone marrow from BG-vehicle, BG-DTx, and BG-DTx + probiotics mice on a Glp1r-ires-cre × ROSA26-iDTR background. **C:** T cells, **D:** CD4 T cells, **E:** CD8 T cells, **F:** Ly6C$^{high}$ monocytes, **G:** Macrophages. In all cases, 2-way RM-ANOVA, group effect and organ × group (except CD8 T cells) $p<0.006$. Bonferroni BG-vehicle *vs*. BG-DTx, and BG-DTx *vs*. BG-DTx+Probiotics in different organs, all comparisons $*p<0.05$. All other comparisons resulted non-significant (n.s.). **H.** Heatmap of the Olink proteomic results. **I.** Pearson correlation matrices based on the Olink proteomics analyses across BG-vehicle, BG-DTx, and BG-DTx+probiotics groups.

### Figure S7

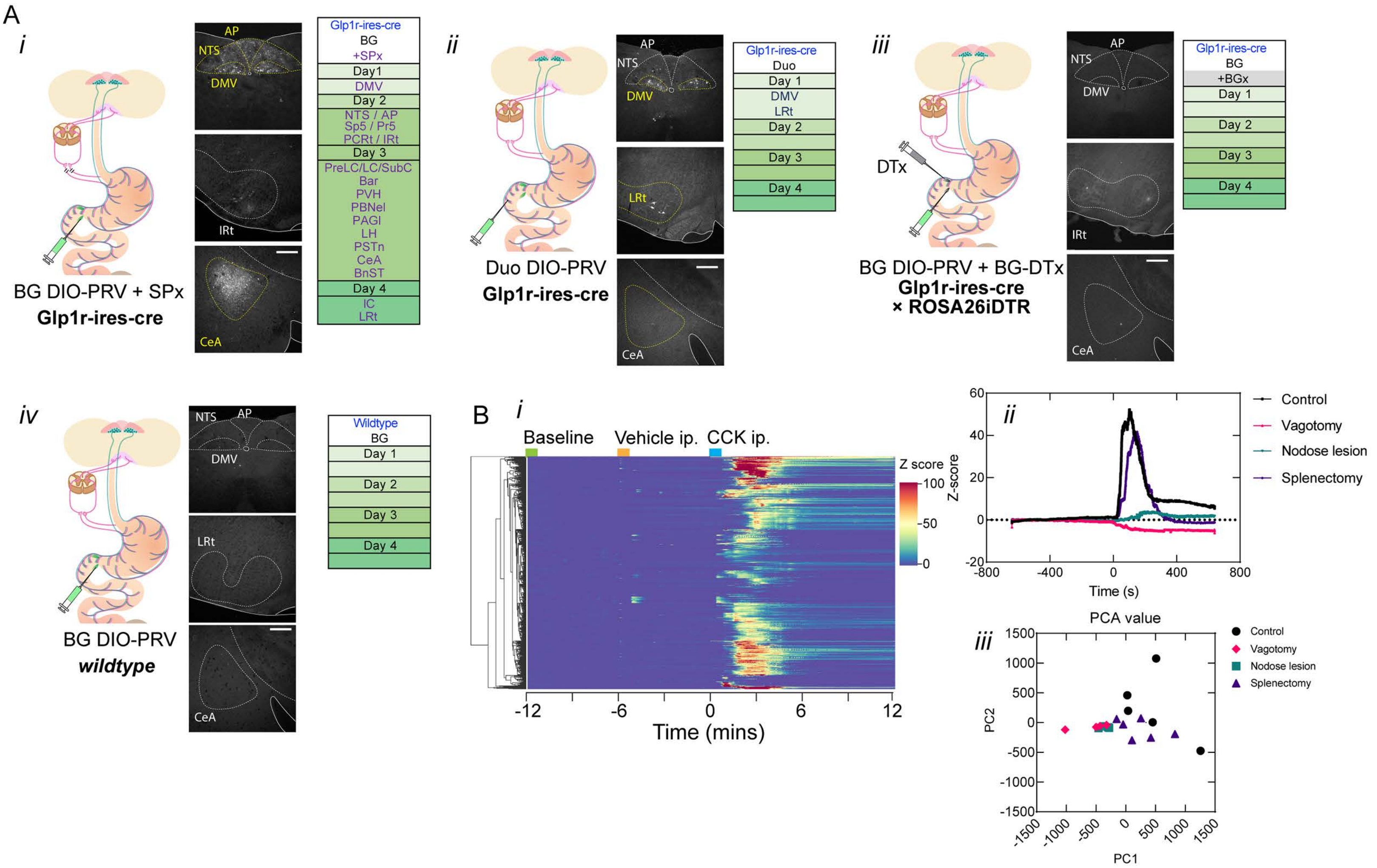
A
i
AP
NTS
DMV
IRt
CeA
Glp1r-ires-cre
BG
+SPx
Day1
DMV
Day 2
NTS / AP
Sp5 / Pr5
PCRt / IRt
Day 3
PreLC/LC/SubC
Bar
PVH
PBNel
PAGl
LH
PSTn
CeA
BnST
Day 4
IC
LRt
BG DIO-PRV + SPx
Glp1r-ires-cre
ii
LRt
Glp1r-ires-cre
Duo
Day 1
DMV
LRt
Day 2
Day 3
Day 4
Duo DIO-PRV
Glp1r-ires-cre
iii
DTx
BG
+BGx
BG DIO-PRV + BG-DTx
Glp1r-ires-cre
× ROSA26iDTR
iv
Wildtype
BG DIO-PRV
wildtype
B
Baseline
Vehicle ip.
CCK ip.
Z score
100
50
0
-12
-6
0
6
12
Time (mins)
Z-score
Control
Vagotomy
Nodose lesion
Splenectomy
Time (s)
PCA value
PC1
PC2

**Figure S7. Related to Figure 7.**

**A.** Control experiments for *Cre*-dependent DIO-PRV tracing. ***(i)*** DIO-PRV tracing from Brunner's glands in Glp1r-ires-cre mice sustaining splenectomies failed to alter labeling; ***(ii)*** DIO-PRV tracing from duodenum Glp1r$^{+}$ enteric neurons in Glp1r-ires-cre mice failed to yield brain labeling; ***(iii)*** DIO-PRV tracing from Brunner's glands in BG lesioned (BG-DTx) Glp1r-ires-cre×ROSA26iDTR mice failed to yield labeling in brain; ***(iv)*** DIO-PRV tracing from Brunner's glands in wildtype mice failed to yield brain labeling. The z-score data for other conditions in ***(ii)*** and ***(iii)*** are also from Figure 1. Bar=100µm. **B.** Duodenal sympathetic denervation (splenectomies) failed to block Brunner's glands calcium transients and secretion after CCK infusions.
