## Supplementary Table 1 for "An Amygdalar-Vagal-Glandular Circuit Controls the Intestinal Microbiome"

**Supplemental Table 1**. Details on surgical procedures.

| Strain | Age | Target | Procedure |  | Age | Procedure | |
| --- | --- | --- | --- | --- | --- | --- | --- |
| Glp1r-ires-Cre × Ai148D | 6 wks | I.P. | Catheterization | | Immediate | I.P. infusion | CCK / Saline |
|  |  | Brunner’s glands | Intravital window | |  | Intravital microscopy | |
| Glp1r-ires-Cre × Ai148D | 5 wks | Nodose Ganglia | Injection | CCK-SAP |  | | |
|  | 6 wks | I.P. | Catheterization | | Immediate | I.P. infusion | CCK / Saline |
|  |  | Brunner’s glands | Intravital window | |  | Intravital microscopy | |
| Glp1r-ires-Cre × Ai148D | 5 wks | Vagotomy | | |  | | |
|  | 6 wks | I.P. | Catheterization | | Immediate | I.P. infusion | CCK / Saline |
|  |  | Brunner’s glands | Intravital window | |  | Intravital microscopy | |
| ChAT-ires-Cre | 6 wks | DMV (AP:-7.5mm, ML: ±0.3mm, DV -5.5 ~-5.3mm) | Viral injection (Unilateral) | AAV1 hSyn FLEx mGFP-2A-Synaptophysin-mRuby | 10 wks | Anatomy analysis | |
| ChAT-ires-Cre | 6 wks | DMV (AP:-7.5mm, ML: ±0.3mm, DV -5.5 ~-5.3mm) | Viral injection (Bilateral) | AAV9-Ef1a-DIO EYFP | 10 wks | Anatomy analysis | |
| C57BL/6J | 4 wks | Brunner's glands | Electric resection / Sham surgery | | 6 wks | I.P. infusion for 6 consective days | CCK ip. |
|  |  |  |  |  | 7 wks | Day7 | Intestine contents collection for 16s rRNA sequencing |
| Glp1r-ires-cre × Ai148D × ROSA26iDTR | 4 wks | Brunner's glands | Injection | DTx / Control-vehicle | 6 wks | Anatomy analysis | |
| Glp1r-ires-cre × Ai148D | 4 wks | Brunner's glands | Electric resection / Sham surgery | | 6 wks | Anatomy analysis | |
| Glp1r-ires-cre × ROSA26iDTR | 4 wks | Brunner's glands | Injection | DTx / Control-vehicle | 6 wks | Anatomy analysis | |
|  |  |  |  |  | 6 wks | Behavior analysis | |
|  |  |  |  |  | 6 wks | In vivo ultrasound analysis | |
| C57BL/6J | 4 wks | Brunner's glands | Electric resection / Sham surgery | | 6 wks | Anatomy analysis | |
|  |  |  |  |  | 6 wks | Behavior analysis | |
|  |  |  |  |  | 6 wks | In vivo ultrasound analysis | |
| Glp1r-ires-cre × ROSA26iDTR | 4 wks | Duodenum | Injection | DTx | 6 wks | Anatomy analysis | |
|  |  |  |  |  | 6 wks | Behavior analysis | |
| C57BL/6J | 4 wks | Duodenum | Electric resection | | 6 wks | Anatomy analysis | |
|  |  |  |  |  | 6 wks | Behavior analysis | |
| Glp1r-ires-cre × ROSA26iDTR | 4 wks | Brunner's glands | Injection | DTx / Control-vehicle | 6 wks | Gavage | 10^10 CFUs of EcAZ-2 |
|  |  |  |  |  | 7 wks | Stools cultures with Kanamycin plate | |
| Glp1r-ires-cre × Ai148D × ROSA26iDTR | 4 wks | Brunner's glands | Injection | DTx / Control-vehicle | 6 wks | Gavage | 10^9 CFUs of Staphylococcus xylosus |
|  |  |  |  |  | 7 wks | Blood cultures with BHI plate | |
| C57BL/6J | 4 wks | Bariatric surgeries: Sham / BG+ bypass / BG- bypass | | | 8 wks | Behavior analysis | |
| Glp1r-ires-cre × ROSA26iDTR | 4 wks | Brunner's glands | Injection | DTx / Control-vehicle | 6 wks | Intra-Cecum infusion for 6 consective days | Probiotics solution / vehicle |
|  |  | Cecum Catheterization | | |  |  |  |
| Glp1r-ires-cre × ROSA26iDTR | 4 wks | Brunner's glands | Injection | DTx / Control-vehicle | 6 wks | Ad libitum drinking for 6 consective days | Mucin solution |
|  |  | Cecum Catheterization | | |  |  |  |
| C57BL/6J | 4 wks | Brunner's glands | Injection | PRV CAG-DIO-TK-GFP | Post-inj Day 1 - 4 | Anatomy analysis | |
| Glp1r-ires-cre | 4 wks | Brunner's glands | Injection | PRV CAG-DIO-TK-GFP | Post-inj Day 1 - 4 | Anatomy analysis | |
| Glp1r-ires-cre | 4 wks | Duodenum | Injection | PRV CAG-DIO-TK-GFP | Post-inj Day 1 - 4 | Anatomy analysis | |
| Glp1r-ires-cre | 3 wks | Vagotomy | | |  | | |
|  | 4 wks | Brunner's glands | Injection | PRV CAG-DIO-TK-GFP | Post-inj Day 1 - 4 | Anatomy analysis | |
| Glp1r-ires-cre | 3 wks | Splanchnicectomy | | |  | | |
|  | 4 wks | Brunner's glands | Injection | PRV CAG-DIO-TK-GFP | Post-inj Day 1 - 4 | Anatomy analysis | |
| Glp1r-ires-cre × ROSA26iDTR | 3 wks | Brunner's glands | Injection | DTx / Control-vehicle |  | | |
|  | 4 wks | Brunner's glands | Injection | PRV CAG-DIO-TK-GFP | Post-inj Day 1 - 4 | Anatomy analysis | |
| Glp1r-ires-cre × Ai148D | 6 wks | CeM (AP:-1.1 mm, ML: ±2.5 mm, DV -5.0mm) | Electrode implantation | | Immediate | I.P. infusion | CCK |
|  |  | I.P. | Catheterization | |  | CeM | electrical stimulation |
|  |  | Brunner’s glands | Intravital window | |  | Intravital microscopy | |
| C57BL/6J | 6 wks | CeM (AP:-1.1 mm, ML: ±2.5 mm, DV -5.0mm) | Viral injection (Bilateral) | AAV5-hSyn-hM3Dq-mCherry | 8 wks | I.P. injection | CCK |
|  |  |  |  |  |  | I.P. injection | CNO |
|  |  |  |  |  |  | Behavior analysis | |
