## Supplementary Table 2 for "An Amygdalar-Vagal-Glandular Circuit Controls the Intestinal Microbiome"

**Supplemental Table 2**. Missing data frequency in Olink proteomics analysis.

| **Olink Target 96 Mouse Exploratory(v.3801)** | | | | | | | |
| --- | --- | --- | --- | --- | --- | --- | --- |
| **Assay** | **Uniprot ID** | **OlinkID** | **Missing Data freq.** | **Assay** | **Uniprot ID** | **OlinkID** | **Missing Data freq.** |
| Clmp | Q8R373 | OID05027 | 0% | Plxna4 | Q80UG2 | OID05077 | 26% |
| Cpe | Q00493 | OID05029 | 0% | Ghrl | Q9EQX0 | OID05108 | 26% |
| Tnfrsf11b | O08712 | OID05036 | 0% | Ahr | P30561 | OID05065 | 29% |
| Pla2g4a | P47713 | OID05038 | 0% | Gcg | P55095 | OID05031 | 30% |
| Tgfa | P48030 | OID05041 | 0% | Plin1 | Q8CGN5 | OID05068 | 32% |
| Epo | P07321 | OID05043 | 0% | Itgb6 | Q9Z0T9 | OID05062 | 36% |
| Fst | P47931 | OID05046 | 0% | Clstn2 | Q9ER65 | OID05053 | 40% |
| Nadk | P58058 | OID05048 | 0% | Ddah1 | Q9CWS0 | OID05087 | 45% |
| Notch3 | Q61982 | OID05050 | 0% | Gdnf | P48540 | OID05032 | 46% |
| Snap29 | Q9ERB0 | OID05051 | 0% | Cant1 | Q8VCF1 | OID05057 | 52% |
| Cntn1 | P12960 | OID05052 | 0% | Fli1 | P26323 | OID05117 | 55% |
| S100a4 | P07091 | OID05054 | 0% | Itgb1bp2 | Q9R000 | OID05095 | 56% |
| Mia | Q61865 | OID05056 | 0% | Il10 | P18893 | OID05088 | 59% |
| Ccl2 | P10148 | OID05066 | 0% | Csf2 | P01587 | OID05092 | 60% |
| Wfikkn2 | Q7TQN3 | OID05069 | 0% | Il1b | P10749 | OID05097 | 60% |
| Qdpr | Q8BVI4 | OID05071 | 0% | Crim1 | Q9JLL0 | OID05080 | 64% |
| Fas | P25446 | OID05074 | 0% | Ccl5 | P30882 | OID05042 | 66% |
| Erbb4 | Q61527 | OID05075 | 0% | Ntf3 | P20181 | OID05114 | 69% |
| Ccl3 | P10855 | OID05079 | 0% | Il6 | P08505 | OID05039 | 78% |
| Hgf | Q08048 | OID05082 | 0% | Pak4 | Q8BTW9 | OID05106 | 93% |
| Sez6l2 | Q4V9Z5 | OID05083 | 0% | Il5 | P04401 | OID05112 | 95% |
| Il1a | P01582 | OID05084 | 0% | Tnf | P06804 | OID05122 | 95% |
| Il23r | Q8K4B4 | OID05085 | 0% | Kitlg | P20826 | OID05058 | 97% |
| Dll1 | Q61483 | OID05086 | 0% |  |  |  |  |
| Tnfrsf12a | Q9CR75 | OID05089 | 0% |  |  |  |  |
| Acvrl1 | Q61288 | OID05090 | 0% |  |  |  |  |
| Lgmn | O89017 | OID05091 | 0% |  |  |  |  |
| Cxcl9 | P18340 | OID05093 | 0% |  |  |  |  |
| Map2k6 | P70236 | OID05094 | 0% |  |  |  |  |
| Casp3 | P70677 | OID05098 | 0% |  |  |  |  |
| Apbb1ip | Q8R5A3 | OID05099 | 0% |  |  |  |  |
| Wisp1 | O54775 | OID05100 | 0% |  |  |  |  |
| Cdh6 | P97326 | OID05101 | 0% |  |  |  |  |
| Pdgfb | P31240 | OID05102 | 0% |  |  |  |  |
| Tgfbr3 | O88393 | OID05104 | 0% |  |  |  |  |
| Cxcl1 | P12850 | OID05105 | 0% |  |  |  |  |
| Cntn4 | Q69Z26 | OID05107 | 0% |  |  |  |  |
| Lpl | P11152 | OID05109 | 0% |  |  |  |  |
| Fstl3 | Q9EQC7 | OID05110 | 0% |  |  |  |  |
| Eda2r | Q8BX35 | OID05113 | 0% |  |  |  |  |
| Tnfsf12 | O54907 | OID05115 | 0% |  |  |  |  |
| Ccl20 | O89093 | OID05116 | 0% |  |  |  |  |
| Tpp1 | O89023 | OID05118 | 0% |  |  |  |  |
| Vegfd | P97946 | OID05120 | 0% |  |  |  |  |
| Matn2 | O08746 | OID05028 | 1% |  |  |  |  |
| Il17a | Q62386 | OID05034 | 1% |  |  |  |  |
| Rgma | Q6PCX7 | OID05047 | 1% |  |  |  |  |
| Ca13 | Q9D6N1 | OID05055 | 1% |  |  |  |  |
| Flrt2 | Q8BLU0 | OID05070 | 2% |  |  |  |  |
| Axin1 | O35625 | OID05044 | 3% |  |  |  |  |
| Riox2 | Q8CD15 | OID05076 | 3% |  |  |  |  |
| Yes1 | Q04736 | OID05033 | 5% |  |  |  |  |
| Tgfb1 | P04202 | OID05037 | 5% |  |  |  |  |
| Cyr61 | P18406 | OID05063 | 5% |  |  |  |  |
| Dlk1 | Q09163 | OID05064 | 5% |  |  |  |  |
| Igsf3 | Q6ZQA6 | OID05103 | 5% |  |  |  |  |
| Tnr | Q8BYI9 | OID05119 | 6% |  |  |  |  |
| Prdx5 | P99029 | OID05040 | 7% |  |  |  |  |
| Ppp1r2 | Q9DCL8 | OID05060 | 7% |  |  |  |  |
| Vsig2 | Q9Z109 | OID05081 | 7% |  |  |  |  |
| Dctn2 | Q99KJ8 | OID05111 | 7% |  |  |  |  |
| Parp1 | P11103 | OID05121 | 7% |  |  |  |  |
| Adam23 | Q9R1V7 | OID05061 | 8% |  |  |  |  |
| Epcam | Q99JW5 | OID05078 | 8% |  |  |  |  |
| Il17f | Q7TNI7 | OID05096 | 11% |  |  |  |  |
| Eno2 | P17183 | OID05067 | 16% |  |  |  |  |
| Foxo1 | Q9R1E0 | OID05035 | 17% |  |  |  |  |
| Gfra1 | P97785 | OID05059 | 17% |  |  |  |  |
| Tnni3 | P48787 | OID05049 | 22% |  |  |  |  |
