## Supplementary Table 3 for "An Amygdalar-Vagal-Glandular Circuit Controls the Intestinal Microbiome"

**Supplemental Table 3.** Details on all statistical tests performed, including type of test, N, degrees of freedom, statistic, and p values.

| **Figure** | **Description** | **Number of**  **animals/trials** | **Statistical test** |  | **Correction for multiple**  **comparison** | **Comparison** | **value** | **Significance** |
| --- | --- | --- | --- | --- | --- | --- | --- | --- |
| **Main Figures** | | | | | | | | |
| 1D | Spontaneous and CCK-induced peaks | Saline^ip^: n of peaks=205  CCK^ip^: n of peaks=1780  After CCK: n of peaks=1092 | One-way ANOVA | F (2, 3074) = 906.4 | Bonferroni post hoc | Group  Peaks z-score:  Spontaneous before vs. CCK-induced  Spontaneous before vs. Spontaneous after  CCK-induced vs. Spontaneous after | <0.0001  <0.0001  0.0876  <0.0001 | *  ns  * |
| 1E |  | Saline^ip^: n of mice=5  CCK^ip^: n of mice=5  After CCK: n of mice=5 | One-way ANOVA | F (2, 8) = 47.28 | Bonferroni post hoc | Group  Percentage of single activated glands:  Spontaneous before vs. CCK-induced  Spontaneous before vs. Spontaneous after  CCK-induced vs. Spontaneous after | <0.0001  <0.0001  0.004  0.0035 | *  *  * |
| 1I (left) | z-score max | Control: n of mice=5  Vagotamy: n of mice=4  Nodose lesion: n of mice=4 | Two-way mixed RM-ANOVA | F (6, 30) = 11.84 | Bonferroni post hoc | lesion×injection  CCK^ip^  Control vs. Vagotomy  Control vs. Nodose lesion  Vagotomy vs. Nodose lesion | <0.001  <0.0001  <0.0001  >0.9999 | *  * |
| 1I (right) | z-score mean | Control: n of mice=5  Vagotamy: n of mice=4  Nodose lesion: n of mice=4 | Two-way mixed RM-ANOVA | F (6, 30) = 10.62 | Bonferroni post hoc | lesion×injection  CCK^ip^  Control vs. Vagotomy  Control vs. Nodose lesion  Vagotomy vs. Nodose lesion | <0.001  <0.0001  <0.0001  0.4121 | *  * |
| 1K (left) | Small intestine | Vehicle^ip^: n of mice=5  CCK^ip^: n of mice=7 | 2-sample t-test | T(10)=2.331 |  | Vehicle^ip^ vs. CCK^ip^ | 0.042 | * |
| 1K (middle) | Large intestine | Vehicle^ip^: n of mice=5  CCK^ip^: n of mice=7 | 2-sample t-test | T(10)=3.616 |  | Vehicle^ip^ vs. CCK^ip^ | 0.0047 | * |
| 1K (right) | feces | Vehicle^ip^: n of mice=11  CCK^ip^: n of mice=13 | 2-sample t-test | T(22)=4.553 |  | Vehicle^ip^ vs. CCK^ip^ | 0.0002 | * |
| 1L (right) | feces | Vehicle^ip^: n of mice=5  CCK^ip^: n of mice=5 | 2-sample t-test | T(8)=2.4 |  | Vehicle^ip^ vs. CCK^ip^ | 0.0432 | * |
| 1M (right) | Relative abundance | Vehicle^ip^: n of mice=7  CCK^ip^: n of mice=10 | 2-sample t-test | T(15)= 2.273 |  | Vehicle^ip^ vs. CCK^ip^ | 0.0382 | * |
| 1N (right) | Relative abundance | Vehicle^ip^: n of mice=7  CCK^ip^: n of mice=11 | 2-sample t-test | T(16) = 2.2836 |  | Vehicle^ip^ vs. CCK^ip^ | 0.0119 | * |
| 1O | Lactobacilli | Sham + vehicle^ip^: n of mice=14  Sham + CCK^ip^: n of mice=16  BG-resected + vehicle^ip^: n of mice=12  BG-resected + CCK^ip^: n of mice=12 | One-way ANOVA | F(3,50)=7.377 | Bonferroni post hoc | Surgery group  Sham + vehicle^ip^ vs. Sham + CCK^ip^  BG-resected + vehicle^ip^ vs. BG-resected + CCK^ip^ | 0.0003  0.0172  >0.9999 | *  ns |
| 1P | Lactobacilli | Sham + vehicle^ip^: n of mice=7  Sham + CCK^ip^: n of mice=8  BG-resected + vehicle^ip^: n of mice=6  BG-resected + CCK^ip^: n of mice=5 | One-way ANOVA | F (3, 22) = 5.219 | Bonferroni post hoc | Surgery group  Sham + vehicle^ip^ vs. Sham + CCK^ip^  BG-resected + vehicle^ip^ vs. BG-resected + CCK^ip^ | 0.0071  0.0372  >0.9999 | *  ns |
| 1Q | staphylococcus | Sham + vehicle^ip^: n of mice=7  Sham + CCK^ip^: n of mice=8  BG-resected + vehicle^ip^: n of mice=6  BG-resected + CCK^ip^: n of mice=5 | One-way ANOVA | F(3,23)=6.121 | Bonferroni post hoc | Surgery group  Sham + vehicle^ip^ vs. Sham + CCK^ip^  BG-resected + vehicle^ip^ vs. BG-resected + CCK^ip^ | 0.0032  >0.9999  0.0187 | ns  * |
| 2A (right) |  | n of mice=4 | Paired t test | T(3)=87.72 |  | Brunner’s glands vs. goblet cells | <0.0001 | * |
| 2B (right) |  | n of mice=6 | Paired t test | T(5)=13.92 |  | Brunner’s glands vs. Villi | <0.0001 | * |
| 2C | Nerve terminals | Brunner’s glands: n of samples=25  Villi: n of samples=15 | 2-sample t-test | T(38)=7.695 |  | Brunner’s glands vs. Villi | <0.0001 | * |
| 3C (left) |  | BG-vehicle: n of mice=5  BG-DTx: n of mice=5 | 2-sample t-test | T(8)=12.70 |  | BG-vehicle vs. BG-DTx | <0.0001 | * |
| 3C (right) |  | Sham: n of mice=5  BG-resected: n of mice=5 | 2-sample t-test | T(8)=7.472 |  | Sham vs. BG-resected | <0.0001 | * |
| 3D (left) |  | BG-vehicle: n of mice=5  BG-DTx: n of mice=5 | 2-sample t-test | T(8)=0.7624 |  | BG-vehicle vs. BG-DTx | 0.4677 | ns |
| 3D (right) |  | Sham: n of mice=5  BG-resected: n of mice=5 | 2-sample t-test | T(8)=0.8931 |  | Sham vs. BG-resected | 0.3979 | ns |
| 3E (left) |  | BG-vehicle: n of mice=5  BG-DTx: n of mice=5 | 2-sample t-test | T(8)=4.143 |  | BG-vehicle vs. BG-DTx | 0.0032 | * |
| 3E (right) |  | BG-vehicle: n of mice=5  BG-DTx: n of mice=5 | 2-sample t-test | T(8)=1.258 |  | BG-vehicle vs. BG-DTx | 0.2439 | ns |
| 3F (left) |  | Sham: n of mice=5  BG-resected: n of mice=5 | 2-sample t-test | T(8)=3.868 |  | Sham vs. BG-resected | 0.0048 | * |
| 3F (right) |  | Sham: n of mice=5  BG-resected: n of mice=5 | 2-sample t-test | T(8)=0.9468 |  | Sham vs. BG-resected | 0.3715 | ns |
| 3G | Food preference | BG-vehicle: n of mice=5  BG-DTx: n of mice=5 | two-way mixed RM-ANOVA | F (1, 8) = 47.69 |  | Food type x Surgery group  High fat: BG-vehicle vs. BG-DTx  High fat (fiber free) : BG-vehicle vs. BG-DTx | 0.0001  0.0003  0.0021 | * |
| 3H | Food preference | Sham: n of mice=5  BG-resected: n of mice=5 | two-way mixed RM-ANOVA | F (1, 8) = 25.60 |  | Food type x Surgery group  High fat: Sham vs. BG-resected  High fat (fiber free) : Sham vs. BG-resected | 0.001  0.0005  0.0015 | * |
| 3I | Stomach size | BG-vehicle: n of mice=6  BG-DTx: n of mice=9 | 2-sample t-test | T(13)=2.816 |  | BG-vehicle vs. BG-DTx | 0.0146 | * |
| 3J | Stomach size | Sham: n of mice=6  BG-resected: n of mice=15 | 2-sample t-test | T(19)=2.979 |  | Sham vs. BG-resected | 0.0077 | * |
| 3K | spleen | BG-vehicle: n of mice=6  BG-DTx: n of mice=5 | 2-sample t-test | T(9)=4.360 |  | BG-vehicle vs. BG-DTx | 0.0018 | * |
| 3L | Germinal center | BG-vehicle: n=14  BG-DTx: n=15 | 2-sample t-test | T(27)=5.045 |  | BG-vehicle vs. BG-DTx | <0.0001 | * |
| 3M | B cells | BG-vehicle: n of mice=6  BG-DTx: n of mice=6 | 2-sample t-test | T(10)=14.54 |  | BG-vehicle vs. BG-DTx | <0.0001 | * |
| 3N | Ki67 | BG-vehicle: n of mice=6  BG-DTx: n of mice=6 | 2-sample t-test | T(10)=11.07 |  | BG-vehicle vs. BG-DTx | <0.0001 | * |
| 3O | FDC light zone | BG-vehicle: n of mice=6  BG-DTx: n of mice=6 | 2-sample t-test | T(10)=7.856 |  | BG-vehicle vs. BG-DTx | <0.0001 | * |
| 3Q | EcAZ-2 | BG-vehicle: n of mice=5  BG-DTx: n of mice=5 | 2-sample t-test | T(8)=5.875 |  | BG-vehicle vs. BG-DTx | <0.0001 | * |
| 3R | Survival curve | BG-vehicle (*S. xylosus* ig): n of mice=5  BG-DTx (*S. xylosus* ig): n of mice=7  BG-vehicle (control ig): n of mice=6  BG-DTx (control ig): n of mice=6 | Log Rank (Mantel-Cox) | χ2(3)=27.88 | Bonferroni post hoc | Group  BG-DTx (*S. xylosus* ig) vs. BG-vehicle (*S. xylosus* ig)  BG-DTx (*S. xylosus* ig) vs. BG-DTx (control ig)  BG-DTx (*S. xylosus* ig) vs. BG-vehicle (control ig) | 0.0001  0.0022  0.0001  0.0001 | *  ** |
| 3S (left) |  | BG-vehicle: n of mice=5  BG-DTx: n of mice=6 | 2-sample t-test | T(9)=3.922 |  | BG-vehicle vs. BG-DTx | 0.0035 | * |
| 3S (right) |  | BG-vehicle: n of mice=5  Duo-DTx: n of mice=6 | 2-sample t-test | T(9)=2.105 |  | BG-vehicle vs. Duo-DTx | 0.0646 | ns |
| 3T |  | BG-vehicle: n of mice=6  BG-DTx: n of mice=6 | 2-sample t-test | T(10)=2.704 |  | BG-vehicle vs. BG-DTx | 0.0221 | * |
| 3U | Abx | BG-vehicle: n of mice=6  BG-DTx: n of mice=6 | Log Rank (Mantel-Cox) | χ2(3)=24.107 |  | Group | <0.001 | * |
| 3V | Probiotics preference | BG-vehicle: n of mice=5  BG-DTx: n of mice=6 | two-way mixed RM-ANOVA | F (1, 9) = 2.061  F (1, 9) = 6.117  F (1, 9) = 19.70 | Bonferroni post hoc | training x Surgery group  training  Surgery group  Baseline vs Test:  BG-vehicle  BG-DTx | 0.1849  0.0354  0.0016  >0.9999  0.0353 | * |
| 4F | Food preference | Sham: n of mice=5  BG+ bypass: n of mice=5  BG- bypass: n of mice=5 | two-way mixed RM-ANOVA | F(2,12) = 29.952 | Bonferroni post hoc | Surgery group x food type  Sham vs. BG- bypass  Sham vs, BG+ bypass  BG+ bypass vs. BG- bypass | <0.001  0.011  >0.999  0.008 | * |
| 4G | Probiotics preference | Sham: n of mice=4  BG+ bypass: n of mice=5  BG- bypass: n of mice=5 |  | F (2, 11) = 2.203  F (1, 11) = 39.14  F (2, 11) = 0.7747 | Bonferroni post hoc | training x Surgery group  training  Surgery group  Baseline vs Test:  Sham:  BG+ bypass:  BG- bypass | 0.1568  <0.0001  0.4844  0.3329  0.0032  0.0013 | ns  *  * |
| 4H | MRS plate | Sham: n of mice=4  BG+ bypass: n of mice=5  BG- bypass: n of mice=5 | One-way ANOVA | F(2,12)=8.172 | Bonferroni post hoc | Group  Sham vs. BG+ bypass  Sham vs. BG- bypass  BG+ bypass vs. BG- bypass | 0.0058  0.0146  >0.9999  0.0118 | *  * |
| 4I |  | Sham: n=21  BG+ bypass: n=14  BG- bypass: n=21 | One-way ANOVA | F (2, 53) = 9.857 | Bonferroni post hoc | Group  Sham vs. BG+ bypass  Sham vs. BG- bypass  BG+ bypass vs. BG- bypass | 0.0002  0.0013  >0.9999  0.0004 | *  * |
| 4J |  | n=56 | Pearson | R^2^=0.2853 |  |  | <0.001 | * |
| 4L | EcAZ-2 | Sham: n of mice=5  BG+ bypass: n of mice=6  BG- bypass: n of mice=6 | One-way ANOVA | F(2,14)=9.763 | Bonferroni post hoc | Group  Sham vs. BG+ bypass  Sham vs. BG- bypass  BG+ bypass vs. BG- bypass | 0.0022  >0.9999  0.0069  0.0051 | ns  *  ** |
| 5C | spleen | BG-DTx+control^ic^: n of mice=5  BG-DTx+probiotics^ic^: n of mice=5  BG-vehicle+control^ic^: n of mice=5  BG-vehicle+probiotic^ic^: n of mice=5 | One-way ANOVA | F(3,16) = 22.15 | Bonferroni post hoc | Group  BG-DTx+control^ic^ vs. BG-DTx+probiotics^ic^  BG-DTx+probiotics^ic^ vs. BG-vehicle+control^ic^ | <0.001  0.006  >0.9 | *  ns |
| 5E | GC | BG-DTx+control^ic^: n=20  BG-DTx+probiotics^ic^: n=11  BG-vehicle+control^ic^: n =11  BG-vehicle+probiotic^ic^: n=20 | One-way ANOVA | F (3, 58) = 14.43 | Bonferroni post hoc | Group  BG-DTx+control^ic^ vs. BG-DTx+probiotics^ic^  BG-DTx+probiotics^ic^ vs. BG-vehicle+control^ic^ | <0.0001  0.0002  >0.9 | *  ns |
| 5G | B cell | BG-DTx+control^ic^: n=10  BG-DTx+probiotics^ic^: n=8  BG-vehicle+control^ic^: n =8  BG-vehicle+probiotic^ic^: n=10 | One-way ANOVA | F (3, 32) = 89.20 | Bonferroni post hoc | Group  BG-DTx+control^ic^ vs. BG-DTx+probiotics^ic^  BG-DTx+probiotics^ic^ vs. BG-vehicle+control^ic^ | <0.001  0.0001  >0.5 | *  ns |
| 5H | Ki67 | BG-DTx+control^ic^: n=10  BG-DTx+probiotics^ic^: n=8  BG-vehicle+control^ic^: n =9  BG-vehicle+probiotic^ic^: n=10 | One-way ANOVA | F(3,33)=14.453 | Bonferroni post hoc | Group  BG-DTx+control^ic^ vs. BG-DTx+probiotics^ic^  BG-DTx+probiotics^ic^ vs. BG-vehicle+control^ic^ | <0.001  <0.0001  >0.5 | *  ns |
| 5I | FDC | BG-DTx+control^ic^: n=11  BG-DTx+probiotics^ic^: n=8  BG-vehicle+control^ic^: n =9  BG-vehicle+probiotic^ic^: n=10 | One-way ANOVA | F (3, 34) = 31.35 | Bonferroni post hoc | Group  BG-DTx+control^ic^ vs. BG-DTx+probiotics^ic^ | <0.001  <0.001 | * |
| 5K |  | BG-DTx+control^ic^ (*S. xylosus* ig): n of mice=5  BG-DTx+probiotics^ic^ (*S. xylosus* ig): n of mice=6  BG-DTx+control^ic^ (vehicle ig): n of mice=6  BG-DTx+probiotics^ic^ (vehicle ig): n of mice=6 | Log Rank (Mantel-Cox) | χ2(3)=24.107 |  | Group  BG-DTx+control^ic^ (*S. xylosus* ig) vs. BG-DTx+probiotics^ic^ (*S. xylosus* ig) | <0.001  0.0001 | *  ** |
| 5L |  | BG-vehicle+control^ic^ (*S. xylosus* ig): n of mice=6  BG-vehicle+probiotics^ic^ (*S. xylosus* ig): n of mice=6  BG-vehicle+control^ic^ (vehicle ig): n of mice=6  BG-vehicle+probiotics^ic^ (vehicle ig): n of mice=6 | Log Rank (Mantel-Cox) | χ2(3)=4.454 |  | Group | 0.216 |  |
| 5M |  | BG-DTx+control^ic^: n of mice=5  BG-DTx+probiotics^ic^: n of mice=7  BG-vehicle+control^ic^: n of mice=8  BG-vehicle+probiotic^ic^: n of mice=6 | One-way ANOVA | F(3,22) = 30.014 | Bonferroni post hoc | Group  BG-DTx+control^ic^ vs. BG-DTx+probiotics^ic^  BG-DTx+control^ic^ vs. BG-vehicle+control^ic^  BG-DTx+control^ic^ vs. BG-vehicle+probiotic^ic^ | <0.001  <0.001  <0.001  <0.001 | * |
| 5O |  | BG-DTx+control (*S. xylosus* ig): n of mice=7  BG-DTx+mucin (*S. xylosus* ig): n of mice=7  BG-DTx+control (vehicle ig): n of mice=6  BG-DTx+mucin (vehicle ig): n of mice=6 | Log Rank (Mantel-Cox) | χ2(3)=21.61 |  | Group  BG-DTx+control (*S. xylosus* ig) vs. BG-DTx+mucin (*S. xylosus* ig) | <0.001  0.0189 | *  ** |
| 5P |  | BG-vehicle+control (*S. xylosus* ig): n of mice=7  BG-vehicle+mucin (*S. xylosus* ig): n of mice=7  BG-vehicle+control (vehicle ig): n of mice=6  BG-vehicle+mucin (vehicle ig): n of mice=6 | Log Rank (Mantel-Cox) | χ2(3)=4.65 |  | Group | 0.199 |  |
| 5Q |  | BG-DTx+control: n of mice=5  BG-DTx+mucin: n of mice=5  BG-vehicle+control: n of mice=5  BG-vehicle+mucin: n of mice=5 | One-way ANOVA | F(3,16) = 11.34 | Bonferroni post hoc | Group  BG-DTx+control^ic^ vs. BG-DTx+probiotics^ic^  BG-DTx+control^ic^ vs. BG-vehicle+control^ic^  BG-DTx+control^ic^ vs. BG-vehicle+probiotic^ic^ | <0.001  0.001  0.001  0.001 | * |
| 6A | B cells | BG-vehicle: n of mice=5  BG-DTx: n of mice=4  BG-DTx+probiotics: n of mice=6 | two-way mixed RM-ANOVA | F(4,24)=7.718  F(2,12)=210.08 | Bonferroni post hoc | organ x group  group  MLNs  BG-vehicle vs. BG-DTx:  BG-vehicle vs. BG-DTx+probiotics:  BG-DTx vs. BG-DTx+probiotics:  Spleen  BG-vehicle vs. BG-DTx:  BG-vehicle vs. BG-DTx+probiotics:  BG-DTx vs. BG-DTx+probiotics:  BM  BG-vehicle vs. BG-DTx:  BG-vehicle vs. BG-DTx+probiotics:  BG-DTx vs. BG-DTx+probiotics: | < 0.001  < 0.001  <0.0001  >0.9999  <0.0001  <0.0001  0.0346  <0.0001  <0.0001  <0.0001  0.2056 | *  *  *  *  *  ns |
| 6C | Immature B cells | BG-vehicle: n of mice=5  BG-DTx: n of mice=4  BG-DTx+probiotics: n of mice=6 | two-way mixed RM-ANOVA | F(4,24)=97.221  F(2,12)=247.972 | Bonferroni post hoc | organ x group  group  MLNs  BG-vehicle vs. BG-DTx:  BG-vehicle vs. BG-DTx+probiotics:  BG-DTx vs. BG-DTx+probiotics:  Spleen  BG-vehicle vs. BG-DTx:  BG-vehicle vs. BG-DTx+probiotics:  BG-DTx vs. BG-DTx+probiotics:  BM  BG-vehicle vs. BG-DTx:  BG-vehicle vs. BG-DTx+probiotics:  BG-DTx vs. BG-DTx+probiotics: | < 0.001  < 0.001  0.0107  0.9006  0.0913  <0.0001  <0.0001  <0.0001  <0.0001  <0.0001  <0.0001 | *  ns  *  *  *  * |
| 6D | Mature B cells | BG-vehicle: n of mice=5  BG-DTx: n of mice=4  BG-DTx+probiotics: n of mice=6 | two-way mixed RM-ANOVA | F(4,24)=9.307  F(2,12)=8.78 | Bonferroni post hoc | organ x group  group  MLNs  BG-vehicle vs. BG-DTx:  BG-vehicle vs. BG-DTx+probiotics:  BG-DTx vs. BG-DTx+probiotics:  Spleen  BG-vehicle vs. BG-DTx:  BG-vehicle vs. BG-DTx+probiotics:  BG-DTx vs. BG-DTx+probiotics:  BM  BG-vehicle vs. BG-DTx:  BG-vehicle vs. BG-DTx+probiotics:  BG-DTx vs. BG-DTx+probiotics: | <0.001  0.004  <0.0001  >0.9999  <0.0001  0.0098  0.0064  >0.9999  0.1672  >0.9999  0.5971 | *  *  *  ns  ns  ns |
| 6E | Dendrite cells | BG-vehicle: n of mice=5  BG-DTx: n of mice=4  BG-DTx+probiotics: n of mice=6 | two-way mixed RM-ANOVA | F(4,24)=19.371  F(2,12)=28.586 | Bonferroni post hoc | organ x group  group  MLNs  BG-vehicle vs. BG-DTx:  BG-vehicle vs. BG-DTx+probiotics:  BG-DTx vs. BG-DTx+probiotics:  Spleen  BG-vehicle vs. BG-DTx:  BG-vehicle vs. BG-DTx+probiotics:  BG-DTx vs. BG-DTx+probiotics:  BM  BG-vehicle vs. BG-DTx:  BG-vehicle vs. BG-DTx+probiotics:  BG-DTx vs. BG-DTx+probiotics: | <0.001  <0.001  0.1686  >0.9999  0.0269  <0.0001  <0.0001  <0.0001  >0.9999  >0.9999  >0.9999 | ns  *  *  *  ns  ns |
| 6F | NK cells | BG-vehicle: n of mice=5  BG-DTx: n of mice=4  BG-DTx+probiotics: n of mice=6 | two-way mixed RM-ANOVA | F(4,24)=44.157  F(2,12)=35.587 | Bonferroni post hoc | organ x group  group  MLNs  BG-vehicle vs. BG-DTx:  BG-vehicle vs. BG-DTx+probiotics:  BG-DTx vs. BG-DTx+probiotics:  Spleen  BG-vehicle vs. BG-DTx:  BG-vehicle vs. BG-DTx+probiotics:  BG-DTx vs. BG-DTx+probiotics:  BM  BG-vehicle vs. BG-DTx:  BG-vehicle vs. BG-DTx+probiotics:  BG-DTx vs. BG-DTx+probiotics: | <0.001  0.001  0.129  >0.9999  0.0845  <0.0001  <0.0001  <0.0001  0.4525  0.8958  >0.9999 | ns  ns  *  *  ns  ns |
| 6G | Ly6C^low^ monocytes | BG-vehicle: n of mice=5  BG-DTx: n of mice=4  BG-DTx+probiotics: n of mice=6 | two-way mixed RM-ANOVA | F(4,24)=5.521  F(2,12)=54.238 | Bonferroni post hoc | organ x group  group  MLNs  BG-vehicle vs. BG-DTx:  BG-vehicle vs. BG-DTx+probiotics:  BG-DTx vs. BG-DTx+probiotics:  Spleen  BG-vehicle vs. BG-DTx:  BG-vehicle vs. BG-DTx+probiotics:  BG-DTx vs. BG-DTx+probiotics:  BM  BG-vehicle vs. BG-DTx:  BG-vehicle vs. BG-DTx+probiotics:  BG-DTx vs. BG-DTx+probiotics: | 0.003  <0.001  >0.9999  0.2659  0.0814  <0.0001  0.0071  <0.0001  0.7592  0.0002  <0.0001 | ns  ns  *  *  ns  * |
| 6M |  | BG-vehicle: n of mice=5  BG-DTx: n of mice=9  BG-DTx+probiotics: n of mice=6 | One-way ANOVA | F(2,17)=57.595 | Bonferroni post hoc | group  BG-vehicle vs. BG-DTx:  BG-vehicle vs. BG-DTx+probiotics:  BG-DTx vs. BG-DTx+probiotics: | <0.001  <0.001  0.817  <0.001 | *  ns  * |
| 6N |  | BG-vehicle: n of mice=5  BG-DTx: n of mice=9  BG-DTx+probiotics: n of mice=6 | One-way ANOVA | F(2,17)=16.081 | Bonferroni post hoc | group  BG-vehicle vs. BG-DTx:  BG-vehicle vs. BG-DTx+probiotics:  BG-DTx vs. BG-DTx+probiotics: | <0.001  <0.001  >0.999  <0.001 | *  ns  * |
| 6O |  | BG-vehicle: n of mice=5  BG-DTx: n of mice=9  BG-DTx+probiotics: n of mice=6 | One-way ANOVA | F(2,17)=15.325 | Bonferroni post hoc | group  BG-vehicle vs. BG-DTx:  BG-vehicle vs. BG-DTx+probiotics:  BG-DTx vs. BG-DTx+probiotics: | <0.001  0.039  0.121  <0.001 | *  ns  * |
| 6P |  | BG-vehicle: n of mice=5  BG-DTx: n of mice=9  BG-DTx+probiotics: n of mice=6 | One-way ANOVA | F(2,17)=12.425 | Bonferroni post hoc | group  BG-vehicle vs. BG-DTx:  BG-vehicle vs. BG-DTx+probiotics:  BG-DTx vs. BG-DTx+probiotics: | <0.001  0.022  0.566  <0.001 | *  ns  * |
| 6Q |  | BG-vehicle: n of mice=5  BG-DTx: n of mice=9  BG-DTx+probiotics: n of mice=6 | One-way ANOVA | F(2,17)=4.934 | Bonferroni post hoc | group  BG-vehicle vs. BG-DTx:  BG-vehicle vs. BG-DTx+probiotics:  BG-DTx vs. BG-DTx+probiotics: | 0.02  0.2456  >0.9999  0.0217 | ns  ns  * |
| 7D | Max z score | n of mice=3  n of glands=300 | One-way ANOVA | F (4, 1196) = 406.2 | Bonferroni post hoc | group  Baseline vs. Electrical Stim. 1  Baseline vs. CCK^low^  Baseline vs. Electrical Stim. 2  Baseline vs. Electrical Stim. 3 | <0.0001  >0.9999  <0.0001  <0.0001  <0.0001 | * |
| 7E | Lactobacilli | Control (CNO): n of mice=5  Control (CCK^low^): n of mice=5  CeA-Gq (vehicle): n of mice=5  CeA-Gq (CNO): n of mice=5  CeA-Gq (CNO+CCK^low^): n of mice=5 | One-way ANOVA | F (4, 20) = 53.01 | Bonferroni post hoc | group  Control (CNO) vs. Control (CCK^low^)  Control (CNO) vs. CeA-Gq (vehicle)  Control (CNO) vs. CeA-Gq (CNO)  Control (CNO) vs. CeA-Gq (CNO+ CCK^low^)  Control (CCK^low^) vs. CeA-Gq (vehicle)  Control (CCK^low^) vs. CeA-Gq (CNO)  Control (CCK^low^) vs. CeA-Gq (CNO+ CCK^low^)  CeA-Gq (vehicle) vs. CeA-Gq (CNO)  CeA-Gq (vehicle) vs. CeA-Gq (CNO+ CCK^low^)  CeA-Gq (CNO) vs. CeA-Gq (CNO+ CCK^low^) | <0.0001  >0.9999  >0.9999  0.001  <0.0001  >0.9999  0.0003  <0.0001  0.0006  <0.0001  <0.0001 | *  *  *  *  * |
| 7F | EcAZ-2 | CeA-Gq (vehicle): n of mice=6  CeA-Gq (CNO+ CCK^low^): n of mice=6 | 2-sample t-test | T(10)=2.234 |  | CeA-Gq (vehicle) vs. CeA-Gq (CNO+ CCK^low^) | 0.0495 | * |
| 7G | Germinal center | CeA-Gq (vehicle): n=33 (4 mice)  CeA-Gq (CNO+ CCK^low^): n=20 (4 mice) | 2-sample t-test | T(51)=2.517 |  | CeA-Gq (vehicle) vs. CeA-Gq (CNO+ CCK^low^) | 0.015 | * |
| 7H | correlation | n=56 (paired data) | Pearson | R^2^=0.1501 |  | Lactobacilli CFU counts vs. spleen germinal center sizes | 0.032 | * |
| 7I | DMV c-fos | CeA-Gq (vehicle): n of mice=6  CeA-Gq (CNO+ CCK^low^): n of mice=6 | 2-sample t-test | T(10)=4.869 |  | CeA-Gq (vehicle) vs. CeA-Gq (CNO+ CCK^low^) | 0.0007 | * |
| **Supplementary Figures** | | | | | | | | |
| S1B (right) | Brunner’s glands | Vehicle^ip^: n of mice=7  CCK^ip^: n of mice=7 | 2-sample t-test | T(12)=6.818 |  | Vehicle^ip^ vs. CCK^ip^ | <0.001 | * |
| S1C | void score | Vehicle^ip^: n of glands=61  CCK^ip^: n of glands=49 | 2-sample t-test | T(108)=4.634 |  | Vehicle^ip^ vs. CCK^ip^ | <0.0001 | * |
| S1D (*ii*) | Mucin thickness, duo | Vehicle^ip^: n of mice=6  CCK^ip^: n of mice=6 | 2-sample t-test | T(10)=7.410 |  | Vehicle^ip^ vs. CCK^ip^ | <0.001 | * |
| S1D (*iii*) | Mucin thickness, ileum | Vehicle^ip^: n of mice=6  CCK^ip^: n of mice=6 | 2-sample t-test | T(10)=0.3742 |  | Vehicle^ip^ vs. CCK^ip^ | 0.7161 | ns |
| S1D (*iv*) | Mucin thickness, colon | Vehicle^ip^: n of mice=6  CCK^ip^: n of mice=6 | 2-sample t-test | T(10)=1.183 |  | Vehicle^ip^ vs. CCK^ip^ | 0.2642 | ns |
| S1E | Goblet cells | Vehicle^ip^: n of mice=6  CCK^ip^: n of mice=6 | two-way mixed RM-ANOVA | F (2, 20) = 2.111  F (1, 10) = 0.08691  F (2, 20) = 5.811 | Bonferroni post hoc | Injection×segment  Injection  Segment  Vehicle^ip^ vs. CCK^ip^ in duodenum  Vehicle^ip^ vs. CCK^ip^ in ileum  Vehicle^ip^ vs. CCK^ip^ in colon | 0.1473  0.7742  0.0102  0.6175  >0.9999  0.6001 | ns |
| S1F | CD63 | saline^ip^: n of mice=3  CCK^ip^: n of mice=3 | Two-way mixed RM-ANOVA | F (1, 10731) = 1604 |  | saline^ip^ vs. CCK^ip^ | 0.0001 | * |
| S1J (right) |  | n of mice=3 | Paired t test | T(2)=27.69 |  | Brunner’s glands vs. Goblet cells | 0.0013 | * |
| S1L (right) | Nodose lesion | Blank-SAP: n=10  CCK-SAP: n=7 | 2-sample t-test | T(15)=7.852 |  | Blank-SAP vs CCK-SAP | <0.0001 | * |
| S1BB | staphylococcus | Sham + vehicle^ip^: n of mice=14  Sham + CCK^ip^: n of mice=16  BG-resected + vehicle^ip^: n of mice=12  BG-resected + CCK^ip^: n of mice=12 | One-way ANOVA | F (3, 50) = 6.421 | Bonferroni post hoc | Surgery group  Sham + vehicle^ip^ vs. Sham + CCK^ip^  BG-resected + vehicle^ip^ vs. BG-resected + CCK^ip^ | 0.0009  0.4932  0.1274 | *  ns  ns |
| S3B |  | BG-vehicle: n of mice=5  BG-DTx: n of mice=5 | 2-sample t-test | T(8)=0.1210 |  | BG-vehicle vs. BG-DTx | 0.9066 | ns |
| S3C | IPGTT | BG-vehicle: n of mice=5  BG-DTx: n of mice=5 | 2-way mixed RM-ANOVA | F(5, 40)= 0.4558  F(1, 8)= 0.6494 |  | Sampling time x Surgery group  Surgery group | 0.8065  0.4436 | ns  ns |
| S3D | OGTT | BG-vehicle: n of mice=5  BG-DTx: n of mice=5 | 2-way mixed RM-ANOVA | F(5, 40)= 2.036  F(1, 8)=0.2004 |  | Sampling time x Surgery group  Surgery group | 0.0942  0.6663 | ns  ns |
| S3F | Food preference | BG-vehicle: n of mice=5  BG-DTx: n of mice=5 | 2-way mixed RM-ANOVA | F (1, 8) = 1.112  F(1, 8)=10.36 | Bonferroni post hoc | Food preference x Surgery group  Surgery group  BGx-DTx - BG-vehicle  High fat  Normal chow | 0.3225  0.0123  0.0223  0.4880 | ns |
| S3G | Food preference | BG-vehicle: n of mice=5  BG-DTx: n of mice=5 | 2-way mixed RM-ANOVA | F (1, 8) = 0.9908  F (1, 8) = 0.7550 | Bonferroni post hoc | Food preference x Surgery group  Surgery group  BGx-DTx - BG-vehicle  High fat (fiber free)  High protein (fiber free) | 0.3487  0.4102  0.4111  >0.9999 | ns |
| S3H | Food preference | BG-vehicle: n of mice=5  BG-DTx: n of mice=5 | 2-way mixed RM-ANOVA | F (1, 8) = 0.1897  F (1, 8) = 0.1897 | Bonferroni post hoc | Food preference x Surgery group  Surgery group  BGx-DTx - BG-vehicle  High protein (fiber free)  High fiber | 0.6747  0.6747  >0.9999  >0.9999 | ns |
| S3I | Food preference | BG-resected: n of mice=6  Sham: n of mice=5 | 2-way mixed RM-ANOVA | F (1, 9) = 0.5022  F (1, 9) = 0.3694 | Bonferroni post hoc | Food preference x Surgery group  Surgery group  BGx-resected - Sham  High fat  Normal chow | 0.4965  0.5584  0.7269  >0.9999 | ns |
| S3J | Food preference | BG-resected: n of mice=5  Sham: n of mice=4 | 2-way mixed RM-ANOVA | F (1, 7) = 0.09771  F (1, 7) = 0.0002131 | Bonferroni post hoc | Food preference x Surgery group  Surgery group  BGx-resected – Sham  High fat (fiber free)  High protein (fiber free) | 0.7637  0.9888  >0.9999  >0.9999 | ns |
| S3K | Food preference | BG-resected: n of mice=5  Sham: n of mice=4 | 2-way mixed RM-ANOVA | F (1, 7) = 2.894  F (1, 7) = 2.972 | Bonferroni post hoc | Food preference x Surgery group  Surgery group  BGx-resected – Sham  High protein (fiber free)  High fiber | 0.1327  0.1284  0.0592  >0.9999 | ns |
| S3M | Stomach size | Duo-lesion: n of mice=4  Sham: n of mice=6 | 2-sample t-test | T(8)=0.2382 |  |  | 0.8177 | ns |
| S3Q |  | BG-vehicle: n of mice=5  BG-DTx: n of mice=6  Duo-DTx: n of mice=6  Sham: n of mice=5  BG-resected: n of mice=6  Duo-lesion: n of mice=5 | One-way ANOVA | F (5, 27) = 11.08 | Bonferroni post hoc | Surgery group  BG-vehicle vs. BG-DTx  BG-vehicle vs. Duo-DTx  Sham vs. BG-resection  Sham vs, Duo-lesion | <0.0001  0.0005  >0.9999  0.0237  0.0022 | *  ns  **  *** |
| S3R | Abx mice body-weight monitor | BG-vehicle-feces transplant mice: n=6  BG-DTx-feces transplant mice: n=6 | 2-way mixed RM-ANOVA | F(2,20)=5.665  F(1,10)=0.048 |  | group×time  group | 0.011  0.831 | * |
| S3S |  | BG-vehicle: n of mice=5  BG-DTx: n of mice=5 | 2-way mixed RM-ANOVA | F (2, 16) = 1.813  F (1, 8) = 0.1183 |  | region x Surgery group  Surgery group | 0.1951  0.7397 | ns |
| S3T middle |  | BG-vehicle: n=9  BG-DTx: n=9 | 2-sample t-test | T(16)=0.1940 |  | BG-vehicle vs. BG-DTx | 0.8486 | ns |
| S3T right |  | BG-vehicle: n of mice=5  BG-DTx: n of mice=5 | 2-sample t-test | T(8)= 0.03135 |  | BG-vehicle vs. BG-DTx | 0.9758 | ns |
| S4A | IPGTT | Sham: n of mice=5  BG+ bypass: n of mice=6  BG- bypass: n of mice=6 | 2-way mixed RM-ANOVA | F(10,70) = 2.747  F(2,14)=0.398 |  | surgery group x sampling time  surgery group | 0.007  0.679 |  |
| S4B | OGTT | Sham: n of mice=5  BG+ bypass: n of mice=5  BG- bypass: n of mice=5 | 2-way mixed RM-ANOVA | F(10,60) = 1.408  F(2,14)=2.223 |  | surgery group x sampling time  surgery group | 0.199  0.902 |  |
| S4D | Spleen weight | Sham: n of mice=4  BG+ bypass: n of mice=5  BG- bypass: n of mice=5 | One-way ANOVA | F(2,11) =1.813 |  | Surgery group | 0.2087 |  |
| S4E | Body weight | Sham: n of mice=4  BG+ bypass: n of mice=5  BG- bypass: n of mice=5 | One-way ANOVA | F(2,11)=0.04796 |  | Surgery group | 0.9534 |  |
| S5A left | Body weight monitor before surgery | BG-DTx+control^ic^: n of mice=6  BG-DTx+probiotics^ic^: n of mice=5  BG-vehicle+control^ic^: n of mice=5  BG-vehicle+probiotic^ic^: n of mice=5 | 2-way mixed RM-ANOVA | F (9, 51) = 0.6443  F (3, 17) = 1.057 |  | Day x Surgery group  Surgery group | 0.7539  0.3932 |  |
| S5A right | Body weight monitor after surgery | BG-DTx+control^ic^: n of mice=6  BG-DTx+probiotics^ic^: n of mice=5  BG-vehicle+control^ic^: n of mice=5  BG-vehicle+probiotic^ic^: n of mice=5 | 2-way mixed RM-ANOVA | F (9, 51) = 4.736  F (3, 17) = 4.701 |  | Day x Surgery group  Surgery group | 0.0001  0.0144 | * |
| S5B (i) |  | n of mice=7 | Paired t test | T(6)=5.174 |  | Before vs After | 0.0021 | * |
| S5B (ii) |  | n of mice=8 | Paired t test | T(7)=1.841 |  | Before vs After | 0.1081 | ns |
| S5B (iii) |  | n of mice=5 | Paired t test | T(4)=1.156 |  | Before vs After | 0.3122 | ns |
| S5B(iv) |  | n of mice=5 | Paired t test | T(4)=0.7608 |  | Before vs After | 0.4892 | ns |
| S5C (i) |  | n of mice=6 | Paired t test | T(5)=3.412 |  | Before vs After | 0.0190 | * |
| S5C (ii) |  | n of mice=5 | Paired t test | T(4)=1.114 |  | Before vs After | 0.3276 | ns |
| S5C (iii) |  | n of mice=5 | Paired t test | T(4)=1.849 |  | Before vs After | 0.1382 | ns |
| S5C(iv) |  | n of mice=5 | Paired t test | T(4)=1.965 |  | Before vs After | 0.1208 | ns |
| S6C | T cells | BG-vehicle: n of mice=5  BG-DTx: n of mice=4  BG-DTx+probiotics: n of mice=6 | two-way mixed RM-ANOVA | F(4,24)=4.959  F(2,12)=17.709 | Bonferroni post hoc | organ x group  group  MLNs  BG-vehicle vs. BG-DTx:  BG-vehicle vs. BG-DTx+probiotics:  BG-DTx vs. BG-DTx+probiotics:  Spleen  BG-vehicle vs. BG-DTx:  BG-vehicle vs. BG-DTx+probiotics:  BG-DTx vs. BG-DTx+probiotics:  BM  BG-vehicle vs. BG-DTx:  BG-vehicle vs. BG-DTx+probiotics:  BG-DTx vs. BG-DTx+probiotics: | 0.005  < 0.001  0.0004  0.9286  <0.0001  0.0097  0.0032  >0.9999  0.9329  >0.9999  0.9682 | *  *  *  ns  ns  ns |
| S6D | CD4 T cells | BG-vehicle: n of mice=5  BG-DTx: n of mice=4  BG-DTx+probiotics: n of mice=6 | two-way mixed RM-ANOVA | F(4,24)=9.569  F(2,12)=18.282 | Bonferroni post hoc | organ x group  group  MLNs  BG-vehicle vs. BG-DTx:  BG-vehicle vs. BG-DTx+probiotics:  BG-DTx vs. BG-DTx+probiotics:  Spleen  BG-vehicle vs. BG-DTx:  BG-vehicle vs. BG-DTx+probiotics:  BG-DTx vs. BG-DTx+probiotics:  BM  BG-vehicle vs. BG-DTx:  BG-vehicle vs. BG-DTx+probiotics:  BG-DTx vs. BG-DTx+probiotics: | <0.001  < 0.001  <0.0001  0.0373  <0.0001  0.0106  0.1091  0.426  0.677  >0.9999  0.6624 | *  *  *  ns  ns  ns |
| S6E | CD8 T cells | BG-vehicle: n of mice=5  BG-DTx: n of mice=4  BG-DTx+probiotics: n of mice=6 | two-way mixed RM-ANOVA | F(4,24)=1.711  F(2,12)=6.957 | Bonferroni post hoc | organ x group  group  MLNs  BG-vehicle vs. BG-DTx:  BG-vehicle vs. BG-DTx+probiotics:  BG-DTx vs. BG-DTx+probiotics:  Spleen  BG-vehicle vs. BG-DTx:  BG-vehicle vs. BG-DTx+probiotics:  BG-DTx vs. BG-DTx+probiotics:  BM  BG-vehicle vs. BG-DTx:  BG-vehicle vs. BG-DTx+probiotics:  BG-DTx vs. BG-DTx+probiotics: | 0.180  0.001  0.3772  0.3911  >0.9999  0.1705  0.0014  0.3759  >0.9999  >0.9999  >0.9999 | ns  ns  ns  ns  ns  ns |
| S6F | Ly6C^high^ monocytes | BG-vehicle: n of mice=5  BG-DTx: n of mice=4  BG-DTx+probiotics: n of mice=6 | two-way mixed RM-ANOVA | F(4,24)=25.667  F(2,12)=20.623 | Bonferroni post hoc | organ x group  group  MLNs  BG-vehicle vs. BG-DTx:  BG-vehicle vs. BG-DTx+probiotics:  BG-DTx vs. BG-DTx+probiotics:  Spleen  BG-vehicle vs. BG-DTx:  BG-vehicle vs. BG-DTx+probiotics:  BG-DTx vs. BG-DTx+probiotics:  BM  BG-vehicle vs. BG-DTx:  BG-vehicle vs. BG-DTx+probiotics:  BG-DTx vs. BG-DTx+probiotics: | <0.001  < 0.001  >0.9999  >0.9999  >0.9999  0.0422  >0.9999  0.054  0.0003  <0.0001  <0.0001 | ns  ns  *  ns  *  * |
| S6G | Macrophages | BG-vehicle: n of mice=5  BG-DTx: n of mice=4  BG-DTx+probiotics: n of mice=6 | two-way mixed RM-ANOVA | F(4,24)=8.6  F(2,12)=12.254 | Bonferroni post hoc | organ x group  group  MLNs  BG-vehicle vs. BG-DTx:  BG-vehicle vs. BG-DTx+probiotics:  BG-DTx vs. BG-DTx+probiotics:  Spleen  BG-vehicle vs. BG-DTx:  BG-vehicle vs. BG-DTx+probiotics:  BG-DTx vs. BG-DTx+probiotics:  BM  BG-vehicle vs. BG-DTx:  BG-vehicle vs. BG-DTx+probiotics:  BG-DTx vs. BG-DTx+probiotics: | <0.001  0.001  >0.9999  >0.9999  >0.9999  0.7215  >0.9999  0.3774  <0.0001  0.5247  <0.0001 | ns  ns  ns  ns  *  * |
